# Prediction of regional wildfire activity with a probabilistic Bayesian framework

**DOI:** 10.1101/2020.05.20.105767

**Authors:** F Pimont, H Fargeon, T Opitz, J Ruffault, R Barbero, N Martin StPaul, E Rigolot, M Rivière, JL Dupuy

## Abstract

- Modelling wildfire activity is crucial for informing science-based risk management and understanding fire-prone ecosystem functioning worldwide. Models also help to disentangle the relative roles of different factors, to understand wildfire predictability or to provide insights into specific events.
- Here, we develop a two-component Bayesian hierarchically-structured probabilistic model of daily fire activity, which are modelled as the outcome of a marked point process in which individual fires are the points (occurrence component) and the fire sizes are the marks (size component). The space-time Poisson model for occurrence is adjusted to gridded fire counts using the integrated nested Laplace approximation (INLA) combined with the Stochastic Partial Differential Equation (SPDE) approach. The size model is based on piecewise-estimated Pareto and Generalized-Pareto distributions, also adjusted with INLA. The Fire Weather Index (FWI) and Forest Area are the main explanatory variables. Seasonal and spatial residuals as well as a post-2003 effect are included to improve the consistency of the relationship between climate and fire occurrence, in accordance with parsimonious criteria.
- A set of 1000 simulations of the posterior model of fire activity is evaluated at various temporal and spatial scales in Mediterranean France. The number of escaped fires (≥1ha) across the region can be coarsely reproduced at the daily scale, and is more accurately predicted on a weekly basis or longer. The regional weekly total number of larger fires (10 to 100 ha) can be predicted as well, but the accuracy decays with size, as the model uncertainty increases with event rareness. Local predictions of fire numbers or burnt areas likewise require a longer aggregation period to maintain model accuracy.
- Regarding the year 2003 -which was characterized by an extreme burnt area in France associated with a heat wave-, the estimation of the number of escaped fires was consistent with observations, but the model systematically underrepresents larger fires and burnt areas, which suggests that the FWI does not consistently rate the danger of large fire occurrence during heat waves.
- Our study sheds new light on the stochastic processes underlying fire hazard, and is promising for predicting and projecting future fire hazard in the context of climate change.

## 1. INTRODUCTION

Wildfires contribute to shape ecosystems across parts of the world and threaten human lives and properties, raising the risk issue. Mapping features of fire regimes such as frequency, size, intensity, severity or pattern of fires, across time and space is useful for planning fire and natural resource management, assessing risk and evaluating ecological conditions (Morgan et al. 2001). Indeed, fire regimes vary substantially over space and time at multiple scales, depending on many factors related to weather, climate, vegetation, orography, as well as local and regional human influences (e.g. Bradstock 2010, Bowman et al. 2011, Parks et al. 2012). Hence, on the one hand, understanding fire regimes and their economic, social and ecological consequences is a major challenge for scientists, especially in the context of climate change, which is expected to increase fire activity in many regions of the world (e.g. Flannigan et al. 2009, Barbero et al. 2015a, Turco et al. 2018, Dupuy et al. 2020). On the other hand, studying ecosystem disturbances, including fires, provide unique insights into ecological patterns and processes (Turner 2010).

Fire regimes are strongly influenced by contemporary fire management, which often aims at reducing fire activity. In some areas (US), burnt areas increased substantially despite large suppression expenditures that likely had positive feedbacks through fuel accumulation, suggesting the need to reexamine policies (Stephens and Ruth 2005, Calkin et al 2015). In contrast, fire suppression policies have likely been effective for reducing burnt areas in many regions of the Mediterranean basin (Turco et al 2016), but the long-term adequacy of such policies in the context of climate warming and fuel build-up is currently debated (Moreira et al 2020). In this context, the design and application of new policies need reinforced land management and planning, while fire suppression must continue to play a key role in the protection of human lives and assets. For planning purposes, managers and policy makers need to anticipate future scenarios-based fire regimes, while for preparedness and response actions, fire managers need to be informed on a daily or weekly basis of the expected number, size, duration and spread rate of fires (Taylor et al. 2013; Xi et al. 2019).

While wildfire regimes depend on multiscale interactions between climate, vegetation and humans (Moritz et al. 2005), weather has long been recognized as the main factor driving regional fire activity from daily to seasonal scale (Abatzoglou and Kolden 2013, Barbero et al. 2015b, Turco et al. 2017). Much effort has been dedicated to developing and evaluating weather-based fire danger rating systems, including the widely used Canadian Fire Weather Index (FWI, Van Wagner 1987), the Australian McArthur index (FFDI, Noble et al. 1980) or the American National Fire Danger Ratings System (NFDRS, Deeming et al. 1978). These indices operate at a daily time scale and are easily derived from local weather variables to inform managers, or can be projected under climatic scenarios to anticipate the effect of climate change (Dupuy et al. 2020). However, the link between fire danger rating systems and observed fire activity is not straightforward. Indeed, fire events are fairly rare at local and daily scales, and hence, highly random in nature. To handle this stochasticity, observations are often aggregated over time and prior to examining empirical relationships between fire activity and average indices, typically from weekly to monthly bases (e.g. Krawchuck et al. 2009, Barbero et al. 2014, Turco et al. 2018). Unfortunately, these correlative approaches cannot appropriately account for a number of operational and research applications that require daily predictions on fine scales. Indeed, climate, land cover and human variables can vary substantially over short distances in some regions (Fréjaville and Curt 2015). Some climate processes that occur on weekly to sub-daily scale, such as wind or hot temperature events can influence fire activity. This is typically the case in the Mediterranean region, where most fires spread during less than a day within less than 10 km^2^, even if a few fires can spread both longer and farther (Tedim et al. 2018).

The rareness and the stochastic nature of individual fire events can be addressed in a formalized probabilistic framework (Brillinger et al. 2003, Preisler et al. 2004, Preisler and Westerling 2007, Turner 2009, Vilar et al. 2010, Woolford et al. 2014, Serra et al. 2014a&b). In this approach, observed patterns of fire occurrences are viewed as realizations of a spatio-temporal point process where points correspond to locations and times of ignition of a fire, and the burnt area is used as a mark for the points. The latent (*i.e.*, unobserved) spatio-temporal intensity function that has generated the observed point pattern is then estimated. In practice, this point process is often approximated by a Bernoulli probability of fire presence in discrete and relatively small space-time cells (called voxels, typically some km^2^ X days) for which usually at most one fire has been observed. The notion of intensity (*i.e.*, expected counts) is crucial since it provides more information than only susceptibility (*i.e.*, presence-absence); in particular, intensities can be additively aggregated within any spatio-temporal mapping unit. Such fire occurrence modelling can be combined with fire size distribution models, typically expressed as the probability for a fire to exceed a given size, to simulate fire hazard (Preisler et al. 2004, 2011). These probabilistic models have most commonly been adjusted using generalized linear model (GLM), or related extensions such as generalized additive models (GAMs, Wood et al. 2006), that have been shown to perform better (Woolford et al. 2011). GAMs account for the non-linear effect of explanatory variables (such as fire danger and/or human activity metrics) and can include model components to account for spatial residuals (Preisler et al. 2004). More recently, Bayesian methods have also been used as an alternative to GAMs (Serra et al. 2014a&b, Joseph et al. 2019). Such models have been used for a variety of applications, including the issue of large fire forecasts (Preisler et al. 2008), the projection of future fire activity (Ager et al. 2018), the estimation of suppression costs (Preisler et al. 2011), or the estimation of extreme fire size (Joseph et al. 2019).

Despite their potential for wildfire predictions, probabilistic approaches still present some challenges and limitations, some of whom have not been fully addressed. First, unaggregated datasets are large, such that computational costs may be expensive and some approximations should often be implemented to run algorithms (e.g. subsampling of non-fire voxels, in Brillinger et al. 2003), thereby decreasing model robustness (Woolford et al. 2011). Second, the evaluation of the underlying model performance in a probabilistic framework is not straightforward. Indeed, it requires checking the goodness-of-fit and model parsimony in the model-building framework through various approaches including information criteria, comparisons to observations aggregated on various time and spatial scales and external validation using hold-out data (Xi et al. 2019). Third, early probabilistic approaches combined models of occurrence and exceedance of fire size above high fixed thresholds, but they did not simulate the size of fire events. Notable exceptions are Westerling et al. 2011 and Ager et al. (2014, 2018), which fitted generalized Pareto distributions with parameters depending on explanatory variables to simulate the size of individual fires. Fourth, the use of the probabilistic frameworks is still mostly limited geographically to North America, with only few applications in small Mediterranean regions (near Madrid, Spain, in Vilar et al. 2010; in Catalonia, Spain, in Serra et al. 2014a&b; or in Sardinia, Italy, in Ager et al. 2014). Finally, probabilistic approaches have seldom been used to evaluate the potential predictability of fire activity across a range of spatial and temporal scales. Indeed, it is expected that fire activity is less predictable at short time and/or fine spatial scales and for rare events (large fires), than more frequent events (small fires) over longer and/or broader scales. Probabilistic approaches provide a convenient framework to explore how this spatio-temporal gradient shapes the predictability of fire regimes. Indeed, it is critical to disentangle which fire components can be accurately predicted from what remains fundamentally, in contrast to fundamentally random behavior, to inform managers. Random behavior can be incorporated into models through stochastic components that aid in accurately assessing uncertainty in predictions, but such components do not contribute to improve predictions of expected occurrence numbers and sizes of fires. Moreover, probabilistic models help understand the extent to which catastrophic events are (un)expected, and can therefore provide useful information regarding their likelihood of occurrence, such as return periods and levels.

The objective of the present study was to understand and assess the predictability of fire activity at various temporal and spatial scales in the French Mediterranean region, through a Bayesian probabilistic approach. To this aim, we present and use a full framework of fire activity modelling, which simulates potential scenarios of daily fires occurring in small pixels (8 × 8 km). We then assess the overall model performance, the relative importance of selected explanatory variables, and the predictability at scales ranging from the pixel to the region and from daily to periods of multiple years. The assessment includes a specific focus on the catastrophic 2003 year characterized by a severe synoptic-scale heat wave in summer following a prolonged drought (Trigo et al. 2005). We finally discuss the strength and weaknesses of the current model and its potential applications for wildfire-related research avenues and the improvement of operational fire suppression and management.

## 2. METHODS

### 2.1. Data and site description

#### Study site and fire activity

The study area consists of 15 NUTS3-level French administrative units located in southeastern France (Fig. 1A, 75,560 km^2^), which concentrate the vast majority of burnt area during the summer season in France. The climate of this area is mostly Mediterranean, characterized by cool and moist winters and hot and dry summers, but exhibits strong variations with orography, from the high mountains in the Alps to coastal plains. The population is mostly concentrated near the Mediterranean coast and the Rhône river valley. These climatic and socio-economic contrasts lead to strong variations in fire activity over time and space. Fire activity is not homogeneously distributed, mostly located near the coast and in Corsica, where human activities are concentrated, and where drought and wind more pronounced (Fig 1C). Burnt area shows a bimodal seasonal pattern, with a first peak in spring associated with agricultural, pastoral and forestry practices, during which fires are generally not a major threat, and a more important second peak during the summer dry season, during which most large fires occur (Fig 1B). At the interannual scale, fire activity is highly variable (Fig 1D) and mostly dictated by annual drought conditions (Ruffault et al. 2016, Barbero et al. 2019). The outstanding burnt area of the 2003 summer was due to several extreme fires that occurred during the severe European heatwave (Trigo et al. 2005, Ruffault et al. 2018a). Following these 2003 extreme fires, fire prevention and fighting was enhanced with a modernization of the fire management law in 2004. This might explain the decrease in the number of escaped fires (larger than 1 ha) and in burnt areas after 2003 (Fig. 1D, Curt et al. 2018).

**FIG. 1.**
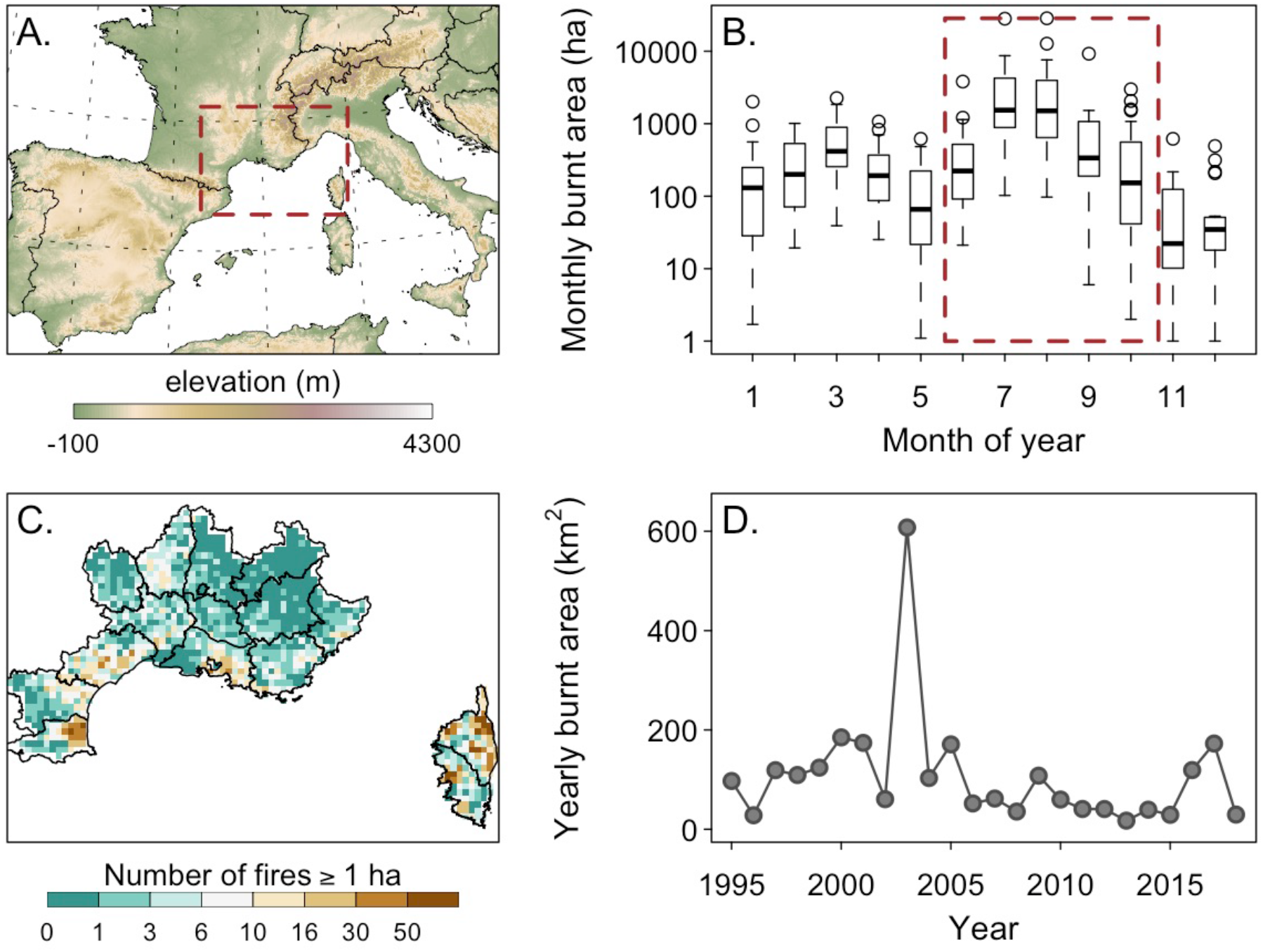
Data site and description: A. Orography (elevation in m) of the study region; Fire activity (1995-2018): B. Monthly distribution of burnt areas; C. Spatial distribution of fires larger or equal to 1 ha; D. Yearly burnt areas. Data were extracted from the Prométhée fire database (http://www.promethee.com/).

Fire records were extracted from the Prométhée fire database (http://www.promethee.com/) for the period from 1995 to 2018. This period was selected so that the dataset was large enough to allow the fitting of robust models. We discarded, however, the pre-1995 period, because of the lack of consistency of the weather data prior 1995 (monitoring stations had evolved until 1995) and of reliability and completeness issues in earlier fire records. Similarly, to limit the uncertainties associated with small fires in fire databases (Turco et al. 2013, Ruffault and Mouillot 2015), only fires larger (or equal) than 1 ha (or escaped fires) were retained. One should note, however, that the increasing precision of size records over time has led to a temporal decline of the proportion of fires exactly equal to 1ha among small fires.

We focused our analysis on the summer season (weeks 22-44, 25^th^ may to 31^th^ October, Fig. 1B), because most burned areas occur during summer, and because the causes and the factors behind spring fires are quite different and would have blurred the fire-climate relationships we wanted to explore.

#### Explanatory variables

The main explanatory variable was the daily Fire Weather Index (FWI), which represents temporal and spatial variations in fire danger. It was computed onto an 8 km-resolution grid from 12:00 LST meteorological variables (cumulated precipitation, mean wind speed, mean temperature and minimum relative humidity, calculated using specific humidity and maximum temperature) following Bedia et al. (2014), using the ‘cffdrs’ R package (Wang et al. 2017). These variables were extracted from the SAFRAN reanalysis, which is consistent in terms of observation sources after 1995 (Vidal et al. 2010).

The second explanatory variable was the forest area in each 8-km pixel, which is expected to affect both the number and size of fires, and shows significant spatial variability (Supplementary 1, map of forest area). Forest area was obtained from the CORINE land-cover database (CLC, https://land.copernicus.eu/pan-european/corine-land-cover), by merging the patch areas covered by sublevels “Forests” and “Scrub and/or herbaceous vegetation association” in each pixel. This forest area (FA, in ha or in % cover of the pixel) was estimated on a yearly basis by linear interpolation of CLC inventories available in 1990, 2000, 2006, 2012 and 2018.

### 2.2. Probabilistic model of fire activity

#### Model overview

The probabilistic model of fire activity consisted of two hierarchically structured components: one describing the occurrence of escaped fires, and another describing the size of each fire event conditional to its occurrence (Fig. 2). For the occurrence component, the response variable was the daily number of escaped fires (i.e. fires larger than 1 ha), for each pixel of the FWI grid. For the size component, the response was a continuous positive quantity (size of each escaped fire event) modelled with a piecewise distribution.

**FIG. 2.**
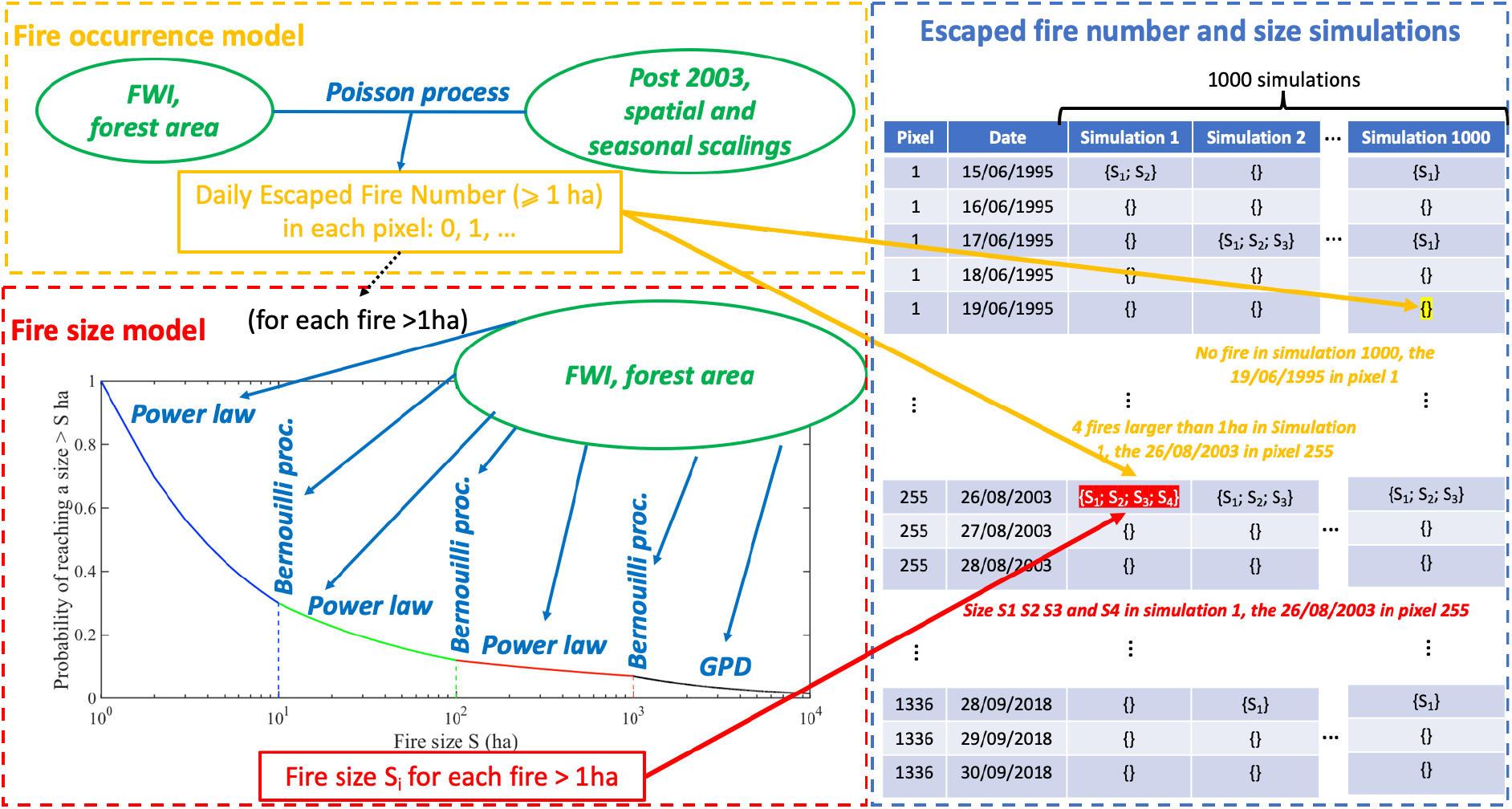
Framework of the “full” probabilistic model of fire activity

Both the occurrence and size components included FWI and forest area as explanatory variables. The occurrence model also included two temporal factors and a spatial model. Models were fitted in a Bayesian framework, using the integrated nested Laplace approximation (INLA) implemented in R software (www.r-inla.org) and described in (Rue et al. 2009, Lindgren and Rue 2015), which enables to apply the Bayesian approach to large datasets using sophisticated hierarchical structure and provides accurate and (relatively) fast inference by means of analytical approximations of the posterior model, which is in contrast to standard, simulation-based Bayesian approaches (Markov-Chain Monte-Carlo). It allows non-linear responses to explanatory variables to be estimated through flexible Gaussian prior distributions for spline functions in combination with spatial models.

Model components were trained with data from 1995-2014 (training sample), the years 2015-2018 (validation sample) being withheld for the evaluation of the predictive performance. The dataset for fire occurrence contained fire counts (≥1 ha) for approximately 4.44 million pixels/days, whereas the dataset of observed fire sizes contained 7193 fires (≥1 ha).

In the next two subsections, we describe the “full” model that includes all explanatory variables (Table 1). To verify the added value of the “full” model and to avoid overfitting (*i.e.* the situation where prediction performance on validation data decreases) (Xi et al. 2019), intermediate models for fire occurrence and size with less explanatory variables were also estimated, and their corresponding information criteria were compared with those of the “full” model.

**Table 1.**
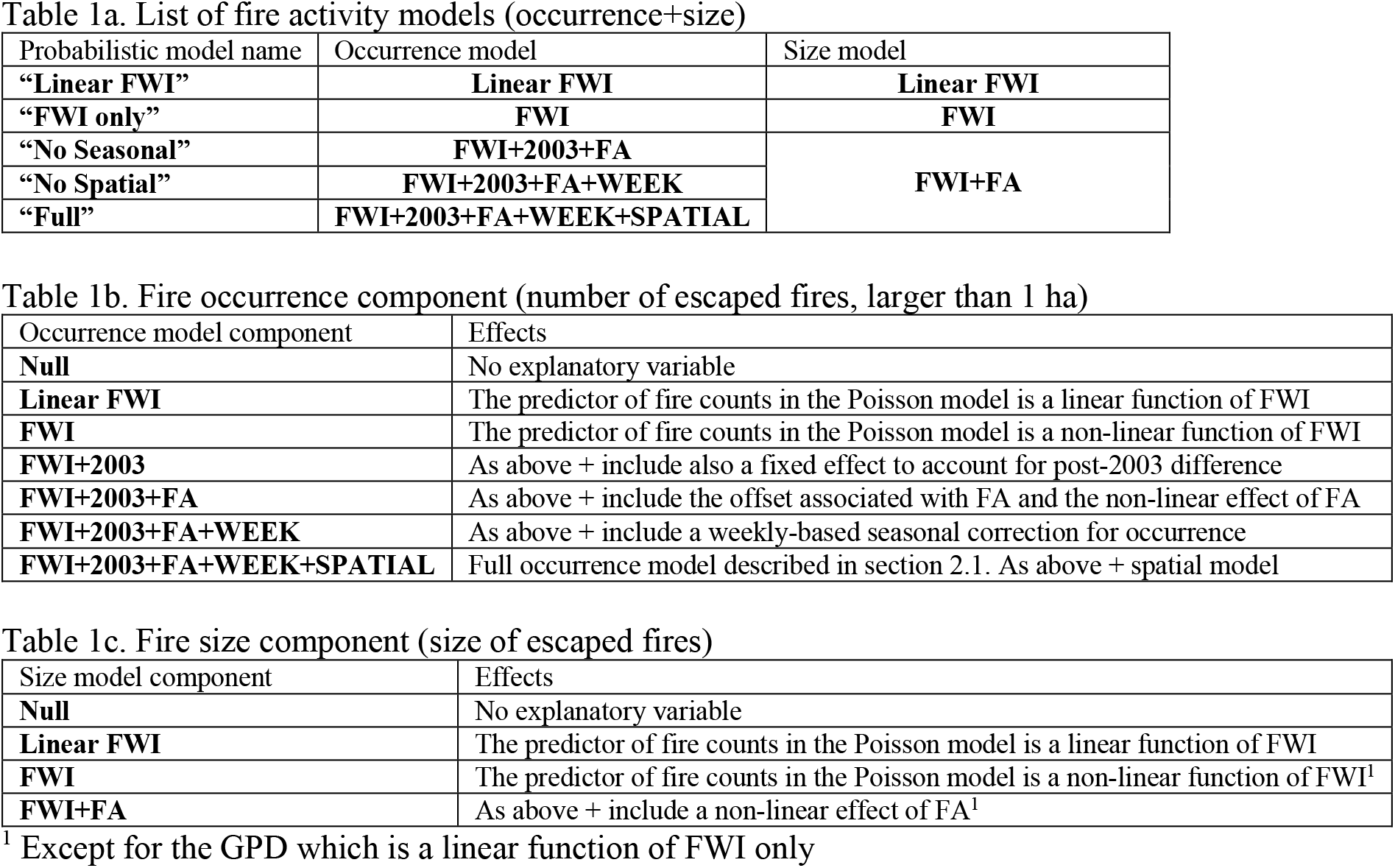
Probabilistic models developed in the present study

Combining both components of the models enable simulation, according to the estimated posterior model, of an unlimited number of scenarios (here: 1000) of the potential daily fire activity in each pixel, in the form of escaped fire lists whose number of elements and size of each were simulated with each model component. These simulations can then be aggregated at different spatial and temporal scales for corresponding predictions and evaluations against observations.

#### Fire occurrence component

We built the fire occurrence model for escaped fire (≥1 ha) counts in 8 km X 8 km daily voxels, as a Poisson random variable (Fig. 2). Following the approach of Brillinger et al. (2003), we incorporated residual spatial and temporal random effects, to account for unknown sources of variations in escaped fire probability. They can be viewed as spatial and temporal scaling factors between FWI and the observed number of escaped fires.

The voxel size was considered as a good approximation for the “true” Poisson distribution resulting from grouping intra pixel variations, since pixel-per-day probabilities remained small (Brillinger et al. 2003). Contrary to large voxels in which multiple fires can occur more often (e.g. Joseph et al. 2019), our Poisson-based method was applied to an almost binary dataset, and spatial correlations were accounted for with a spatial model, so that over-dispersion was less of a concern (Taylor et al. 2013). For the same reasons, the use of a zero-inflated Poisson model was not required. Moreover, this resolution was fine enough to explicitly link the fire occurrence probability to locally observed fire conditions (weather data and forest area), rather than to some average value at a coarser scale (Taylor et al. 2013). To appropriately identify the range of variation of spatial biases not explained by the available predictor variables (FWI, FA, season), the pixel size should be by a multiple smaller than the distance at which the correlation in the spatial component of the model drops to near zero, e.g. drops below 0.1. This avoids issues related to within-pixel overdispersion and to wrongly reporting very smooth spatial maps of occurrence intensity. Model fit showed that this range was approximately 30 km, which was indeed substantially larger than pixel size. Finally, this size was consistent with the computational and memory costs of INLA, which strongly increase with the size of the dataset and the resolution of spatial and temporal random effects.

The partial effects of the models were assumed to be multiplicative, based on additive decomposition of the log of expected fire counts, which has been shown to be adequate for time and space in Woolford et al. (2011). The form of the “full” model, which included all explanatory variables (“FWI+2003+FA+WEEK+SPATIAL”, see Table 1b) was:

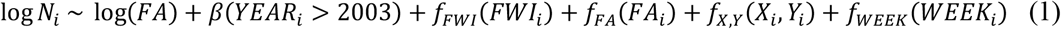

where log(*FA*) was a deterministic fixed offset, *β* the fixed effect (with a different value before and after 2003), and the f-terms captured nonlinear influences of the covariates FWI and FA and spatio-temporal effects.

Because escaped fires cannot occur in non-forested area (urban areas, crops, etc.), the area of each pixel in which fire “points” could happen was not spatially constant. This variability was incorporated in the Poisson model with an offset equal to forest area (FA). The model allowed non-linear effects of FWI, but also of FA – in addition to the offset – as a land use factor. Indeed, it is expected that the probability to get a fire per area of forest decreased for high forest area, since interface, road and urban densities decreased. Spatial effects were represented using the Stochastic Partial Differential Equation approach (SPDE, Lindgren et al. 2011), which estimates the spatial model for residuals, through continuous spatial random effects at high resolution. Temporal effects were incorporated as a non-linear weekly seasonal factor and a fixed effect “post 2003”. This “post 2003” effect should not be interpreted as an actual shift if the relationship that occurred exactly in 2003, but more as a convenient and simple manner to incorporate in the model the temporal evolution of the fire-weather relationship. In this study, we decided to ignore other annual effects in order to develop a model applicable to predictions and projections. Contrary to other studies (e.g. Opitz et al. 2020), the focus of the present model was not to model time variation in spatial patterns.

The prior distributions of the different predictors were Gaussian processes. The nonlinear f-functions in (1) were modelled with piecewise constant first-order random walks, with 30, 18 and 23 segments, for FWI, FA and the seasonal effect, respectively. For each of them, one hyperparameter (called *precision*) governed curve smoothness (i.e., the size of the small steps between consecutive segments), as a Bayesian variant of smoothers used in GAM models (e.g. Preisler et al. 2004). For the spatial component, the SPDE approach consists in implementing a numerically convenient approximation to the Matérn covariance function for the Gaussian random field prior of *f_X,Y_* in the 1143 pixels (meshing the study site). Two hyperparameters were estimated for this random field: precision (to control the spatial variability of field values) and range (to control spatial dependence, *i.e.* the smoothness of the spatial surface). For hyperparameters, we specified Penalized Complexity priors (Bakka et al. 2018), which penalized the distance of a model component towards a simpler baseline specification (*i.e.*, absence of an effect such that its contribution to the log-intensity is 0), and we fixed penalty parameters that ensured relatively smooth estimated posterior effects. In order to limit computational and memory costs, we took advantage of the additivity of the Poisson process to aggregate data in segment classes to reduce dataset size, which initially contained 4.44 million voxels. The numerical design described above enabled keeping the number of observed classes below 500,000, which avoids numerical instabilities when running R-INLA, and models are estimated within several minutes to several hours in case of the full model.

#### Fire size component

We built a probabilistic model for sizes of escaped fires (conditional on a fire being larger than 1 ha), which corresponded to the marks of the “fire” point-process (Fig. 2). Because the fire size distribution is usually not well reproduced by any of the commonly used probability distributions over the whole range of observations -in particular for small and large fires- (Cui and Perera 2008), we used a piecewise specification of the distribution based on Pareto and Generalized Pareto Distributions in the different size segments, as justified by the asymptotic theory of threshold exceedances (Davison and Huser 2015). In each segment, size distributions depended on both FWI and FA of the voxel (8 km X 8 km by daily cell) in which the fire initially spread. In principle, we could have estimated the probability of a given fire to exceed the upper threshold of a segment by using the exceedance probability derived from the fire size distribution within this segment. However, because of the small fraction of fire sizes in the higher parts of each segment (*i.e.*, most fires have size closer to the lower than the upper bound of each segment), we obtained more accurate estimates of exceedance probabilities with specific logistic regressions for each threshold. In summary, the size model was generative, such that we can simulate from it, and it had hierarchical structure using a piece-wise specification over intervals of burnt area (Fig. 2). First, a logistic-regression-based model determined the segment into which each individual fire should fall (1-10 ha, 10-100 ha, 100-1000 ha or larger than 1000 ha). Then, the exact size was simulated according to the distribution of the corresponding segment.

Contrary to other regions of the world where fires can spread over tens of km^2^ during several days (e.g. Joseph et al. 2019), most fires in the study area spread for less than a day and were smaller than 1000 ha (which was much smaller than the pixel area). Therefore, it is appropriate to stick to the voxel scale for fire size modeling, even if a very small number of fires spread over more than one voxel. The rationale for including FA (in addition to the FWI) was that a small forest area is thought to limit fire spread. Contrary to the fire occurrence model, we did not include any other spatio-temporal factors in the size model, as the dataset size was too small to develop robust models. The form of the size component of this “full” model was hence “FWI+FA” (table 1a), except for the GPD (Table 1c).

We developed a piecewise model of fire size distribution that did not make any restrictive assumption on the general shape of the probability density of the distribution. Instead, we used standard modeling techniques suggested by extreme-value theory, based on threshold exceedances. We carried out preliminary analyses of the response of fire size distributions in different FWI classes based on mean excess plots (Hall and Wellner 1981) of the log-transformed escaped fire sizes. The number of exceedances over increasingly high thresholds suggested a slow power-law-like tail decay for most of the thresholds except the highest ones, for which exceedance numbers seem to decrease much faster as in the power-law setting, similar to the findings in Cui and Perera (2008). For our data, the behavior of mean excess curves of log fire sizes, and of related curves (cumulative distributions in log-log scale), tended to change around fire sizes of 10 ha, 100 ha and 1000 ha. Therefore, we assumed that the distribution of fire sizes could be modelled through piecewise Pareto distributions between thresholds *u*_1_ = 1, *u*_2_ = 10, *u*_3_ = 100 and *u*_4_ = 1000 ha, which depended on both FWI and FA (equivalently, through piecewise exponential distributions for log fire sizes). More precisely, given a threshold *u*_*k*_, we estimated exponential regression models for 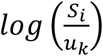, where *S_i_* corresponds to an observed fire size larger than *u*_*k*_, for segments *k*=1, 2, 3; moreover, we censored observations *S_i_* > *u*_*k*+1_, so that the model provides a good fit for *u*_*k*_ ≤ *S_i_* < *u*_*k*+1_ by construction. The estimation was conducted with INLA using its survival model framework for handling censoring, and FWI and FA were used as covariates with potentially nonlinear influence:

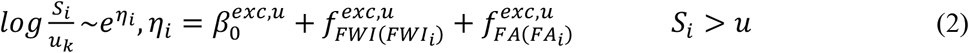

where the f-terms captured nonlinear influences of the covariates FWI and FA, with piecewise constant first-order random walks, with 20 and 10 segments, respectively.

Finally, for the category with largest fire sizes (exceeding 1000 ha and containing 33 and 7 fires for the periods 1995-2014 and 2015-2018, respectively), we took into account physical considerations and selected a generalized Pareto distribution (GPD), which allows for a finite upper endpoint if its shape parameter is negative, since an upper bound – even a very large one – must necessarily exist. The GPD has shown to perform generally well for large fire sizes (Schoenberg et al. 2003; Westerling et al. 2011). Therefore, we estimated the shape *ξ* and scale *σ* parameters of the GPD, by fitting it 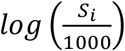 for observations *S_i_* > 1000. Since this model with the possibility for negative shape parameter was not available within INLA due to some peculiarities of its density (*e.g.*, de Haan and Ferreira 2007), we estimated the GPD parameters using frequentist maximum likelihood, followed by a careful inspection of the estimated model. Owing to the small sample size, we chose a more parsimonious parametrization of covariate influence using only linear coefficients:

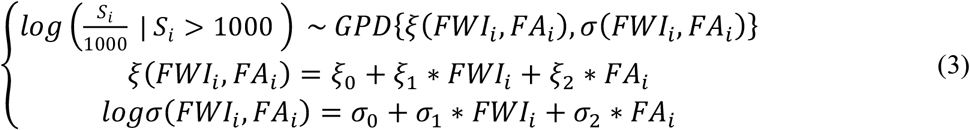

However, because the sample size was very small, uncertainty on the FA coefficient was very high and confidence intervals and information criteria advised against including FA in this model (see supplementary 2 for details). We hence selected for the “Full” model a GPD with parameters function of FWI only.

As mentioned above, we cannot expect a good estimation of exceedance probabilities derived from the three power laws due to the relatively small sample fraction of fire sizes in the higher segments. For example, a only moderately large number of fires were greater than 10 ha (1348 fires among 7193), and only few of them (280, approx. 4 % of fires larger than 1 ha) were also larger than 100 ha. Between 1995 and 2018, only 40 fires reached more than 1000 ha (0.6% of fires larger than 1 ha). Therefore, we separately modeled and estimated these exceedance probabilities 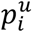, for a threshold u and a voxel i, with INLA, based on logistic regressions for the indicator variables of threshold exceedances (i.e., 1 if fire size exceeds *u_k_* and 0 otherwise), given FWI and FA:

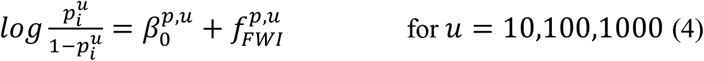

where the f-terms captured nonlinear influences of the covariates FWI and FA, with piecewise constant first-order random walks, with 20 and 10 segments, respectively.

These three probabilities and the four estimated fire size distributions were hence combined to predict the size of each fire larger than 1 ha, given FWI and FA, through a sequential approach consisting in simulating i) in which segment the size of the fire is, and, conditional to the segment, ii) the exact size (in this segment).

#### Variable selection and model evaluation

The final “full” model was developed by including the different explanatory variables and non-linear functions step by step, checking information criteria of the intermediate probabilistic models. A selection of intermediate models is presented in Tables 1a,b,c, ranging from the simple “FWI-linear” to the “full” model. We used the DIC (Deviance Information Criterion) and the WAIC (Widely-Applicable Information Criterion) for variable selection in submodels, which are generalizations of the well-known Akaike Information Criterion (AIC) for Bayesian models. The WAIC is known to reflect posterior uncertainty in the models’ prediction more fully than DIC, which can sometimes select over-fitted models (Vehtari et al 2017). For the Generalized Pareto Distribution of largest fires, we simply used the AIC of the model fit. The robustness of the occurrence model was checked in a preliminary development, thanks to a 7-fold cross validation procedure, holding out 3 randomly selected years in each fold, which demonstrated very little sensitivity to data sample (Fargeon 2019).

We evaluated the performance of the model by two different means. First, we evaluated the subcomponents of the model with Area-under-the-Curve measures (AUC, Fawcett 2006). AUCs rate the model ability to diagnose the realization in voxels of different events, such as “at least one escaped fire” and a selection of “size exceedances”. AUC values range between 0 and 1, with 1 indicating perfect prediction of the binary presence/absence information, whereas 0.5 indicates a random prediction. AUCs were computed for both the 1995-2014 (training) and 2015-2018 (validation) periods. Second, we evaluated model performance by comparing simulations with historical observations aggregated on various temporal and spatial scales (Xi et al. 2019). These evaluations were carried out from 1000 model simulations of fire occurrence per voxel, which were sampled as a Poisson process according to draws from the posterior predictive distributions of the intensity in occurrence model. Note that INLA (in contrast to Markov Chain MonteCarlo, MCMC) does not provide simulations of the posterior model’s component during the estimation process, but sampling from the fitted model is nevertheless straightforward and useful for calculating sophisticated predictions going beyond the standard output of R-INLA (e.g. Fuglstad and Beguin 2018). A fire size was then randomly assigned to each simulated escaped fire based on the size submodels (Bernoulli processes for size segment, Pareto and Generalized Pareto distributions) parametrized with posterior mean parameters. This approach allowed considering the inherent variability of the stochastic processes at stake. This variability was used to draw pointwise envelopes showing the spread between 5^th^ and 95^th^ percentiles of fire activity, and to compute expected trends for the different spatio-temporal aggregations of simulated fire activity.

The overall goodness-of-fit between expected trends and observations were measured with mean absolute error (MAE, in %). These errors were examined with respect to model uncertainty (MU, in %), which quantified the stochasticity of the corresponding trend, expressed as the variability of simulated quantities over the 1000 simulations. MU was computed as the mean absolute deviation of the simulated activities to rate the model spread, expressed in % of the observed value. The last metric used to evaluate the model was the coverage probability (CP) of the 95% confidence interval, which measured how often observations fall within the estimated confidence interval. If the model was perfect, it would be equal to 95 %. A coverage significantly different from 95% means that the model was either biased or exhibited an incorrect variability.

### 2.3. Model applications

Once the model has been evaluated, it can be used to provide understanding of the stochasticity of fire activity, given that 1000 simulations of the models can provide more insight than the single realization of observations. Two example applications were developed in the present paper.

#### Detailed analysis of year 2003

For the first application, we provided detailed comparisons of seasonal predictions and observations in 2003, during which the total burnt area was extremely high for study area. Some metrics were expected to be in the upper part or even out of confidence intervals, as it was a catastrophic summer season, even if fire danger was high because of the heat wave.

#### Predictability analysis

In order to better understand the predictability of fire activity, we compared simulated fire activity for years 2015-2018 to observations (validation sample) within a crossover plan of spatio-temporal aggregations, and for a selection of fire sizes. We used six classes for spatial aggregation (ranging from the single 8-km pixel to the whole studied area) that were crossed with seven classes for time aggregation (ranging from a single day to the four 2015-2018 years). MAE and MU, as defined in the previous section, were computed based on the 1000 simulations for the corresponding 42 aggregation classes, for fire numbers ranging between 1 and 500 ha and for total burnt area. This approach allowed to diagnose the fire sizes and aggregation scales for which simulations were in agreement with observations.

## 3. RESULTS

### 3.1. Presentation of the “Full” model

#### Partial effects of the “Full” model

The partial effects of both components (occurrence, size) of the “full” fire activity model for the different explanatory variables on escaped fire numbers and on a selection of exceedance probabilities are shown in Fig. 3. The 95^th^ credible intervals were obtained from the posterior predictive distributions. As expected, the FWI had a very strong effect on the expected number of escaped fires, which was about 60 times higher for a FWI of 60 than of 5 (Fig. 3A). This effect was very limited for FWI above 60, with wider credible intervals, due to smaller sample size for the most extreme values. As for the FWI, we observed a positive effect of forest area (FA, including both *f_FA_* and the offset, see Eq. 1) on the expected number of escaped fires with a maximum reached around 30%. The slight decrease starting at around 40 % reflected a strong decrease in escaped fire density (number of fires per unit of forest area) observed in pixels with the highest FA.

**FIG. 3.**
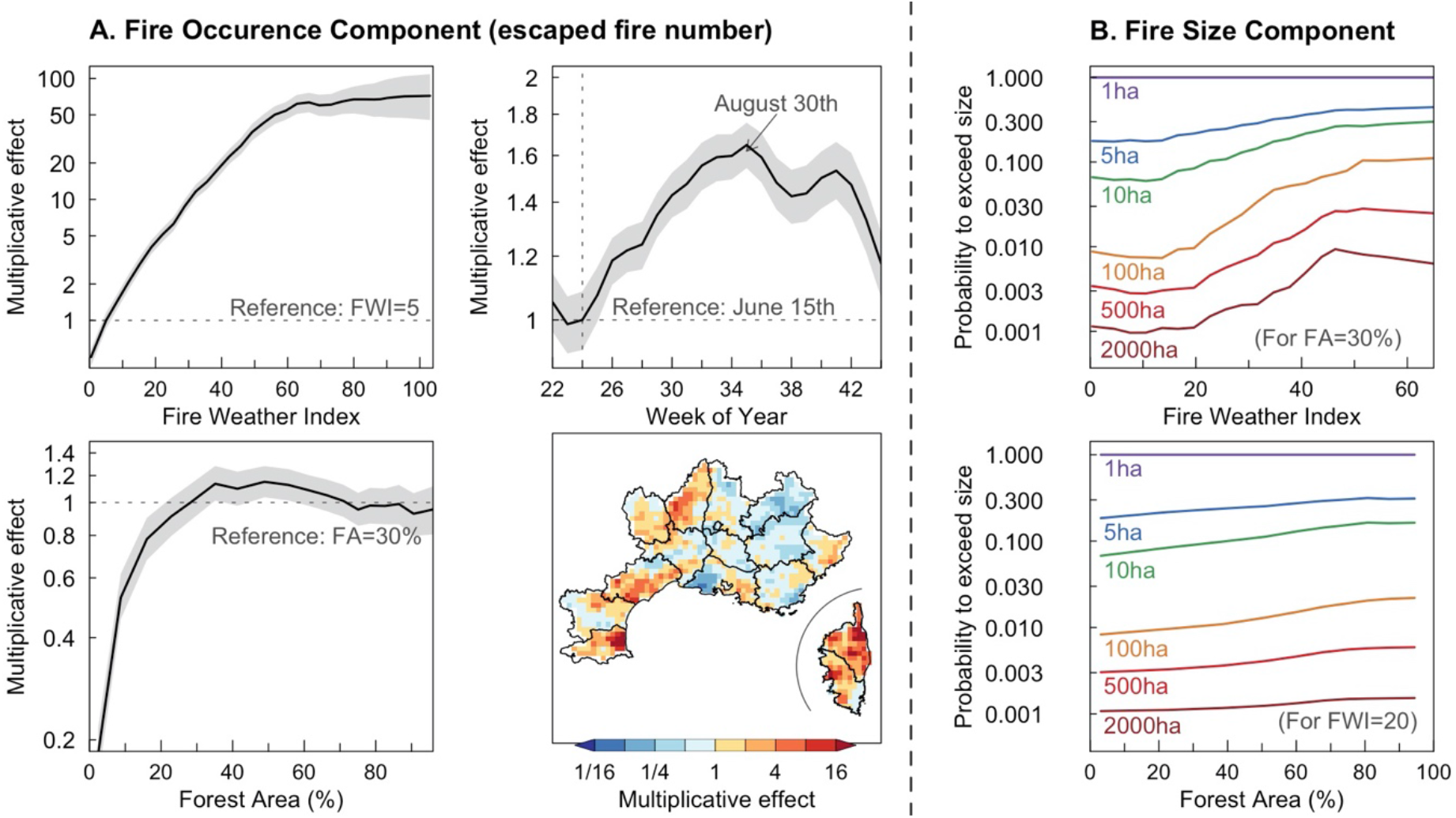
Partial effects of the “Full” fire activity model: A. Occurrence component (number of escaped fires, i.e. larger than 1 ha): effects of FWI, Forest Area (including offset), Week of Year, and location; B. Size component (exceedance probabilities of escaped fires for a selection of thresholds ranging between 1 and 2000 ha): effects of FWI and FA, for a Forest Area of 30 % (top) and a FWI of 20 (bottom), respectively.

The partial effect of the season showed a constant increase between mid-June and the end of August, and then decreased during autumn. Even if its amplitude implies only a moderate variability in relative wildfire risk, it indicated that the FWI was not fully consistent to rate escaped fire occurrence over the course of the fire season. For instance, for a same fire danger level, escaped fires were 1.6 times more numerous in late August than in mid-June. According to the estimations of the spatial SPDE model, the spatial effect was much stronger in amplitude, indicating that very different fire activities were associated with the same FWI level at different locations. The “post-2003” effect (Eq. 1), which was not shown in Fig 3A, was equal to 0.46 in posterior mean, meaning that the number of fires was less than half as small for years 2004-2014 than for years 1995-2003, confirming the important difference in fire-weather relationship between the two periods. At this stage, the relevance of this fixed effect for predictive purposes is not proved, because the difference between the two periods could not be well modelled by this crude “post-2003” effect, despite its statistical significance (with 1 not contained in its 95% credible interval). The detailed comparisons between simulations and observations conducted in the next subsections would confirm the truth of such an abrupt transition. Among these different effects, the FWI and the spatial model exhibited the strongest amplitudes.

The effects of FWI and FA on the size of escaped fires showed that the probability to exceed a given size generally increased with both explanatory variables. Moreover, these exceedance probabilities decreased when larger fire size were considered, as expected (Fig. 3B). For example, the probability to exceed 10 ha in a pixel with FA=30 % increased from 0.069 for a FWI of 7.3 to 0.325 for a FWI of 64. It is also interesting to note that the probability to exceed 500 ha was higher when FWI was high than the probability to exceed 100 ha when FWI was low. It should be noted, however, that the amplitude of these effects was generally much smaller than in the occurrence model.

Surprisingly, the exceedance probability decreased for fires larger than 2000 ha at higher FWI, but it had low significance because of the very small sample of fires larger than 2000 ha. We further point out that highest FWI values were not equally distributed in space but were most often observed in areas less prone to large fires (e.g. coastal populated areas). As shown later for the occurrence model, the use of a spatial model could have limited such a confounding effect, but the dataset was too small to afford it for the size model. More surprising was the moderate decrease observed for FWI lower than 10. In the dataset, a non-negligible number of medium and large fires was recorded for very low FWI (<5), with 60 fires larger than 10 ha occurring with FWI lower than 5. For example, three large fires (936, 2369 and 4378 ha) occurred in 2003 when FWI was lower than 1, 2 and 4, respectively. This could be explained by either uncertainties in the weather reanalysis (SAFRAN) or by daily-scale events (e.g. afternoon thunderstorms following fire events) or simply, poor rating of actual fire danger conditions by the FWI.

#### Example simulated scenarios of fire activity with the “Full” fire activity model

As an illustration of the model practical utility, fire activity simulations aggregated for the whole zone were compared to seasonal historical observations (black dots) at daily (for escaped fires only) or weekly scales (escaped fires, fire number larger than 10, 50 and 100 ha, as well as burnt areas). Results are shown in Fig. 4 for the example year 2001, but similar figures for other years of the study period are available in Supplementary 4. As expected, the uncertainty due to stochasticity (MU, in %) was larger for daily than for weekly escaped fires, and tended to increase with the fire size of interest, partly because the numbers to predict were smaller. Although not exactly equal to 95%, coverage probabilities (CP) were of the right order of magnitude, even when the width of the confidence intervals was relatively narrow (e.g. weekly escaped fire number, CP=70%). MAE were most often slightly larger than MU, which illustrated model skills, despite high stochasticity in the data.

**FIG. 4.**
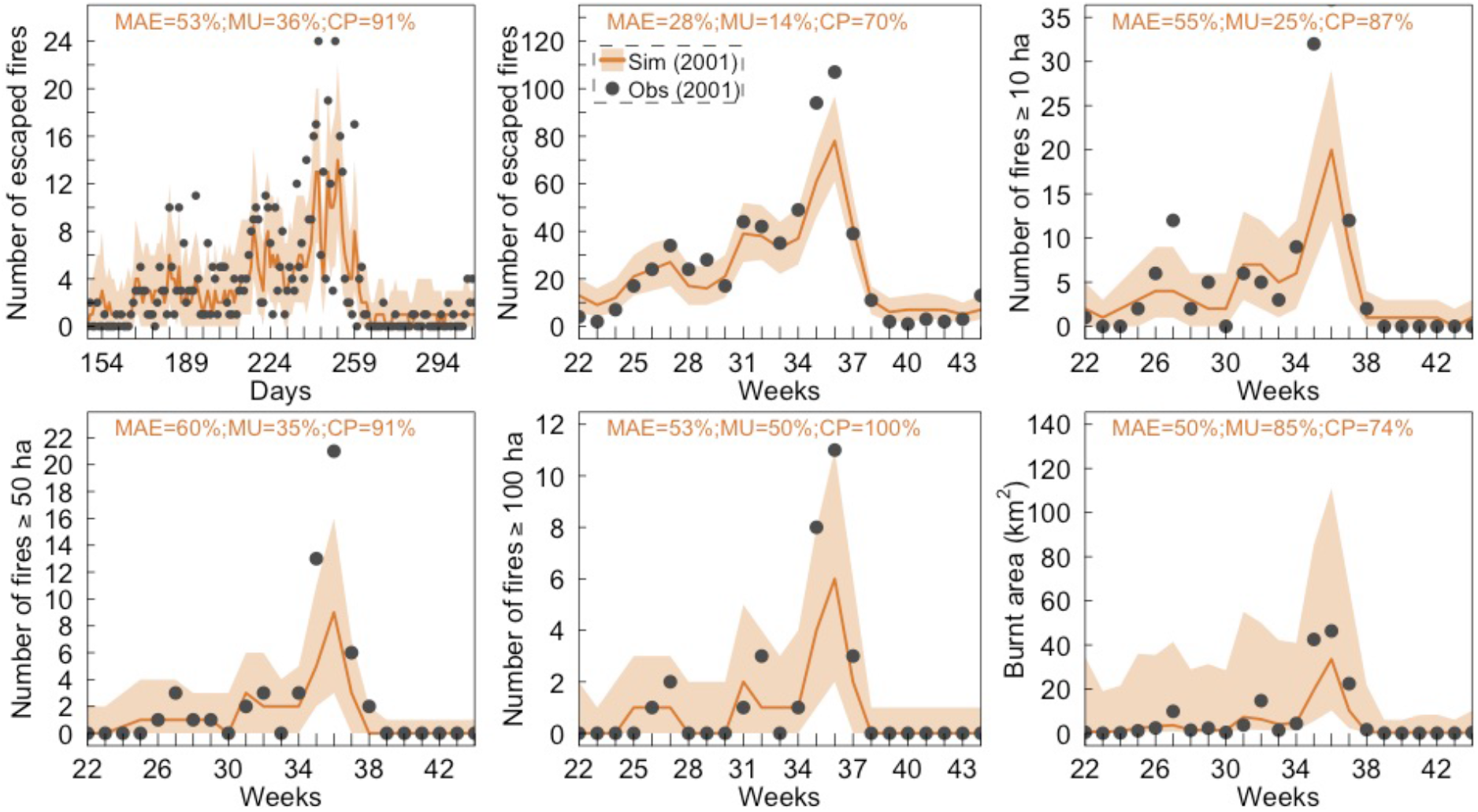
Simulated fire activity (in orange) and observations (black dots) for year 2001: daily and weekly escaped fire numbers, as well as weekly number of fires larger than 10, 50, 100 ha and weekly burnt areas added up over the whole study area. Expected trends (red line) were surrounded by the 95^th^ confidence interval in orange and were based on averages computed over 1000 simulations of fire activities for all voxels of year 2001.

Next, to study in detail the ability of our modeling framework to reproduce observed patterns of fire size distribution, simulated cumulative distributions of fire size were compared to observations for the same example year in Fig. 5. Although observations may deviate from expectations for the largest fires, most exceedance probabilities fell into the simulation-based 95^th^ confidence interval. The simulated trend for 2001 was close to the mean simulation (orange dotted line, for year 1995-2018), but this was not the case in general (e.g. years 1997 or 2002, see supplementary 5 for details).

**FIG. 5.**
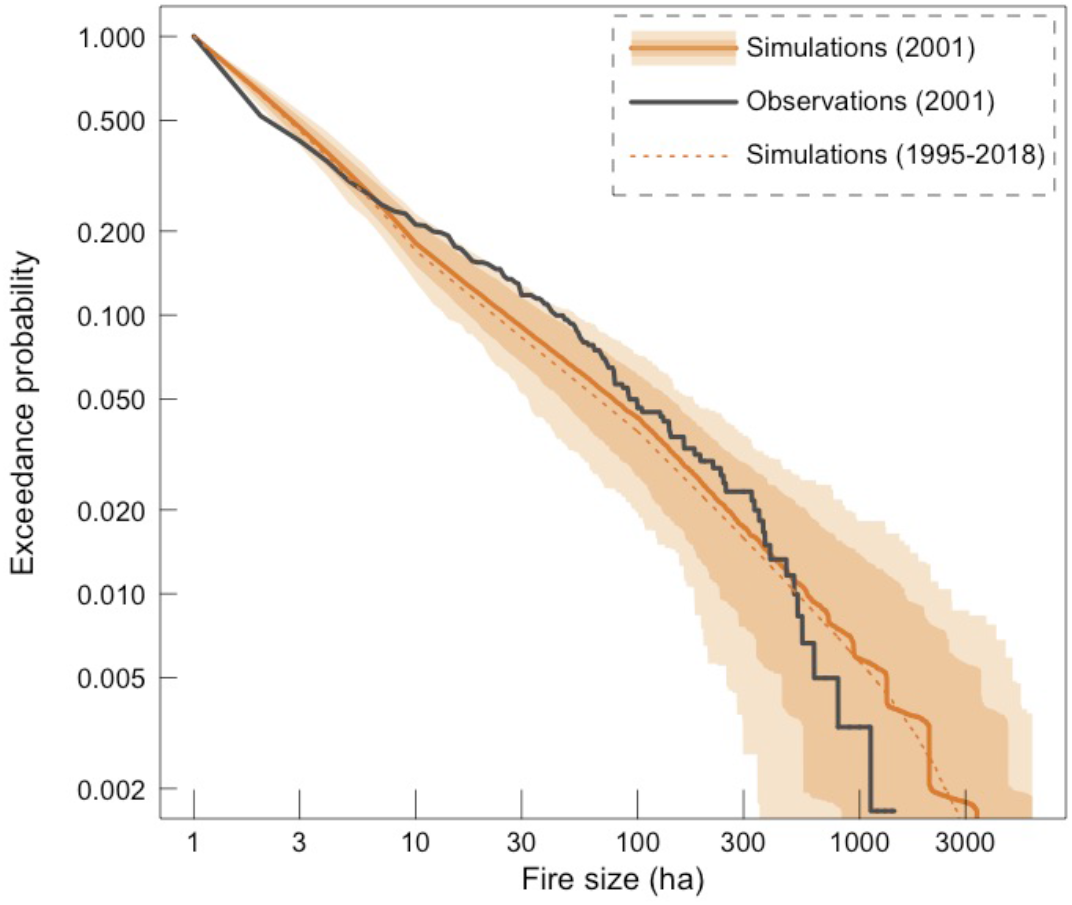
Simulated fire size cumulative distribution and observations for year 2001: Expected trend (orange line) was surrounded by the 95^th^ confidence interval in orange and the 99.9^th^ confidence interval (light orange), computed from 1000 simulations of fire sizes for year 2001. The dotted line corresponds to the fire size distribution from 1995-2018.

### 3.2. Model evaluation and importance of each explanatory variable

#### Variable selection and model fits

Model information criteria (DIC, WAIC), and AUCs for years 1995-2014 (training sample) and 2015-2018 (validation sample), are reported in several Tables in Supplementary 2. Information criteria aimed to assess goodness-of-fit of models while safeguarding against overly complex models that overfit data (Vehtari et al 2017), and were an appropriate means to check if the structure of each response to an explanatory variable, as implemented in the “full” model, was significant and parsimonious. AUCs enabled verifying that the probabilities to get at least one fire and to exceed fire size thresholds were better predicted with the selected models that included more explanatory variables and/or non-linear responses. A selection of these AUCs is presented in Fig. 6. The performance of the “full” occurrence component was high (>0.8) on both training and validation subsets, and better than the simple FWI-linear model. Regarding the size component model, the predictability of medium fire sizes (50 to 500 ha) was highest with AUC > 0.75. AUCs were in general on the same order for the validation (2015-2018) and the training sample (1995-2014).

**FIG. 6.**
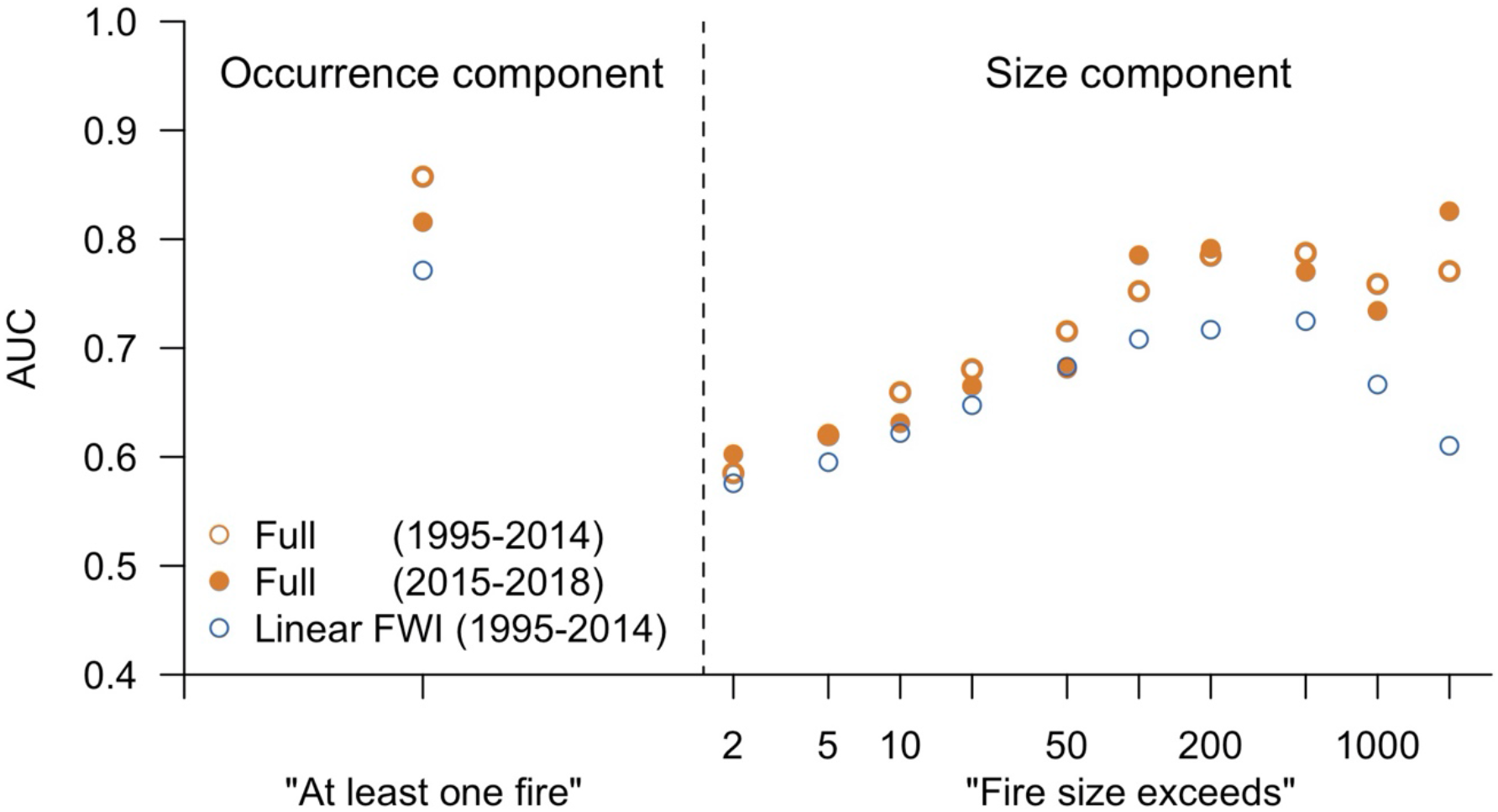
“Areas Under the Curve” (AUCs) corresponding to the realization of events diagnosed according to the two components of the fire activity model. The different series correspond to the “full” model (on the subset used to fit data, 1995-2014; and for the subset used to test the model, 2015-2018) and to a more basic “Linear FWI” model (before 2014), which only implemented the linear effect of the FWI as explanatory variable.

#### Evaluation of expected trends and spatial patterns

In order to more comprehensively evaluate the model, fire activity simulations were compared to the historical observations for spatially aggregated annual and seasonal data as well as temporally aggregated data at the pixel level in FIG. 7 (escaped fire numbers) and FIG. 8 (burnt areas). Orange lines and left maps (“Full” model) compared generally well to observations, contrary to blue lines and right maps, which correspond to different intermediate models, with less explanatory variables. This demonstrated the absence of major bias of the “Full” model (metrics in orange), as well as the limitations of intermediate models (metrics in blue).

**FIG. 7.**
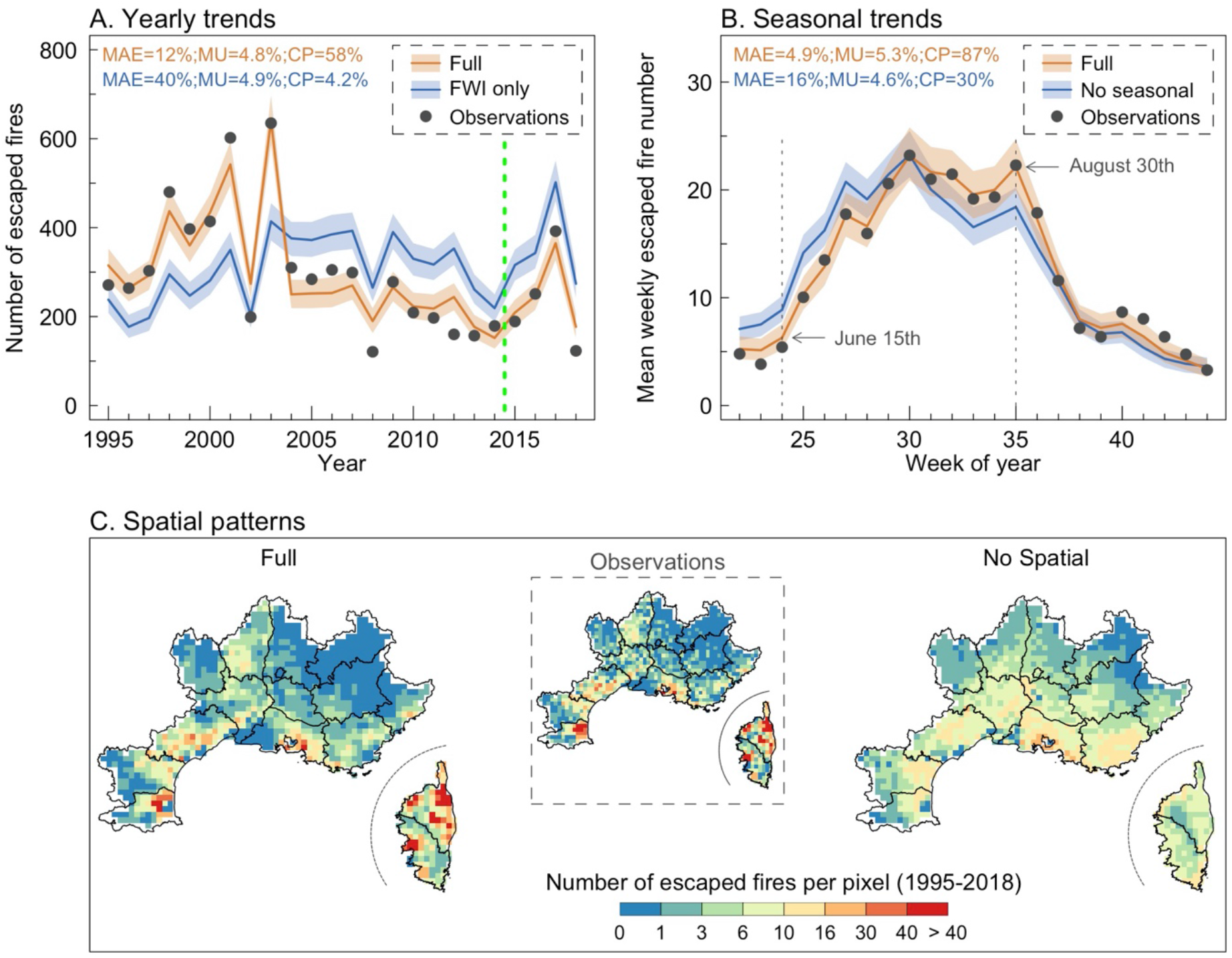
Evaluation of the occurrence model component (escaped fire number) for a selection of aggregation scales. In each subplot, the “Full” model is compared to an intermediate model where one critical effect required for consistent simulations at this aggregation scale was not included. A. Yearly trends for the whole studied area (the dashed vertical line shows the separation between training and validation sample); B. Seasonal trends at the weekly scale; C. Spatial patterns in number of escaped fires.

**FIG. 8.**
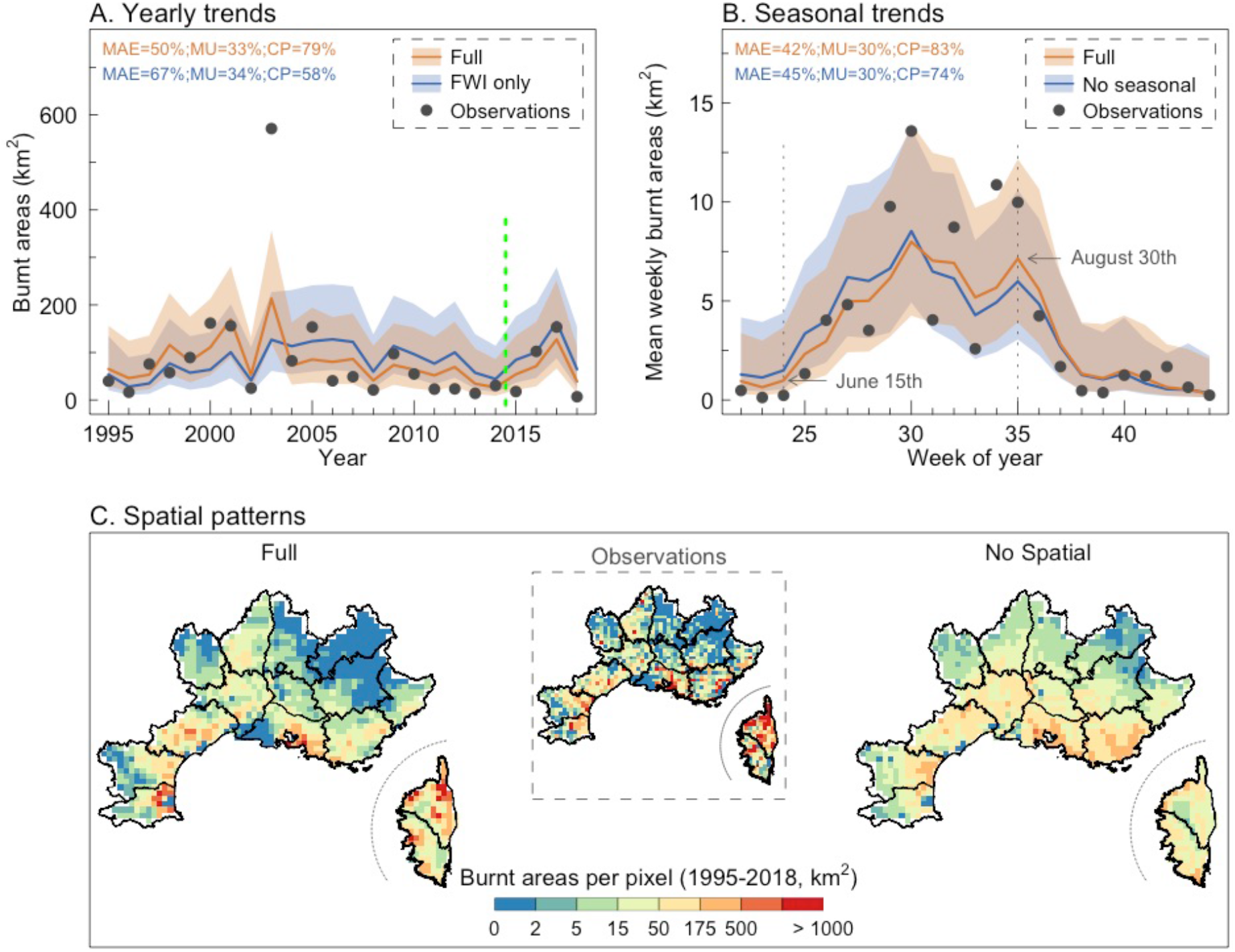
Same as Fig. 7 for burnt area.

In particular, annual trends in escaped fire number (Fig. 7A) were poorly predicted without the “Post-2003” effect, with a CP of only 4.2 % and a MAE of 40 %. The systematic overestimation of fire activity after 2003 with this model clearly indicates that the fire-weather relationship changed over time. The “Full” model, which included a “Post-2003” effect, performed much better than the intermediate one but remained still slightly biased and did not fully accounted for the evolution of the fire-climate relationship over time, or underestimated confidence intervals (CP of 58%, lower than 95%). In particular, we note that the “Full” model underestimated escaped fire numbers in years 2004-2007, suggesting that the transition was probably less abrupt in observations than assumed with a single fixed “Post-2003” effect. Several factors could explain this strong evolution near 2003. This includes the evolutions in fire management after 2003 (modernization of the fire management law, increase in airborne armed-guard funding), as well as the decrease in 1ha fire proportion in records (because of increase in size precision), which was marked near 2003. At the end, we considered the “Full” model satisfactory, because trends for recent years -and especially those simulated after 2015 (validation sample)- were in good agreement with historical observations.

Although moderate, the seasonal correction at the weekly scale enabled to match observations closely, with a CP of 87% (Fig. 7B), which was relatively close to 95 %. The trends observed with the “No seasonal” model showed that FWI explained a large part of the seasonal dynamics, but that the escaped fire number was overestimated until the end of July and underestimated in late August, which was consistent with the partial effect of the Week of Year shown in Fig. 3. When the spatial model component was not included (“No spatial”), the occurrence component did not simulate the spatial patterns of escaped fires well (Fig. 7C, right). In particular, hot spots were not well predicted, whereas the model predicted too many fires in the Alps and in the Camargue region (coastal plain in the Rhône valley).

In general, burnt areas exhibited similar results (Fig. 8), albeit with a few notable differences. First, the observed burnt area in 2003 was strongly underestimated, and was way above the upper bounds of confidence intervals (Fig. 8A). This point will be further analyzed in section 3.4. Second, the confidence interval widths, which expressed the amount of stochasticity, were much larger for burnt areas than for escaped fire numbers, with model uncertainties on the order of 35 % for both annual and seasonal predictions. Although such a high stochasticity was expected due to the very flat tail of the fire size distribution between moderately high and the most extreme values, one could argue that it was overestimated by the model. Two main clues indicated that it was not the case. Although no obvious bias was evident in temporal and spatial trends, CP were on the order of 75-80 %, which was not too far of the target (95 %), suggesting that stochasticity was on the right order of magnitude. Moreover, observed mean weekly burnt areas (averaged over 1995-2018) exhibited very large fluctuations between consecutive weeks (between weeks 28 and 36), whereas no obvious mechanisms except randomness could explain such a behavior (Fig. 8B). In particular, even if strong North winds generally occur more frequently in July than in August in this part of France, the reasons why burnt areas for the last week of July and the first week of August were extremely contrasted (13.6 km^2^ for week 30; 4.0 km^2^ for week 31) remained random.

Beyond the uncertainties, we would like to point out that the “full” model performed very decently, especially when compared to intermediate models. It was all the more remarkable as the size component only included FWI and FA as explanatory variables. Indeed, the temporal and spatial correction factors in the occurrence component were sufficient to obtain a general agreement with burnt area observations. Although 1 ha fires (escaped fires) only played a minor role in the overall burnt areas, this finding highlights their importance in the simulation of burnt areas.

Expected temporal trends and spatial patterns in simulations looked like a smoothed expression of observations, in which stochasticity would have been removed. However, one should notice a few spurious differences. Simulations seemed to slightly overestimate burnt areas at the beginning and the end of the season, and to exhibit less burnt areas than observed in the Corsica and Var NUTS3 units, where occurred most of the very large fires of the 2003 season. Interestingly, simulated burnt areas were only slightly better predicted when they were simulated from escaped fire observations (using the fire size component only, see Supplementary 3.1). This revealed that the limitation in burnt area simulations mostly arose from the fire size model and that the full occurrence model performed well.

#### Sensitivity of response functions to explanatory variable selection

Beyond their limited ability to reproduce observations, intermediate models also revealed that the accuracy and shape of response functions could also be greatly impacted by modeling choices and the non-inclusion of some key effects. The response function of FWI to escaped fire number for the intermediate models were both limited in magnitude and exhibited spurious decreases, when compared to the “Full” model (Fig. 9). In particular, the “Linear-FWI” model (for which the log number of escaped fire has a linear response to FWI) was penalized by both low and high FWI, for which the actual response to FWI was respectively stronger and lower than exponential. For other intermediate models, the decrease observed above the FWI level of 65 could be explained by confounding effects between FWI and space. Indeed, highest FWI values mostly occurred in coastal populated areas where fire density was lower (at constant FWI). For the “Full” model, a small decrease was also observed at 65, but its magnitude was much smaller and was followed by slight increase above 70, thanks to the spatial model that considerably limited the impact of the confounding effect.

**FIG. 9.**
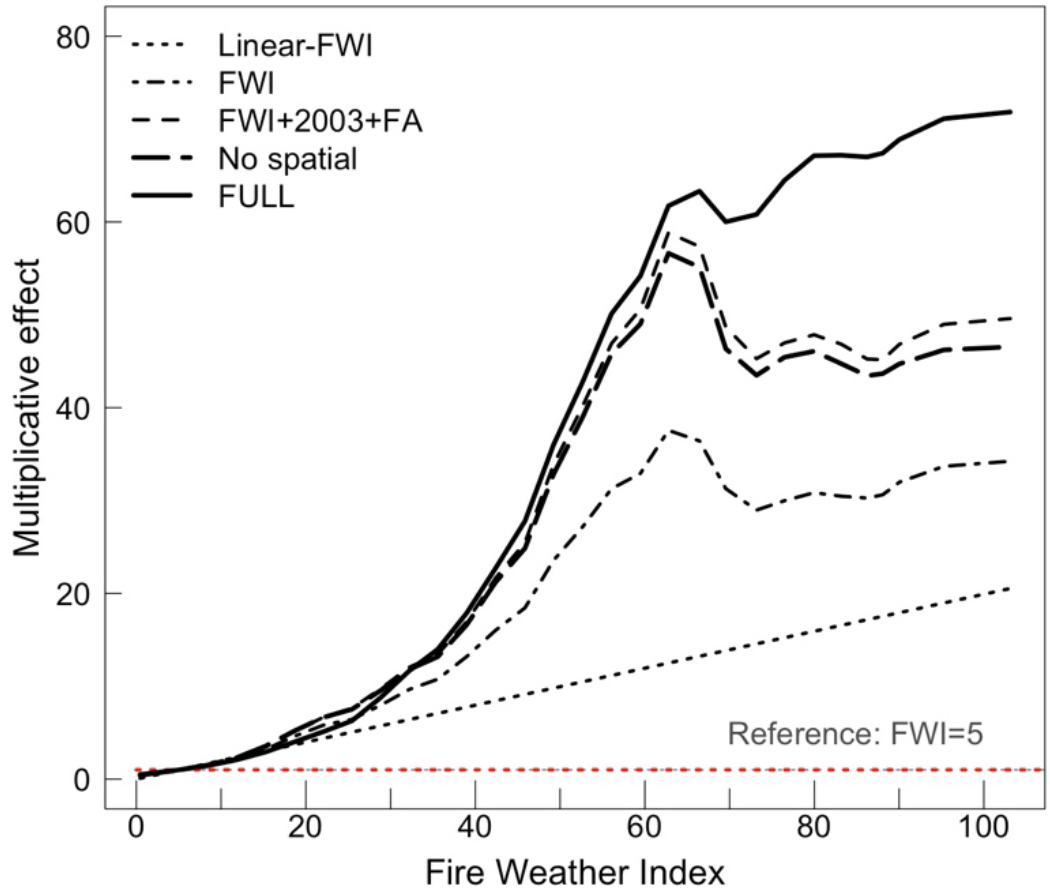
Partial effect of the FWI for the different occurrence models

### 3.3. Insights on an extreme year: the example of 2003

Here, we examine in detail why the model underestimated burnt area for the year 2003. Because most of the burnt area was caused by a very limited number of large fires, one might believe that i) observed burnt area for 2003 was very unlikely considering fire weather, but possible with a low probability (“bad luck”); ii) most fire observations were expected (i.e. consistent with the usual fire-weather relationship), but the occurrence of a few very large fires that disproportionately contributed to the total burned area. In this context, arson is frequently mentioned (and an arsonist was trully involved in a limited number of large fires in Var NUTS 3 division).

According to Fig. 8, the expected trend and the upper bound of the 95^th^ CI (0.975 quantile) for burnt areas predicted by the model were respectively of 213 and 357 km^2^, which was well below observations (610 km^2^). Similarly, the quantile 0.999, corresponding to a millennial event according to the model was 469 km^2^, which remained below observations. We can then conclude that the model failed to simulate the likelihood of observed burned area in 2003. We then analyzed time series corresponding to 2003 seasonal fire activities in Fig. 10 (similar to Fig. 4, but with 99.9^th^ confidence intervals added to show unlikely events). The predictions of escaped fires were consistent with observations at both the daily and weekly scales, which shows that fires did not escape more frequently than expected all along the season, without any exceptional week or day. However, more than six weeks were largely above the expected trends in numbers of fires larger than 100 ha and in burnt areas. Four weeks were above the 0.975 quantile and one was even above the 0.9995 quantile (week 35). This shows that 2003 was atypical during several weeks, with a very early start. The analysis of the distribution of fire size (Fig. 12), showed that all fires larger than 10 ha occurred more often than expected. Hence, even if the presence of a few very large fire constituted most of the burned area, most fires (larger than 10 ha and during most of the season) were exceptionally large with respect to observed FWI, which invalidates the arson assumption. The decrease in fire suppression efficiency with increasing escaped fire number can be invoked. However, it is important to acknowledge that it had apparently not affected the number of escaped fires, as their number was consistent with expectations.

**FIG. 10.**
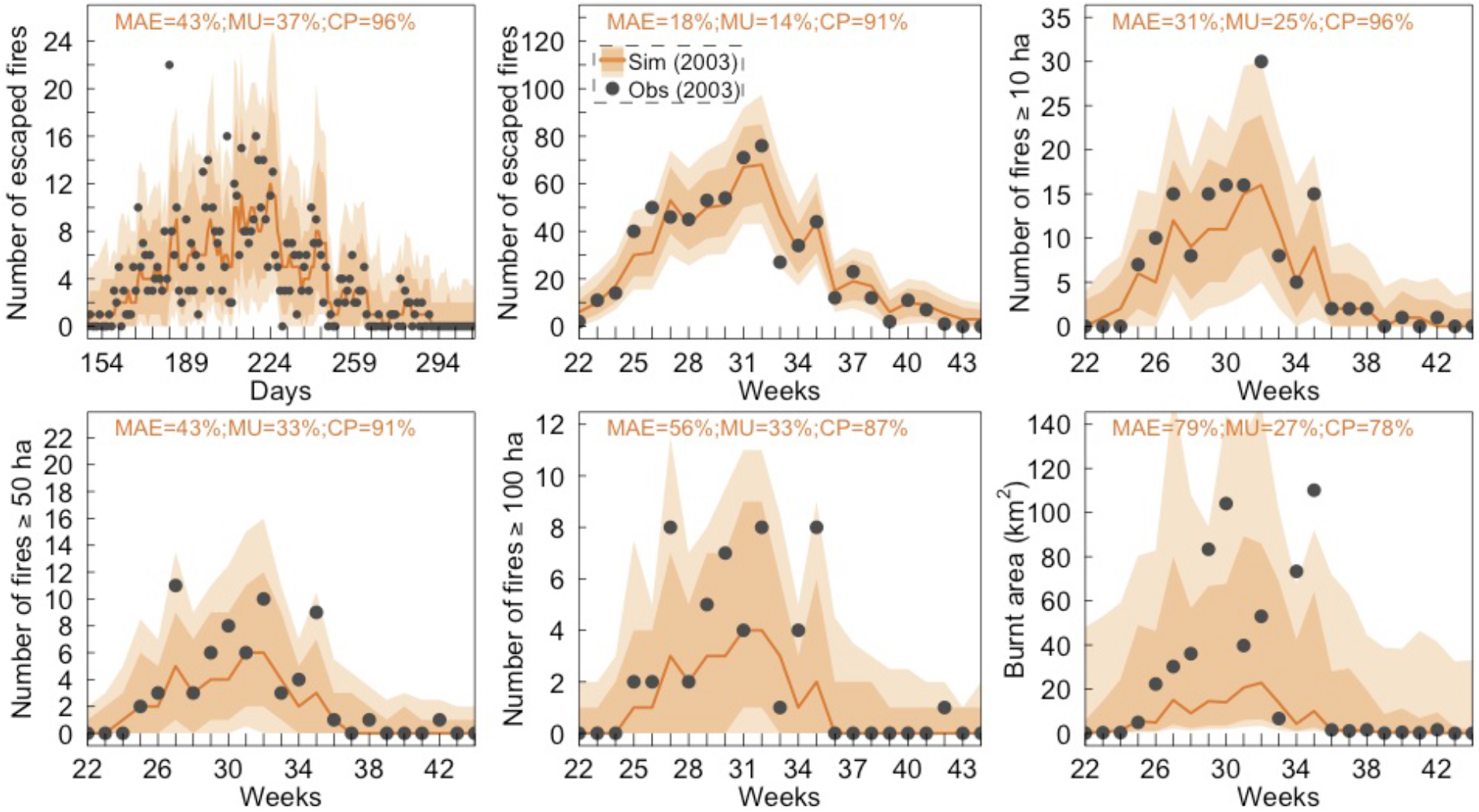
Same as FIG. 4, for year 2003. Comparison of simulated fire activity (in red) with observation (black dots): daily and weekly escaped fire numbers, as well as weekly number of fire larger than 10, 50, 100 ha and weekly burnt areas were summed for the whole study area. Expected trend (red line) was surrounded by the 95^th^ and 99.9^th^ confidence intervals in orange and light orange (computed from 1000 simulations of fire activities).

**FIG. 11.**
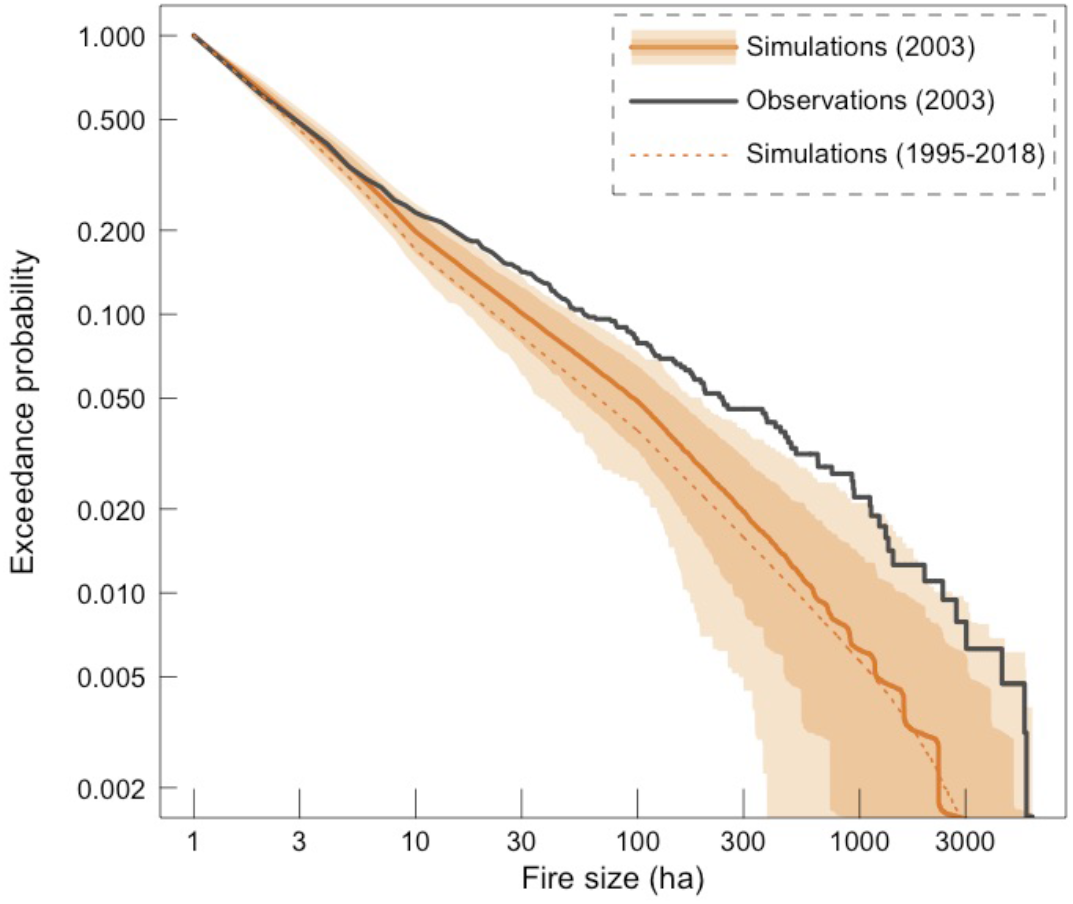
Same as FIG. 5 for year 2003. Comparison of simulated fire size distribution with observation: Expected trend (red line) was surrounded by the 95^th^ confidence interval in orange and the 99.9^th^ confidence interval (light orange), computed from 1000 simulations of fire sizes for year 2003.

**FIG. 12.**
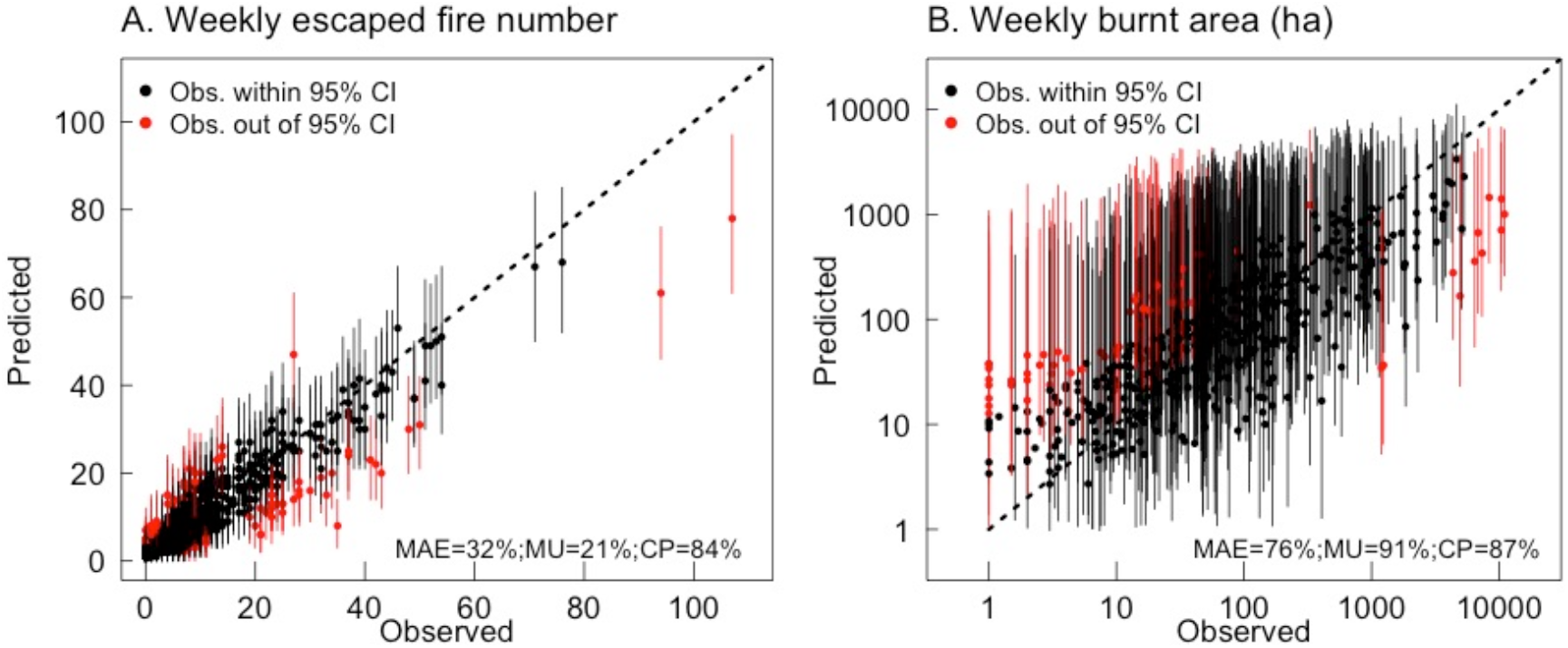
Predictability of weekly fire activities for the whole area (1995-2018).

### 3.4. Predictability of fire activity

We proposed a detailed analysis of the predictability of fire activities at various temporal and spatial aggregation scales to better understand the role of stochasticity in fire activity patterns. In general, stochasticity in observations (fire counts and sizes) typically decreases when aggregating them to larger scales, such that the nature of both observations and model predictions becomes more deterministic. Slight biases of models arising at the voxel scale may then lead to stronger biases at aggregated scales.

#### Predictability and confidence intervals at weekly scale

As shown in Fig. 4, aggregating fire activity over the whole area at the weekly scale led to reasonable confidence intervals and MAE for year 2001. In Fig. 12, we studied predictions against observations for weekly escaped fire numbers and burnt areas from 1995-2018.

The overall predictability of escaped fires at the week scale was satisfactory with a MAE of 32% and a CP of 84% (FIG. 12A). However, FIG. 12A allows to investigate why the number of escaped fires out of confidence intervals (16%) was larger than expected (5%). The majority of these numbers consisted in false “high” numbers for small observed numbers and false “low” numbers for high observed numbers. In particular, the upper bound of the simulated CI underestimated the two extreme weeks that occurred in 2001 (>90 during weeks 35 and 36, Fig. 4). Moreover, it frequently underestimated the 2 or 3 highest values of the year (typically weeks with more than 20 fires), even if corresponding predictions were generally high (see supplementary 4 for a year-by-year assessment). On the contrary, the lower bounds of simulated escaped fires were often higher than observed for low fire numbers. Such discrepancies are partly explained by stochasticity (as the “high” and “low” observations more likely constituted unfortunate and fortunate events, respectively), but they should not occur more often than 5%. A first explanation is the slight tendency of sophisticated Bayesian models -even if its components are correctly structured- to be smoother than “reality” due to the influence of the prior structure. A second explanation could be that the confidence interval widths for low and high fire numbers could also be underestimated because fire suppression was more effective when fire activity was low (means available to suppress individual fires tended to saturate when fire number increased). Indeed, variations in suppression efficiency were accounted for in the expected trends simulated by the model, because they affect the relationship between FWI and observations, but they were not accounted for in confidence interval width, because escaped fires were simulated as if they were independent between pixels. These variations could result in an over-dispersion of observations with respect to aggregated Poisson laws, and in turn, tended to limit the extents of upper and lower bounds of CI, respectively in high and low conditions.

Although the expected trend in burnt area was positively correlated with observed burnt area, MAE was very high (76 %), because of the width of the confidence intervals (MU=91 %), which illustrates the huge role of stochasticity in observed burnt areas at this scale (Fig. 12B). False “high” and “low” burnt areas happened, as for escaped fire numbers. In particular, 4 of the 7 most extreme weeks (> 2000 ha) for which the upper bound was underestimated occurred in 2003. False “high” burnt areas were mostly related to the difference between simulated and observed occurrences. Indeed, when the size component of the fire activity model was applied to observed occurrence (rather than simulated occurrence), most of the false “high” for low burnt areas of Fig. 12B were correctly simulated (see Supplementary 3.2) and CP rose up to 89% and MAE decreased to 67%.

#### Predictability at other scales

The MAE and MU are presented for 42 spatio-temporal levels of aggregation, ranging from one pixel-day to the whole area during the four years of the validation sample (Fig. 13). As expected, the MAE increased for smaller aggregations, and large fire numbers and burnt areas were more uncertain than escaped fire number (higher MU). Beyond this general trend, Fig. 13 allows us to identify which scales led to reasonable predictions (typically, those with a MAE lower than 30 %) and which were, on the contrary, subject to too much stochasticity for valuable predictions (typically, MAE above 60-70 %). In particular, the spatial aggregation drastically reduced the MAE, while sub-regional predictions remained quite poor, even for escaped fire number, when predictions were made at a finer scale than the full season. This could be partly explained by the fact that spatial patterns of ignitions have slightly evolved over the 24 years of the study period, as suggested by the lower AUC of the occurrence model in recent years (Supplementary 2). In particular, less fire activity was observed in North Corsica and more escaped fires in the western part of the basin during the recent years. The pattern followed by MU was slightly smoother regarding the effect of spatial aggregation, which suggests that the MAE of refined spatial predictions could probably be improved with a more adequate spatial model for years 2015-2018. It could be either fitted to more recent years or evolved over time instead of being constant. Also, it should be noted that the fire activity was relatively limited during the validation period, which tended to increase the magnitude of relative errors, as model uncertainty became larger when expected trends decreased. It is hence expected that the predictability of a larger number of fire events would be more accurate (even if the model failed to predict 2003).

**FIG. 13.**
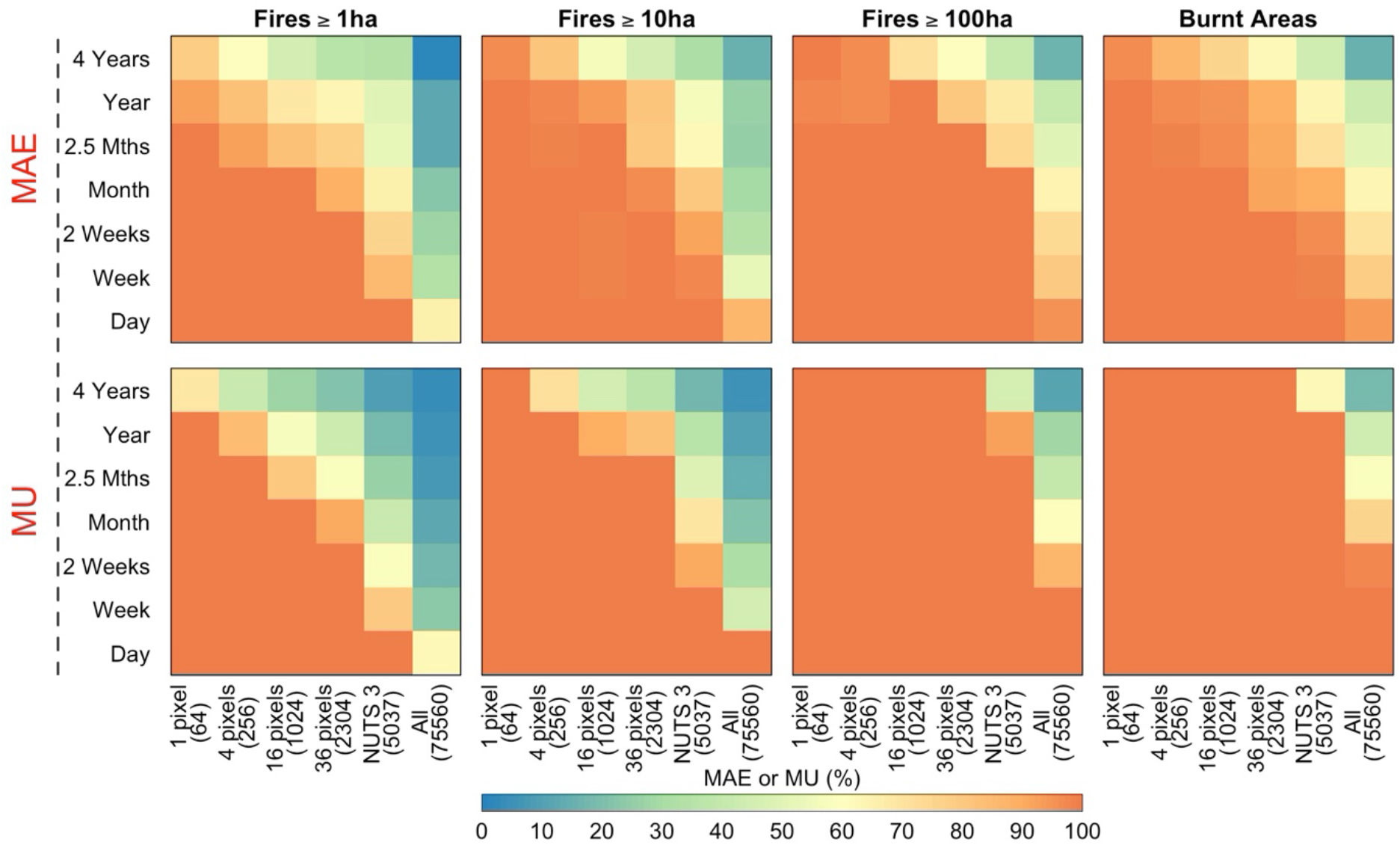
Mean Absolute Error (top) and Model Uncertainty (bottom) of the model at various spatial (1, 4, 16, 36 pixels, NUTS 3, all region, with size in km^2^ in brackets) and temporal (1 day, 1 week, two weeks,…, one year, four years) aggregations for fire numbers of 1, 10, 100 ha and burnt area for the period 2015-2018.

## 4. DISCUSSION

### Predictability of fire activity

The wildfire phenomenon combines multiple biophysical and human factors at various spatio-temporal scales, which realize toward individual events with a high degree of stochasticity. We presented a probabilistic model for regional fire activity that simulates realizations of individual fires with their size on a daily basis at 8km-pixel resolution. The model offers an unique opportunity to study the predictability of fire activity. Our results showed that small fire numbers were relatively deterministic, when considered for the whole study area on a weekly basis and can be well predicted for years following the training period. The predictability was much lower for larger fires and for sub-regional fire activities. Refined analyses are required to understand the differences between spatial and temporal aggregations in terms of predictability, but preliminary analysis suggested that it could be due to the time-variation of spatial effects.

Even if the model still present a few limitation in its rendering of stochasticity and the modeling of fire activity during heat waves (see below), ensemble simulations show promises to rate the likelihood of observed extreme.

Overall, the model highlighted that stochasticity was a major component of observed fire activities at the regional scale, so that individual events, pixels, years or fire size class, etc. are often far from deterministic and should be analyzed with caution by researchers and operational services. In this regard, one obvious outcome is that, observed monthly and annual burnt areas, which are often used to evaluate fire activity and/or fire suppression were shown to be highly random, so that we recommend to use more deterministic metrics such as number of escaped fires or fires larger than 50 or 100 ha, instead.

### Factors of fire activity

Our study demonstrated that climate and weather conditions are major drivers of fire activity in Mediterranean France and confirmed that fire-weather relationships are variable over time and space (e.g. Litell et al. 2009; Higuera et al. 2015, Ruffault and Mouillot 2015).

FWI was a suitable metric of fire danger in Mediterranean France, as suggested by earlier studies based on FWI subcomponents (Ruffault et al. 2016, Fréjaville and Curt 2017, Barbero et al. 2019), as it allowed to rate the variations in fire activity probabilities with a very encouraging accuracy. In particular, the seasonality of the fire activity was reasonably predicted, even without seasonal correction (FIG.7). Moreover, most spatial variations in fire weather relationships could likely be attributed to differences in LU-LC, rather than to potential weaknesses to the FWI. However, our study highlighted several limitations of the FWI. First, a saturation in the fire occurrence was observed beyond FWI=60 in Mediterranean France. This might be attributed to the exponential response of FWI to wind. Indeed, fires driven by strong winds are numerous in this region (Ruffault et al. 2017), but where extreme wind values (mean wind speed beyond 60 km.h^−1^) do not necessarily lead to increased fire activity. Moreover, the seasonality of the FWI did not match observations and seems to be somehow two or three weeks in advance, leading to an overestimation of fire activity in the mid of June, and an underestimation in late August, as already reported for the KBDI (Ganatsas et al. 2011). This could be explained by the desiccation processes empirically modelled in the Drought Code (a subcomponent of the FWI reflecting monthly variations in fuel moisture content), which only poorly explains Live Fuel Moisture Content (Ruffault et al. 2018b), which is increasingly recognized as an important factor of fire behavior (e.g. Pimont et al. 2019). Fire danger rating could be improved by the use of more mechanistic models for fuel moisture (Jolly and Johnson 2018, Martin StPaul et al. 2020 in prep). Finally, the most important limitation of the FWI was probably suggested by the failure of our model to simulate observed burned area in 2003. Indeed, several results suggested that the response of FWI to the 2003 drought was too soft. i) The FWI was higher in 2003 than for other years, leading to the highest predictions for 2003 (FIG.8), but this value was only slightly higher than other years such as 2001 or 2017 – even when burned areas were simulated from observed occurrence, see supplementary 3 – whereas the observations of 2003 were considerably different from other years; ii) Numbers of fire with size larger than 10 ha were systematically underestimated; iii) These fires were underestimated during the whole drought period, with almost all weeks exhibiting higher large fire activity than expected during July and August. Hence, FWI was an adequate metric of fire occurrence (once rescaled according to the fire occurrence model of study), but that it was inaccurate for fire size during the 2003 heat wave. We also hypothesize that it could be due to an inconsistent rating of fire danger between wind-driven and hot drought conditions, which could be tested by comparing the responses of fire activities in different fire weather type conditions (Ruffault et al. 2020). Further researches and analyses should help to clarify these critical points.

The Forest Area was the second important factor of spatial variations (Ganteaume and Barbero 2019), with less escaped fires in pixels with low forest area (typically below 20 %) than in those with moderate to high forest area (typically higher than 40 %), but with a saturation or slight decrease above 40 %. This patterns was explained by a trade-off between two different mechanisms explicitly accounted for in our model. First, fire numbers are limited by the area in which they can be ignited (“offset” effect). Second, fire density (i.e. number of fires per unit of forest area, *f_FA_*) decreased with forest area because human activities is often very limited (Costafreda-Aumedes et al. 2018, Syphard et al. 2007). The combination of these two mechanisms (“offset”+*f_FA_*) led to the pattern showed in Fig. 3. This pattern could be explained by some specificities of either the studied area, or the spatio-temporal characteristics of the probabilistic model, since effects can be scale-dependent (Parisien et al. 2011).

### Methodological insights for modelling of fire activity

One of the critical aspects of this work was the determination of the appropriate voxel size. Typically, very small voxels lead to issues related to both data reliability, as well as computational and memory costs, but larger voxels result in information loss. In the present model, we aimed at accounting for fine scale variations in fire weather. We ignored hourly variations in FWI and spatial variations smaller than 8 km, but proposed a general approach applicable to relatively fine scale at a reasonable cost with a sophisticated Bayesian method, thanks to the aggregation of escaped fire counts in classes. Such an aggregation led to a reduction of the dataset effective size by a factor close to 10 (4.44 million voxels led to only 497,000 observed classes), which makes it possible.

Our study also confirmed the importance of accounting for non-linearities (FWI) and of cofactors, in accordance with Woolford et al. (2011). We found that ignoring these effects could lead to a poor estimation of the effects of a main variable, because the FWI effect was either underestimated or inconsistent, when non-linearities/cofactors were ignored (FIG 9). In particular, the use of the spatial model was critical to properly address fine scale data in which spatial autocorrelation is present.

Thanks to a piecewise modelling framework, the fire size distribution and its dependency to explanatory variables were accurately modelled through flexible nonlinear response functions.

The multiple realization approach allowed to compute expected trends, but also confidence intervals and eventually return intervals of specific events. This approach is hence purely probabilistic; in particular, probabilistic statements about the uncertainty of specific components of the model (*e.g.*, the response functions) are possible, and the goodness-of-fit of models can be compared formally through probabilistic information criteria. These ensemble simulations can be evaluated against observations thanks to coverage probabilities of the confidence intervals.

Beyond the sophisticated Bayesian approach using R-INLA, the final model was simply expressed through a list of 1000 simulated sets of parameters for basic distributions (poisson, binomial, exponential, gpd) per classes of FWI, FA, pixel and date (*i.e.*, an ensemble of 1000 simulations from the posterior distribution of the “Full” model).

### Model limitations

A first limitation of the present model was the limited number of explanatory variables, especially Land Use and Land Cover factors (Forest Area only). A systematic analysis of the role of these variables has been addressed in Opitz et al. (2020), which confirmed with other studies (Ruffault & Mouillot 2017) that road length, proportion of conifers, regulated tourist area, building coverage and simultaneous presence of both building and forest strongly modify the probability of fire occurrence. In the present study, we did not explicitly account for those factors, but relied on the SPDE approach to model spatial variations in escaped fire density, since most of these Land-Use-Land-Cover (LU-LC) variables are almost static during the study period. Hence, we do not expect large performance improvement from the inclusion of these variables in the occurrence model, even if it would increase the genericity of the model (for better understanding of effective factors, or extrapolation out of the zone of model fit for example). Larger benefit could be expected of an inclusion of LU-LC variables in the size component of the model. Indeed, this component did not include any modelled spatial effect (because the size of the dataset was too small to afford it in combination with a piecewise modeling of the fire distribution), so that including LU-LC factors may improve spatial patterns in predictions. For example, differentiating shrub and forest cover is expected to increase model skills (Moreira et al. 2011). This is all the more important as our results suggested that the main limitation of the “Full” model lied in this fire size component. First, simulated burnt areas were similar when simulations of size were carried out from actual or simulated occurrence. Second, the amplitude of explanatory variables (FWI, FA) was smaller in the size component than in the escaped fire number component. Third, the size model failed to reproduce the extreme year (2003), although the occurrence model accurately simulated the weekly number of escaped fires during this period.

Overall, one very challenging aspect of fire activity modelling is the evolution of the fire-weather relationship over time, which is explained, amongst other causes, by changes in agricultural practices (including land abandonment), in LU-LC factors, in suppression means, but also in detection efficiency (Xi et al. 2019). In the present study, we used a crude approach to model the very abrupt shift in the relationship occurring between 2003 and 2005 (using a single global, not spatially resolved regression coefficient), resulting in a reduction by more than two of the escaped fire density given the same FWI value. We also noticed that spatial patterns have evolved, with less recent fire activity in North Corsica and more escaped fires in the western part of the basin during the recent years, resulting in a slight decrease in AUC after 2015 and in lower predictability skills for recent spatial patterns. These elements could call for a reduction of the temporal period used in fire activity studies to increase fire-weather relationship consistency. However, our study also highlight the need for long time series to include exceptional events such as 2003. In this context, the best approach is hence to allow for temporal variation of response functions or spatial bias components in the models. The use of spatio-temporal log-Gaussian Cox process model for fire occurrence is particularly promising to account for time-varying spatial effects (Serra et al. 2014b; Opitz et al. 2020). To date, this approach has only been applied with a yearly temporal resolution of spatial effects, and with monthly resolution of occurrence and size observations. Its implementation on a daily basis poses considerable methodological challenges with current state-of-the-art Bayesian modeling due to prohibitively high computational and numerical requirements.

Regarding the selected model of fire occurrence, some studies have concluded that the observed fire numbers could be better represented with a negative binomial model than with a Poisson model (Marchal et al. 2017; Joseph et al. 2019). Indeed, this model accounts for the overdispersion of the data, which means that the variance in observed count values can be higher than the intrinsic variance of the estimated model. This situation can arise because of a too low space-time resolution of the model and explanatory variables (such that important variability within a pixel-timestep unit is not properly taken into account), by missing explanatory variables, and spatio-temporal effects unaccounted for in the model (e.g., time-varying spatial effect). In particular, we identified in section 3.4 that the decrease in suppression efficiency when escaped fire number increase could be involved.

In this context, preliminary investigations (where we replaced the Poisson distribution with a negative binomial one) confirmed the presence of overdispersion with respect to the Poisson model, but simulations from a model fitted with a negative binomial response showed highly unrealistic values for the number of escaped fires per voxels, with simulated counts being until 10 times larger than maximum observations. This means that the model performance deteriorates by capturing overdispersion at the pixel-day scale. A more robust, but highly challenging approach would be to identify the appropriate spatial and temporal scales where overdispersion arises, and to include in the model some components appropriately resolved in space and time. Therefore, we used the Poisson-based model as the more realistic model for all our analyses, but confidence intervals can be too narrow in aggregated simulations (with CP lower than 95), with excess of “false” very high and very low fire activity.

### Model applications and perspectives

A model that enables to simulate many realizations of a highly stochastic disturbance such as forest fire offers a variety of ecological, operational and economic applications. The present study addressed the example application of the predictability, but other ecological applications are possible. The model can be used to compute and map the return interval of the disturbance more accurately than by averaging sparse observations and can encompass larger time span than the observed period. It can also be used to anticipate the potential effect of climate change, assuming that the fire-weather relationship remains constant (e.g. Wotton et al. 2003; Fargeon 2019) or to conduct attribution studies, which have to date been limited to fire weather (e.g. Barbero et al. 2020), and not activity, mostly because of the lack of consistency of observed data.

Such a model obviously offers operational applications, since weather forecasts can be used as input to forecast changes in fire activity across the landscape (Woolford et al. 2011). As weather forecasts are now quite accurate for periods of seven days, it means that the number of fires that escapes and of large fire (50-100ha) can be well anticipated, which can help organizing suppression means. It is important to recognize that the present study was carried out from the reanalysis of weather observations (SAFRAN), not from predictions (hindcasts). Uncertainties arising from weather prediction uncertainties might degrade the prediction performance. This could be tested in the future, as hindcasts for the historical periods are available. Moreover, such probabilistic approach, as well as those including time-varying spatial effects can help to identify changes in fire-weather relationship over time, related to operational aspects. This includes detection efficiency variation, as well as the evolution in strategies, tools and techniques for fire suppression (Xi et al 2019).

Statistical models of fire activity may also help gaining insight into the socio-economic impacts of changing fire regimes. At local scales, they can be used to estimate and forecast suppression costs (Preisler et al. 2011). At larger scales, economic activity and climate mitigation in the forest sector may be affected by disturbances (Lindner et al. 2010; Seidl et al. 2014). Long-term forecasts often rely on deterministic simulators where the inclusion of risk is a methodological challenge (Chudy, Sjølie, and Solberg 2016; Riviere, Caurla, and Delacote 2020): models such as ours may provide a way to consider fire activity in such assessments.

The present probabilistic approach differ from mixed models, in which the probabilistic approach for occurrence is combined with a mechanistic approach for fire size (Parisien et al. 2013; Finney et al. 2011). The mechanistic approach for fire size is based on the spread of fire contours on a landscape and allow to better account for fuel, topography and land use and cover. However, such mixed approaches could be refined by building on the strengths of the occurrence model presented here (e.g. SPDE approach for spatial variations) and on their mechanistic model for fire spread.

## Conclusion

This study proposed a comprehensive framework for fire activity in Mediterranean France, but the methodology was general enough to be applied to many regions or landscapes. Model performance was very encouraging, especially for escaped fire numbers, and allowed to better understand the role of stochasticity in fire activity in Mediterranean France.

We raised a few challenges in terms of explanatory variables (because of its failure in 2003 and potential limitation of the FWI for fire size assessment) and because of time variations in spatial effects.

Our analyses also suggested some bases for future researches, including a variety of ecological, operational and economic applications.

## Author contributions

HF led an earlier manuscript describing the model development in her PhD dissertation (Fargeon 2019), supervised by FP, JLD and NM. TO, HF and FP built the models and ran the simulations with RINLA. FP, TO and JR wrote the present version of the manuscript with the help of other co-authors.

## Acknowledgements

This study was done in HF’s PhD, which was funded by French Ministry of Agriculture. We thank Fire Department of the French Forest Fire Service for their helpful comments when developing the model and writing the manuscript (Yvon Duché, Rémi Savazzi, Benoit Reymont and Marion Toutchkoff).

