## Supplementaries for "Prediction of regional wildfire activity with a probabilistic Bayesian framework"

**Supplementary materials**

**Supplementary 1.** Map of forest cover in the studied area

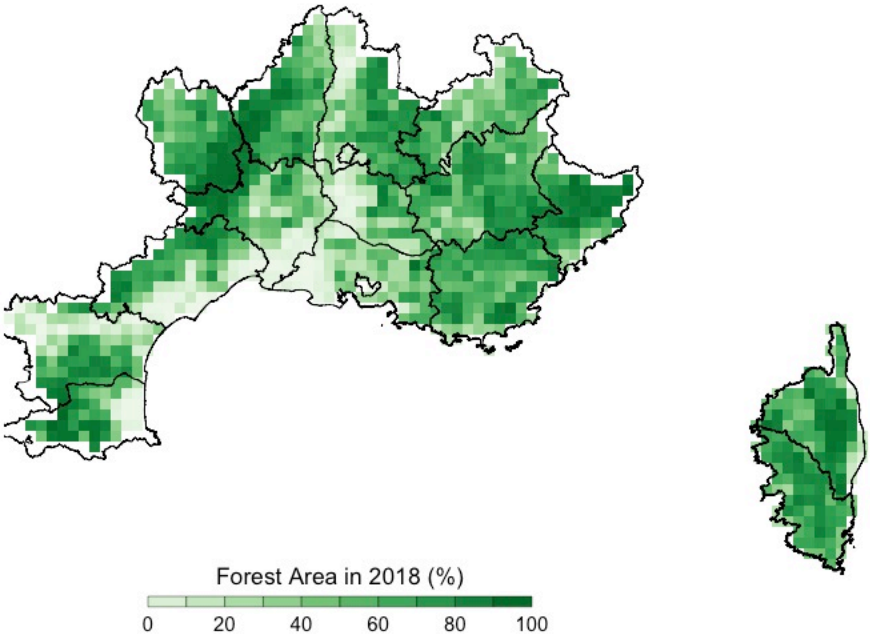

**Supplementary 2.**

Here, we described the metrics used to evaluate the intermediate models and how they led to the selection of the "Full" model. In Table S2.1, the deviance information criteria (DIC and WAIC) both decreased when more explanatory variables were included in the occurrence model component, showing that all variables led to significant improvements in the goodness-of-fit of the model. The occurrence model "FWI+2003+FA+WEEK+SPATIAL" was the one used in the "Full" fire activity model.
It provides an AUC (corresponding to the event "at least one escaped fire occurred in the voxel") higher than 0.8 for both the training and evaluation sample, and substantially improved on the other models. Overall, AUC values were similar for the training and evaluation samples, which suggests that the models were generally not overfitted. The difference AUC was slightly stronger when the SPATIAL effect was included in the validation (0.816) sample as compared to the training sample (0.857), which suggests that fire activity, and in particular spatial biases not explained by explanatory variables, might have slightly evolved over time.
Explanatory variables such as "Post-2003", FA and WEEK effects reduced the value of the Deviance Information criterion, but did not increase the AUC (and sometimes even slightly reduced it). This highlights that the main explanatory variables were FWI and the SPATIAL effect, even if predictions were biased when the other effects were not included (FIG. 7).

**Table S2.1. Fire occurrence model component fit**

| | Deviance Information Criterion (DIC) | Watanabe-Akaike Information Criterion (WAIC) | AUC for $\boldsymbol{P(N_1 \geq 1)}$ **(year≤2014)** | AUC for $\boldsymbol{P(N_1 \geq 1)}$ **(year>2014)** |
|---|---|---|---|---|
| Null | 70679 | 70679 | 0.500 | 0.500 |
| FWI-Linear | 64174 | 64174 | 0.771 | 0.775 |
| FWI | 63222 | 63225 | 0.785 | 0.784 |
| FWI+2003 | 62297 | 62300 | 0.798 | 0.781 |
| FWI+2003+FA | 61788 | 61794 | 0.781 | 0.748 |
| FWI+2003+FA+WEEK | 61548 | 61556 | 0.784 | 0.750 |
| FWI+2003+FA+WEEK +SPATIAL | 54143 | 54127 | 0.857 | 0.816 |

Tables S2.2 to S2.4 show similar statistics for the logistic models of the probability of exceeding size thresholds. For FWI+FA (which was used in the "Full" model), the DIC values were the lowest and AUC values were the highest. Also, AUC values were similar for both samples. The AUC for the 10-ha exceedance thresholds remained relatively poor, even for the "FWI+FA" model (0.66), which underscores the high stochasticity in the fire sizes, and the difficulty to predict them with high accuracy.

**Table S2.2. Exceedance probability $\boldsymbol{P(S \geq 10|S \geq 1)}$ fit (fire size model component)**

| | Deviance Information Criterion (DIC) | Watanabe-Akaike Information Criterion (WAIC) | AUC for $\boldsymbol{P(S \geq 10|S \geq 1)}$ **(year≤2014)** | AUC for $\boldsymbol{P(S \geq 10|S \geq 1)}$ **(year>2014)** |
|---|---|---|---|---|
| Null | 5519 | 5519 | 0.495 | 0.493 |
| FWI-Linear | 5349 | 5349 | 0.622 | 0.594 |
| FWI | 5345 | 5345 | 0.621 | 0.578 |
| FWI+FA | 5230 | 5329 | 0.660 | 0.631 |

**Table S2.3. Exceedance probability $\boldsymbol{P(S \geq 100|S \geq 1)}$ fit (fire size model component)**

| | Deviance Information | Watanabe-Akaike Information | AUC for $\boldsymbol{P(S \geq 100|S \geq 1)}$ | AUC for $\boldsymbol{P(S \geq 100|S \geq 1)}$ |
|---|---|---|---|---|

| | Criterion (DIC) | Criterion (WAIC) | **(year≤2014)** | **(year>2014)** |
|---|---|---|---|---|
| Null | 1945 | 1945 | 0.504 | 0.488 |
| FWI-Linear | 1813 | 1813 | 0.708 | 0.756 |
| FWI | 1807 | 1807 | 0.717 | 0.729 |
| FWI+FA | 1753 | 1753 | 0.754 | 0.764 |

**Table S2.4. Exceedance probability $P(S \geq 1000|S \geq 1)$) fit (fire size model component)**

| | Deviance Information Criterion (DIC) (saturated) | Watanabe-Akaike Information Criterion (WAIC) | AUC for $P(S \geq 1000|S \geq 1)$ **(year≤2014)** | AUC for $P(S \geq 1000|S \geq 1)$ **(year>2014)** |
|---|---|---|---|---|
| Null | 412 | 412 | 0.492 | 0.220 |
| FWI-Linear | 400 | 400 | 0.666 | 0.756 |
| FWI | 395 | 395 | 0.716 | 0.729 |
| FWI+FA | 390 | 390 | 0.747 | 0.752 |

Tables S2.5 to S2.7 show similar statistics for the exponential distributions in the first three segments. For FWI+FA (which was used in the "Full" model), DIC values were the lowest and AUC values were the highest. Also, AUC values were similar for both samples. Here, AUC corresponded to the event "fire size was larger than the middle of the segment", which combined both the exceedance and the exponential distribution. The AUC for the 5-ha exceedance thresholds remained relatively poor, even for the "FWI+FA" model (0.615).

**Table S2.5. Exponential distribution between 1 and 10ha fit (fire size model component)**

| | Deviance Information Criterion (DIC) | Watanabe-Akaike Information Criterion (WAIC) | AUC for $P(S \geq 5|S \geq 1)$ **(year≤2014)** | AUC for $P(S \geq 5|S \geq 1)$ **(year>2014)** |
|---|---|---|---|---|
| Null | 11673 | 11672 | 0.498 | 0.473 |
| FWI-Linear | 11556 | 11555 | 0.595 | 0.588 |
| FWI | 11561 | 11559 | 0.592 | 0.585 |
| FWI+FA | 11486 | 11482 | 0.615 | 0.617 |

**Table S2.6. Exponential distribution between 10 and 100ha fit (fire size model component)**

| | Deviance Information Criterion (DIC) | Watanabe-Akaike Information Criterion (WAIC) | AUC for $P(S \geq 50|S \geq 1)$ **(year≤2014)** | AUC for $P(S \geq 50|S \geq 1)$ **(year>2014)** |
|---|---|---|---|---|
| Null | 2341 | 2341 | 0.522 | 0.511 |
| FWI-Linear | 2392 | 2392 | 0.683 | 0.674 |
| FWI | 2344 | 2343 | 0.686 | 0.656 |
| FWI+FA | 2331 | 2329 | 0.718 | 0.684 |

**Table S2.7. Exponential distribution between 100 and 1000ha fit (fire size model component)**

| | Deviance Information Criterion (DIC) | Watanabe-Akaike Information Criterion (WAIC) | AUC for $P(S \geq 500|S \geq 1)$ **(year≤2014)** | AUC for $P(S \geq 500|S \geq 1)$ **(year>2014)** |
|---|---|---|---|---|
| Null | 505 | 505 | 0.462 | 0.460 |

| | | | | |
|---|---|---|---|---|
| FWI-Linear | 507 | 507 | 0.735 | 0.753 |
| FWI | 506 | 506 | 0.737 | 0.723 |
| FWI+FA | 504 | 503 | 0.802 | 0.757 |

Table S2.8 shows similar statistics for the Generalized Pareto Distribution used for the last segment (>1000 ha). Because the number of such fires was very small (only 33 and 7 in the training and validation samples, respectively), AIC increased when explanatory variables FWI and FA were included, and interpreting AUCs for the post-2014 period is not useful. We also point out that such model comparison criteria based on asymptotic large-sample theory in statistics should be interpreted with care. To complete the above analysis, we also estimated the variability of estimated parameters through a more robust procedure ("jackknife", where the model is re-estimated several times by holding out a single observation each time). The resulting estimates were all positive for the FWI-coefficient, but no clear pattern of positive or negative effect arises for FA; this has led us to keep only FWI but not FA as explanatory variable in the Full model for the Generalized Pareto Distribution of the size excesses over the highest threshold 1000.
AUC were presented only for the training sample, because the number of fires larger than 2000 ha was only 2, which was too small to compute robust AUC.

**Table S2.8. Generalized Pareto Distribution beyond 1000 ha (fire size model component)**

| | AIC | AUC for $\boldsymbol{P(S \geq 2000 \mid S \geq 1)}$ **(year≤2014)** | AUC for $\boldsymbol{P(S \geq 2000 \mid S \geq 1)}$ **(year>2014)** |
|---|---|---|---|
| Null | 41.2 | 0.483 | NA[1] |
| FWI | 42.6 | 0.710 | NA[1] |
| FWI+FA | 44.5 | 0.793 | NA[1] |

[1] The number of fire larger than 2000 ha in the validation dataset is too small to compute AUC (< 8)

**Supplementary 3.** Same figure as 8 and 12B, but when burnt areas were simulated from observed escaped fires (based on the size component of the fire activity model). It shows that using observed occurrence did not strongly increase the accuracy of simulated burnt areas, suggesting that the limitation in burnt area simulations mostly arose from the fire size model and that the full occurrence model performed well.

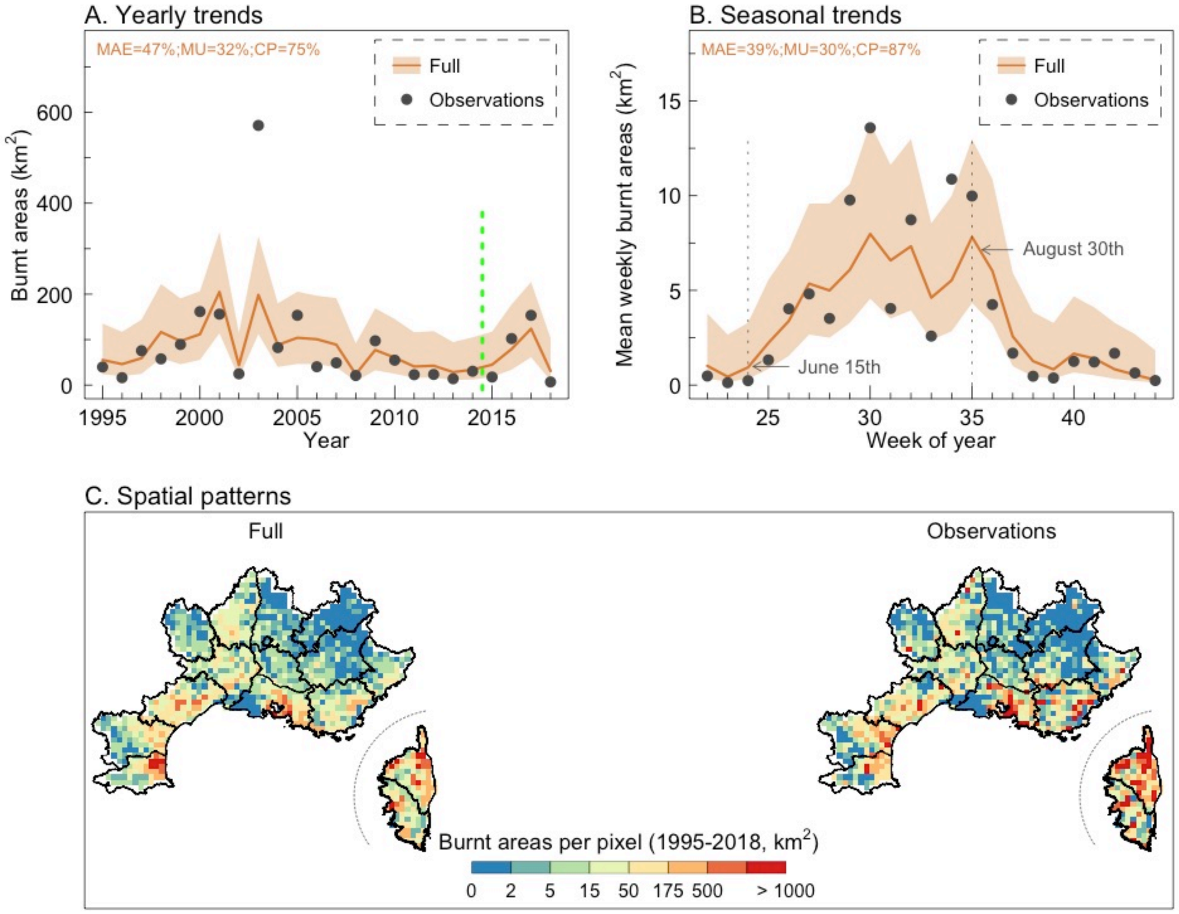


FIG. S3.1. Same figure as FIG. 8, but burnt areas were simulated from observed escaped fires, rather than from simulated escaped fire as in FIG.8.

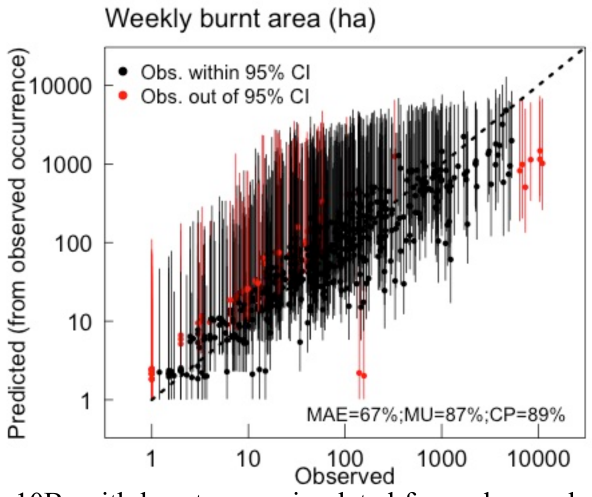


FIG. S3.2. Same as FIG. 10B, with burnt areas simulated from observed occurrence. Predictability of weekly burnt areas for the whole area (from observed escaped fires).

**Supplementary 4.** Same as FIG. 4 and 10 for other years. Comparison of yearly simulated fire activity (in red) with observation (black dots): daily and weekly escaped fire numbers, as well as weekly number of fire larger than 10, 50, 100 ha and weekly burnt areas were summed for the whole study area. Expected trend (red line) was surrounded by the 95$^{th}$ confidence interval in orange (computed from 1000 simulations of fire activities for all voxels of the year of interest).

**A. 1995**

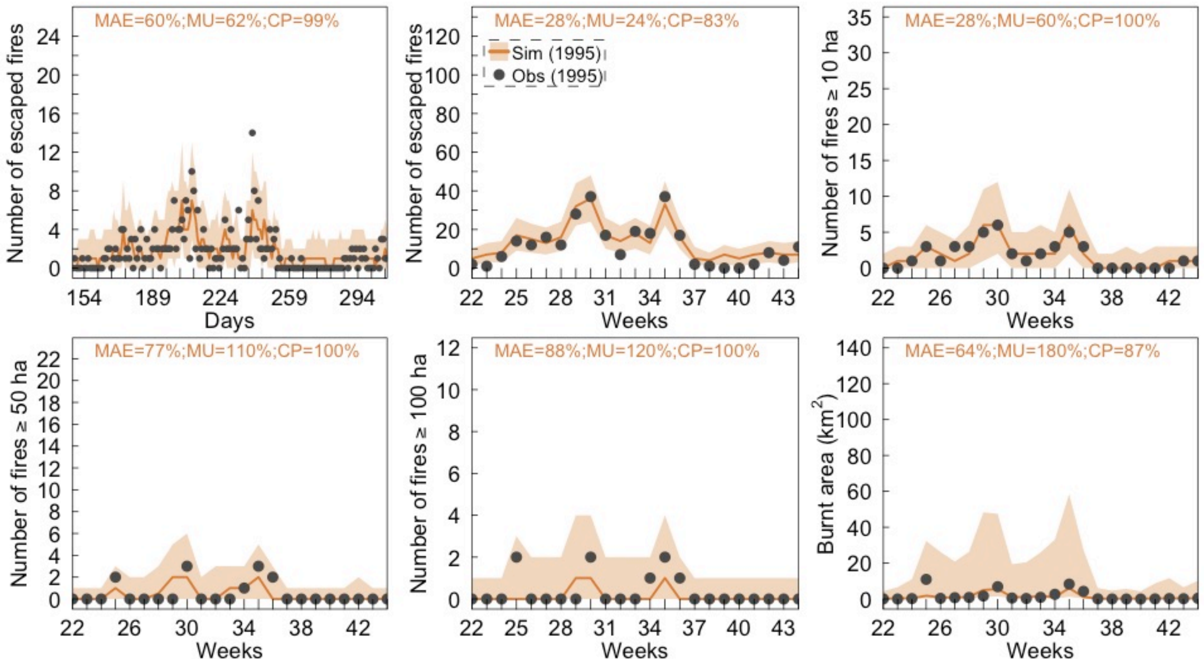


**B. 1996**

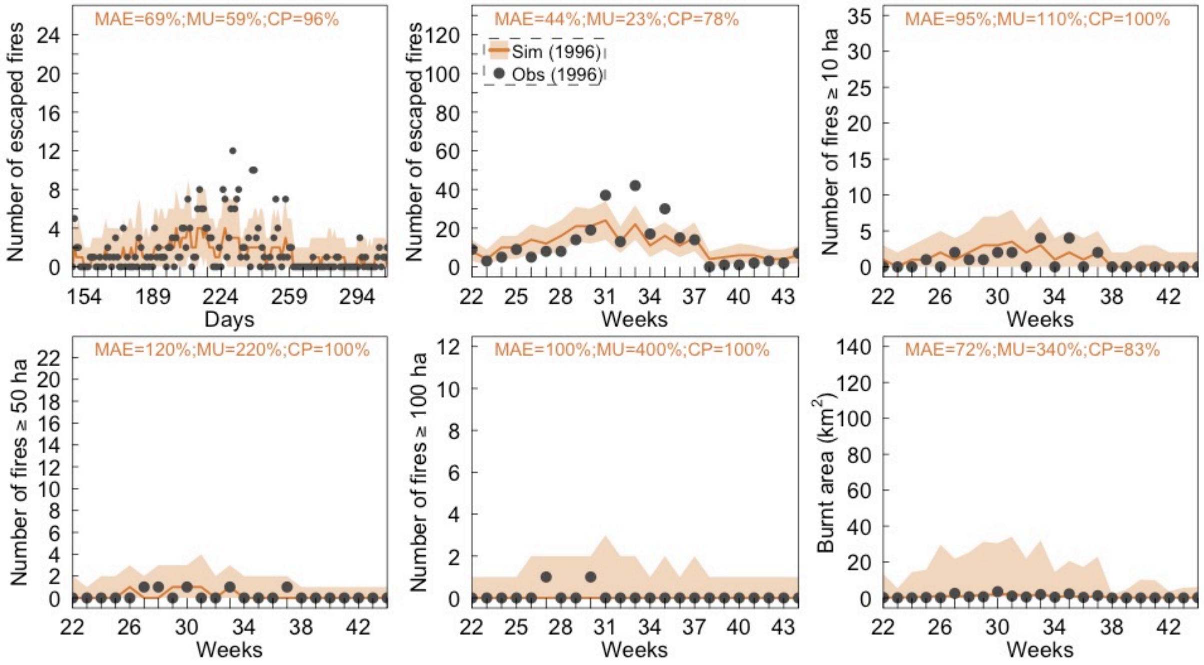


**C. 1997**

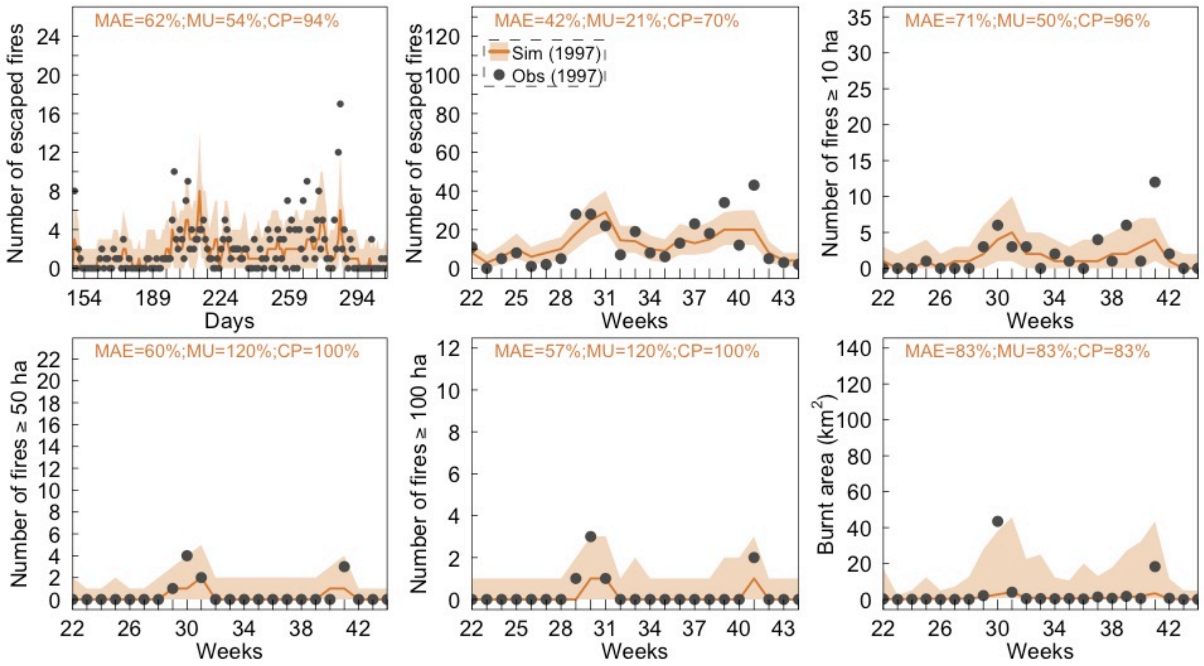
MAE=62%;MU=54%;CP=94%
Number of escaped fires
Days
MAE=42%;MU=21%;CP=70%
Sim (1997)
Obs (1997)
Number of escaped fires
Weeks
MAE=71%;MU=50%;CP=96%
Number of fires ≥ 10 ha
Weeks
MAE=60%;MU=120%;CP=100%
Number of fires ≥ 50 ha
Weeks
MAE=57%;MU=120%;CP=100%
Number of fires ≥ 100 ha
Weeks
MAE=83%;MU=83%;CP=83%
Burnt area (km2)
Weeks

**D. 1998**

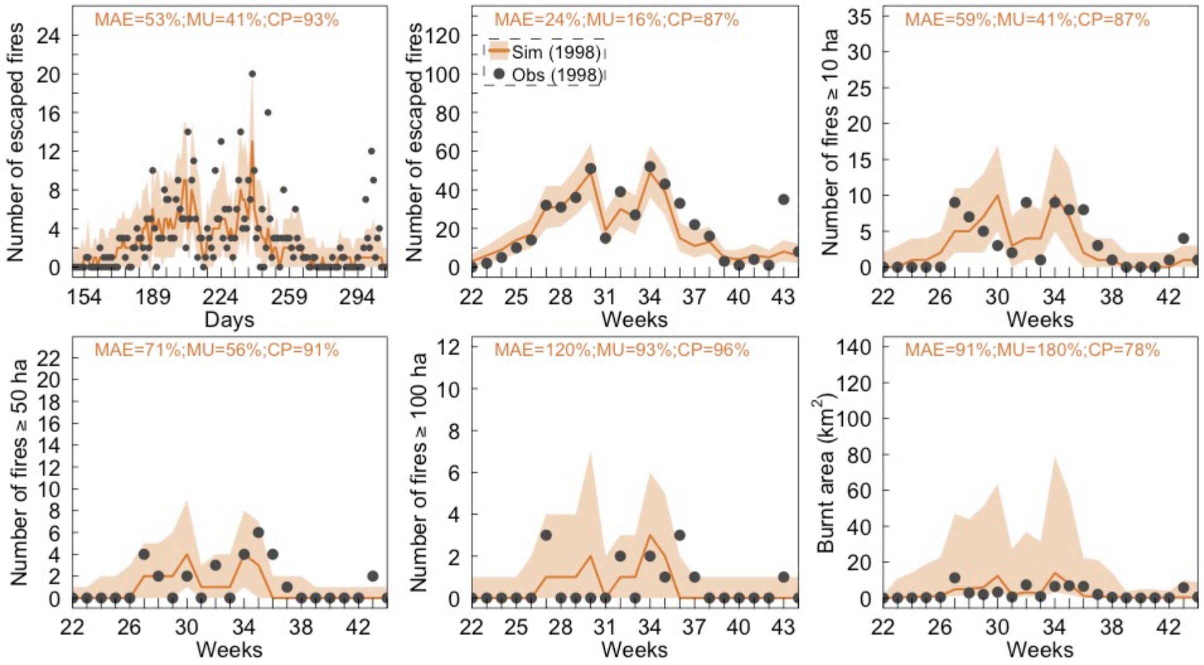
MAE=53%;MU=41%;CP=93%
Number of escaped fires
Days
MAE=24%;MU=16%;CP=87%
Sim (1998)
Obs (1998)
Number of escaped fires
Weeks
MAE=59%;MU=41%;CP=87%
Number of fires ≥ 10 ha
Weeks
MAE=71%;MU=56%;CP=91%
Number of fires ≥ 50 ha
Weeks
MAE=120%;MU=93%;CP=96%
Number of fires ≥ 100 ha
Weeks
MAE=91%;MU=180%;CP=78%
Burnt area (km2)
Weeks

**E. 1999**

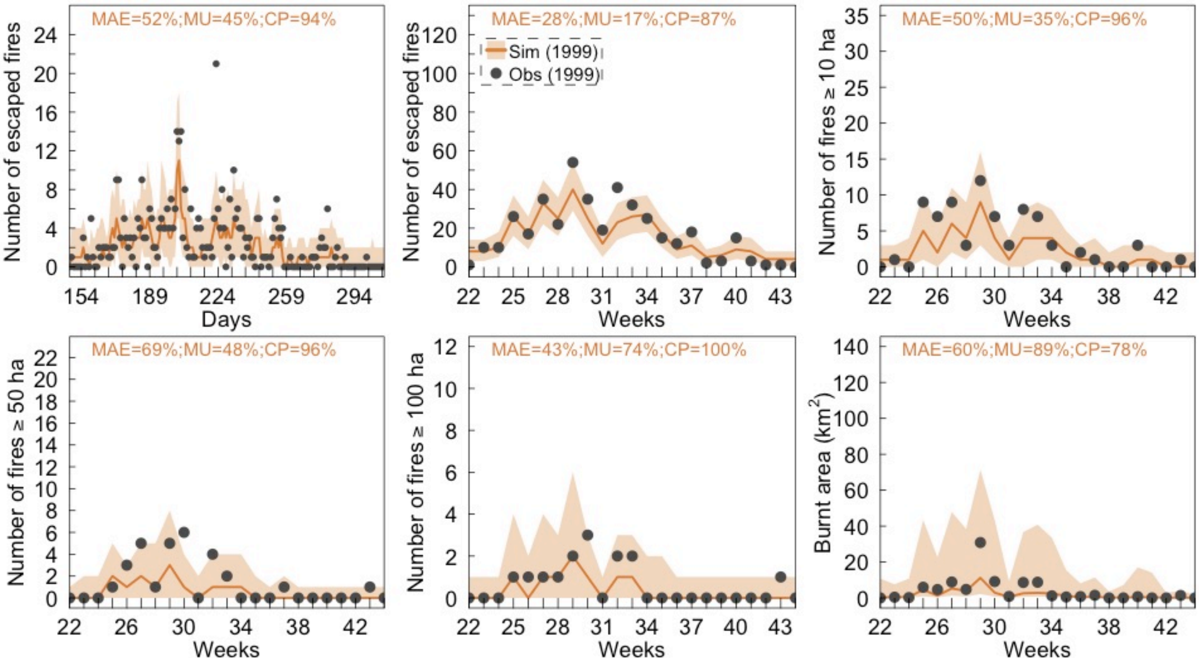
MAE=52%;MU=45%;CP=94%
Number of escaped fires
Days
MAE=28%;MU=17%;CP=87%
Sim (1999)
Obs (1999)
Number of escaped fires
Weeks
MAE=50%;MU=35%;CP=96%
Number of fires ≥ 10 ha
Weeks
MAE=69%;MU=48%;CP=96%
Number of fires ≥ 50 ha
Weeks
MAE=43%;MU=74%;CP=100%
Number of fires ≥ 100 ha
Weeks
MAE=60%;MU=89%;CP=78%
Burnt area (km$^2$)
Weeks

**F. 2000**

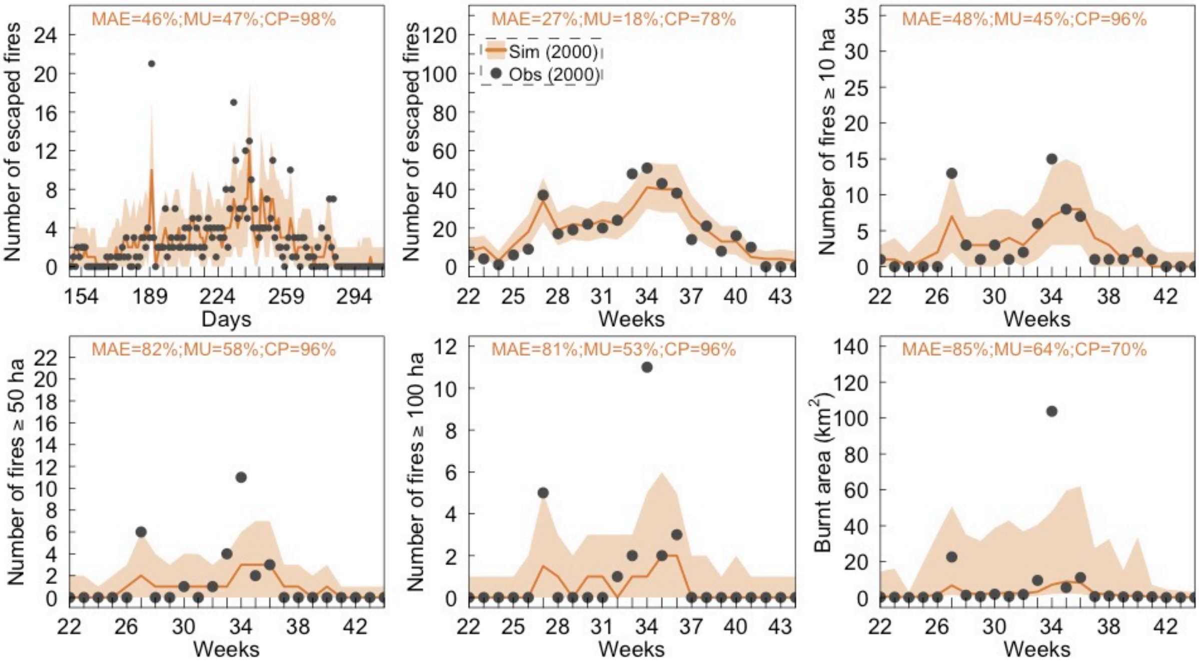
MAE=46%;MU=47%;CP=98%
Number of escaped fires
Days
MAE=27%;MU=18%;CP=78%
Sim (2000)
Obs (2000)
Number of escaped fires
Weeks
MAE=48%;MU=45%;CP=96%
Number of fires ≥ 10 ha
Weeks
MAE=82%;MU=58%;CP=96%
Number of fires ≥ 50 ha
Weeks
MAE=81%;MU=53%;CP=96%
Number of fires ≥ 100 ha
Weeks
MAE=85%;MU=64%;CP=70%
Burnt area (km$^2$)
Weeks

**G. 2001**

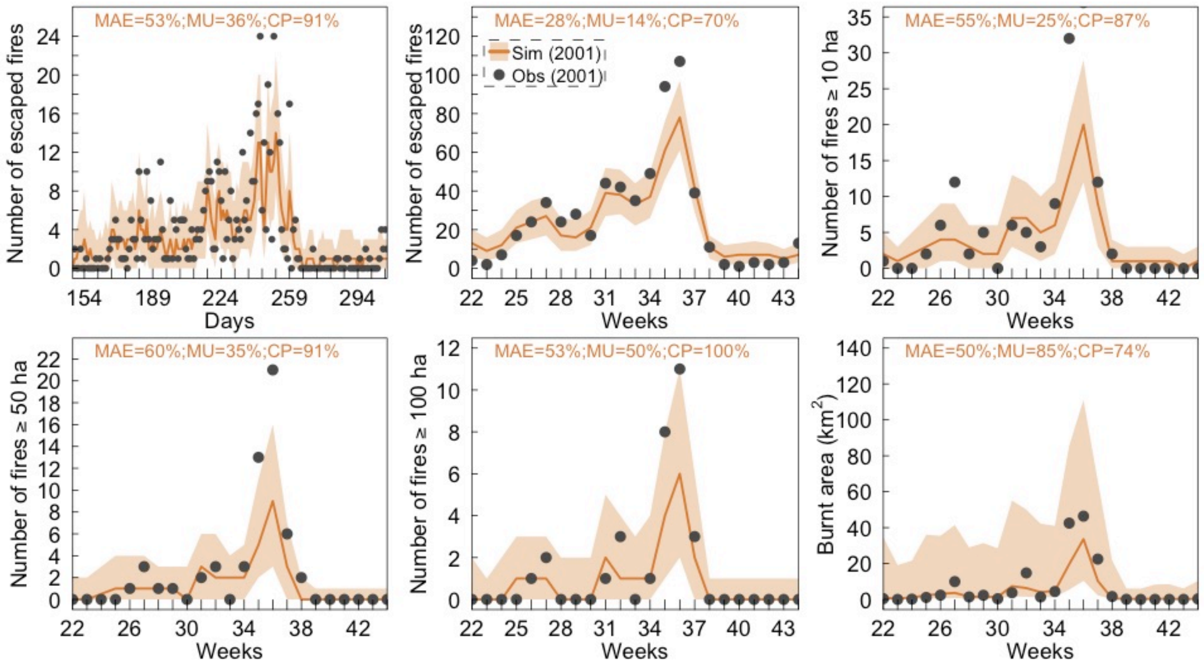

MAE=53%;MU=36%;CP=91%
Number of escaped fires
Days
MAE=28%;MU=14%;CP=70%
Sim (2001)
Obs (2001)
Number of escaped fires
Weeks
MAE=55%;MU=25%;CP=87%
Number of fires ≥ 10 ha
Weeks
MAE=60%;MU=35%;CP=91%
Number of fires ≥ 50 ha
Weeks
MAE=53%;MU=50%;CP=100%
Number of fires ≥ 100 ha
Weeks
MAE=50%;MU=85%;CP=74%
Burnt area (km2)
Weeks


**H. 2002**

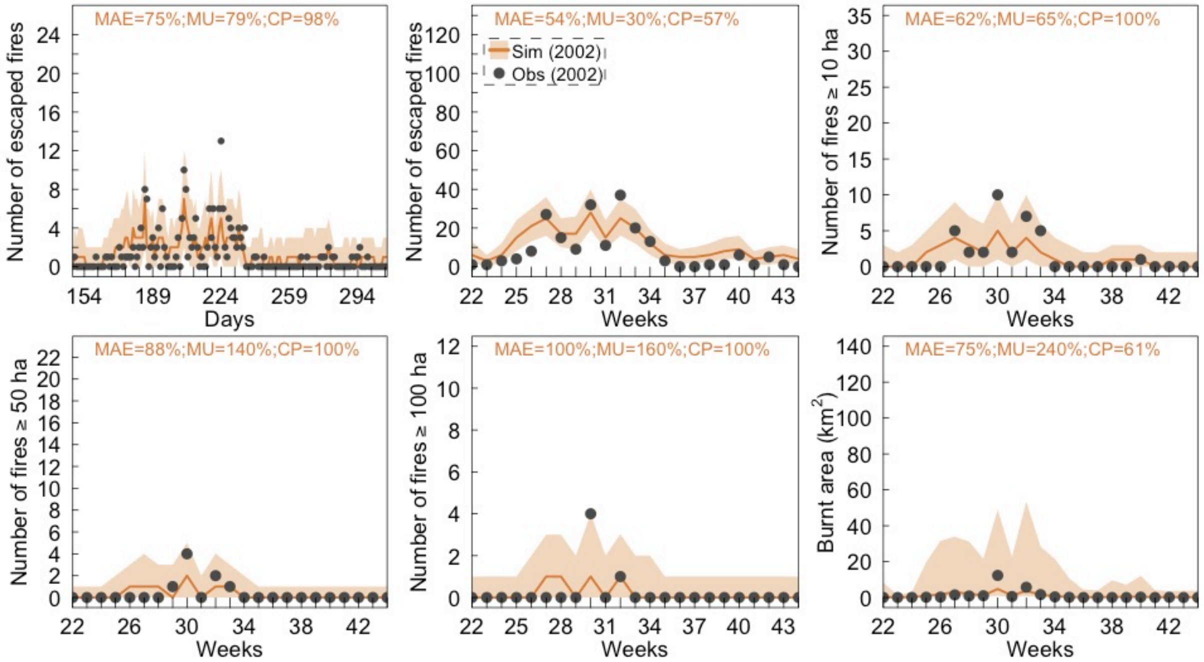

MAE=75%;MU=79%;CP=98%
Number of escaped fires
Days
MAE=54%;MU=30%;CP=57%
Sim (2002)
Obs (2002)
Number of escaped fires
Weeks
MAE=62%;MU=65%;CP=100%
Number of fires ≥ 10 ha
Weeks
MAE=88%;MU=140%;CP=100%
Number of fires ≥ 50 ha
Weeks
MAE=100%;MU=160%;CP=100%
Number of fires ≥ 100 ha
Weeks
MAE=75%;MU=240%;CP=61%
Burnt area (km2)
Weeks


**I. 2003**

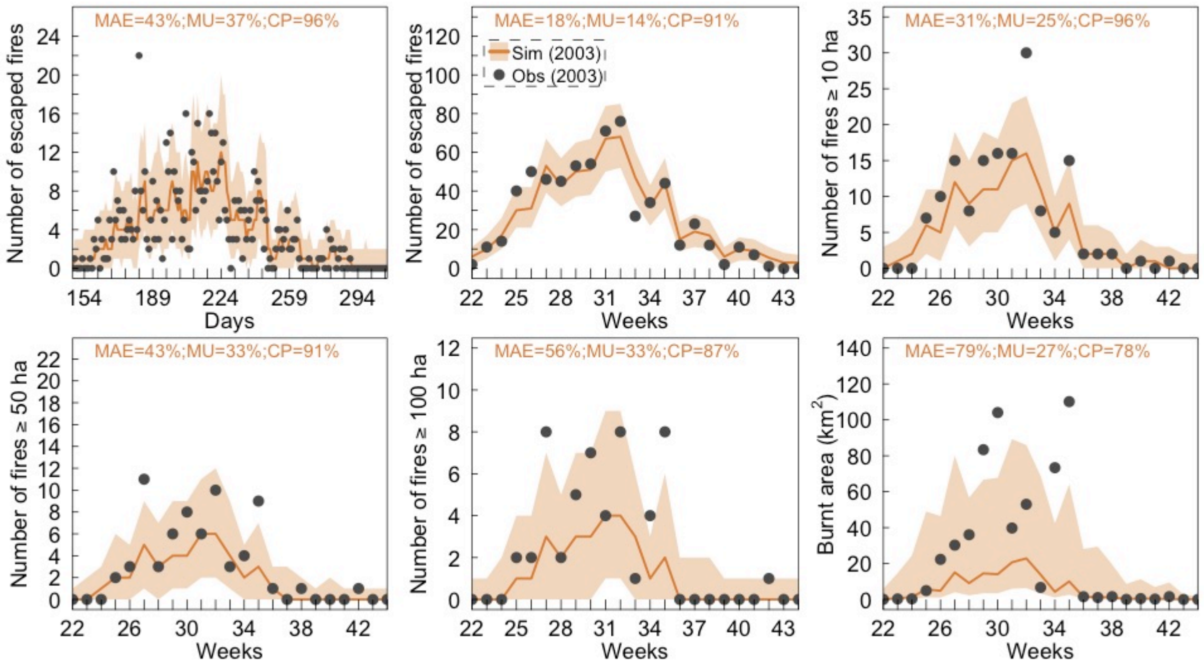
MAE=43%;MU=37%;CP=96%
Number of escaped fires
Days
MAE=18%;MU=14%;CP=91%
Sim (2003)
Obs (2003)
Number of escaped fires
Weeks
MAE=31%;MU=25%;CP=96%
Number of fires ≥ 10 ha
Weeks
MAE=43%;MU=33%;CP=91%
Number of fires ≥ 50 ha
Weeks
MAE=56%;MU=33%;CP=87%
Number of fires ≥ 100 ha
Weeks
MAE=79%;MU=27%;CP=78%
Burnt area (km2)
Weeks

**J. 2004**

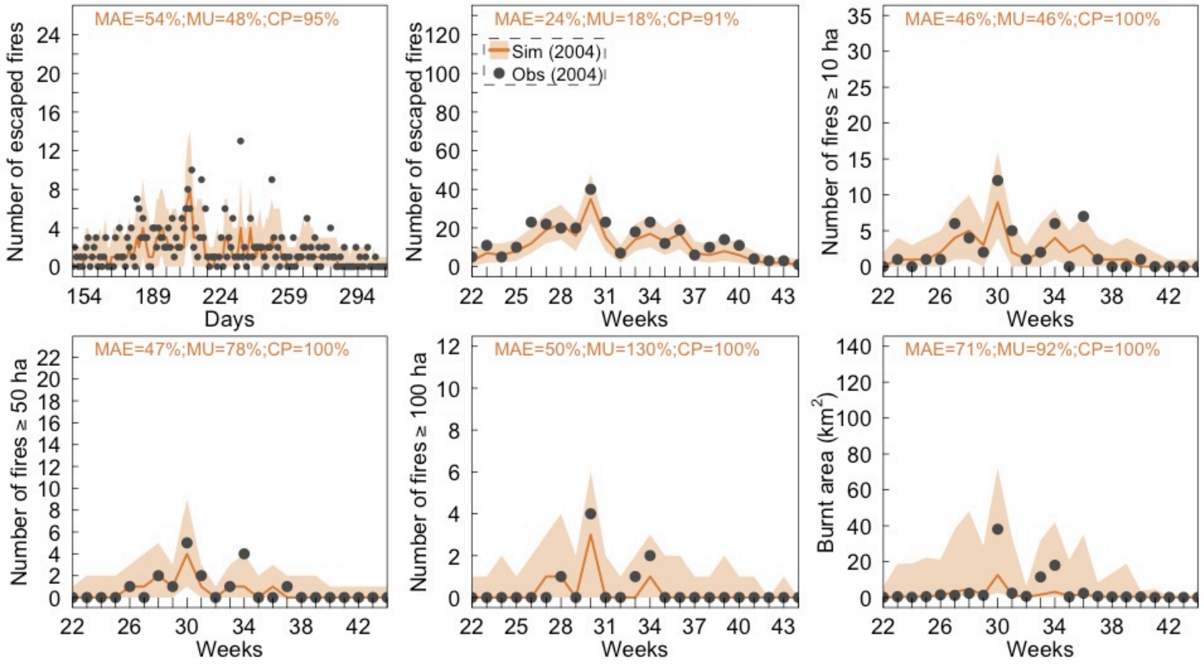
MAE=54%;MU=48%;CP=95%
Number of escaped fires
Days
MAE=24%;MU=18%;CP=91%
Sim (2004)
Obs (2004)
Number of escaped fires
Weeks
MAE=46%;MU=46%;CP=100%
Number of fires ≥ 10 ha
Weeks
MAE=47%;MU=78%;CP=100%
Number of fires ≥ 50 ha
Weeks
MAE=50%;MU=130%;CP=100%
Number of fires ≥ 100 ha
Weeks
MAE=71%;MU=92%;CP=100%
Burnt area (km2)
Weeks

**K. 2005**

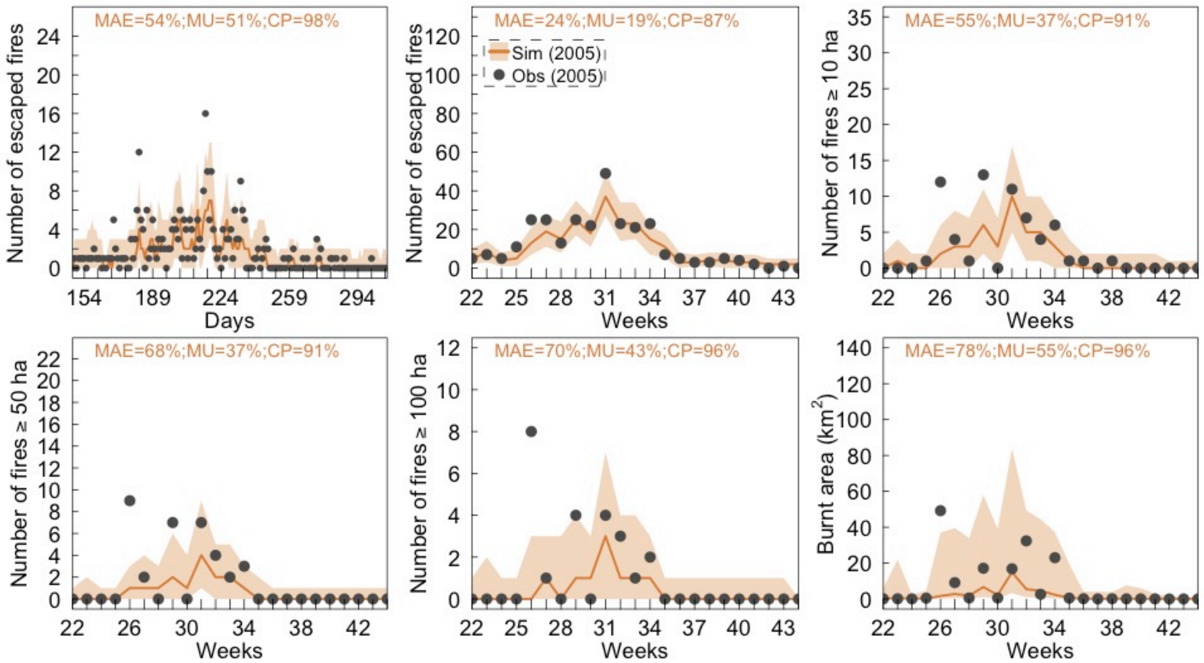
MAE=54%;MU=51%;CP=98%
Number of escaped fires
Days
MAE=24%;MU=19%;CP=87%
Sim (2005)
Obs (2005)
Number of escaped fires
Weeks
MAE=55%;MU=37%;CP=91%
Number of fires ≥ 10 ha
Weeks
MAE=68%;MU=37%;CP=91%
Number of fires ≥ 50 ha
Weeks
MAE=70%;MU=43%;CP=96%
Number of fires ≥ 100 ha
Weeks
MAE=78%;MU=55%;CP=96%
Burnt area (km2)
Weeks

**L. 2006**

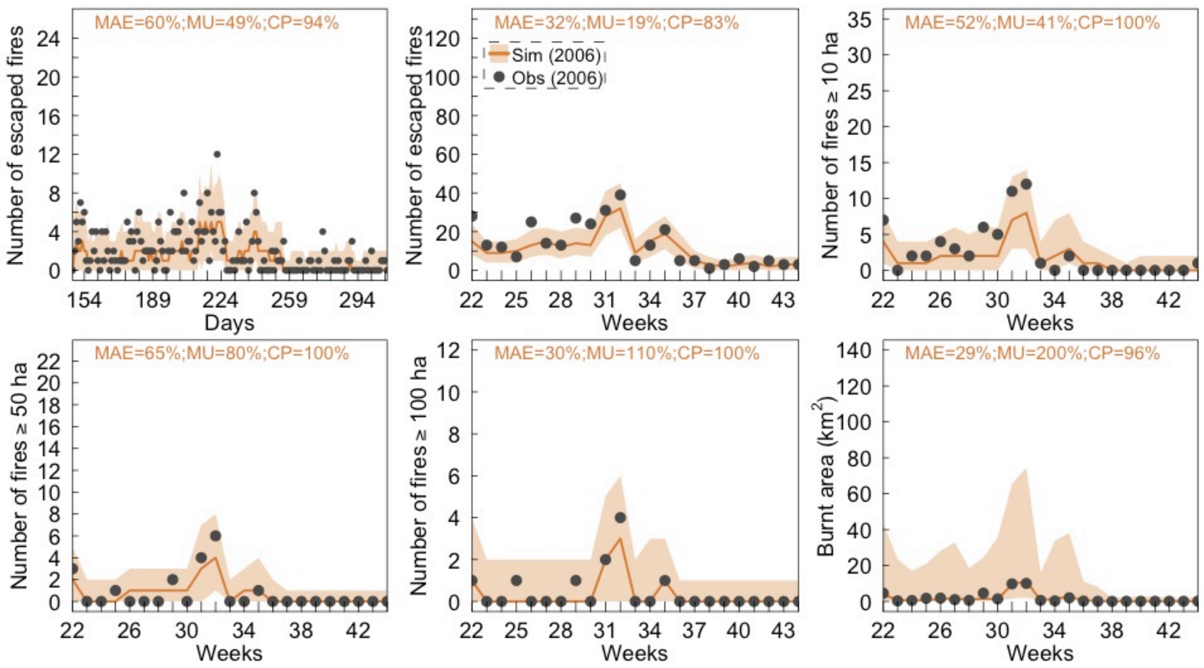
MAE=60%;MU=49%;CP=94%
Number of escaped fires
Days
MAE=32%;MU=19%;CP=83%
Sim (2006)
Obs (2006)
Number of escaped fires
Weeks
MAE=52%;MU=41%;CP=100%
Number of fires ≥ 10 ha
Weeks
MAE=65%;MU=80%;CP=100%
Number of fires ≥ 50 ha
Weeks
MAE=30%;MU=110%;CP=100%
Number of fires ≥ 100 ha
Weeks
MAE=29%;MU=200%;CP=96%
Burnt area (km2)
Weeks

**M. 2007**

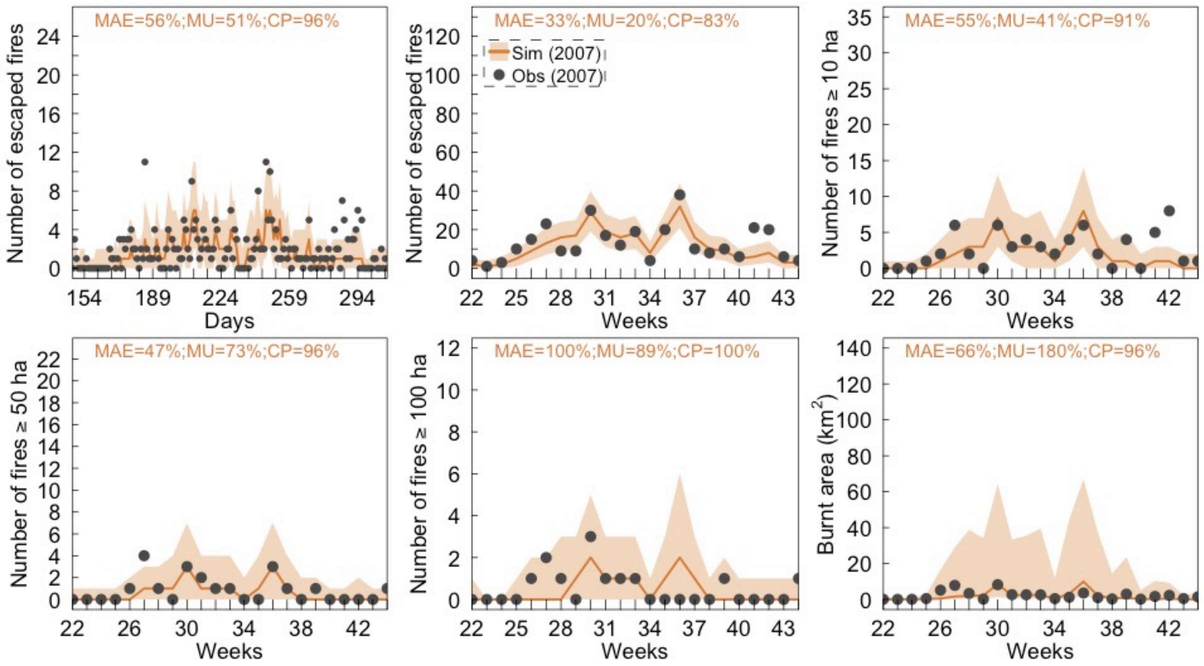
MAE=56%;MU=51%;CP=96%
Number of escaped fires
Days
MAE=33%;MU=20%;CP=83%
Sim (2007)
Obs (2007)
Number of escaped fires
Weeks
MAE=55%;MU=41%;CP=91%
Number of fires ≥ 10 ha
Weeks
MAE=47%;MU=73%;CP=96%
Number of fires ≥ 50 ha
Weeks
MAE=100%;MU=89%;CP=100%
Number of fires ≥ 100 ha
Weeks
MAE=66%;MU=180%;CP=96%
Burnt area (km²)
Weeks

N. 2008

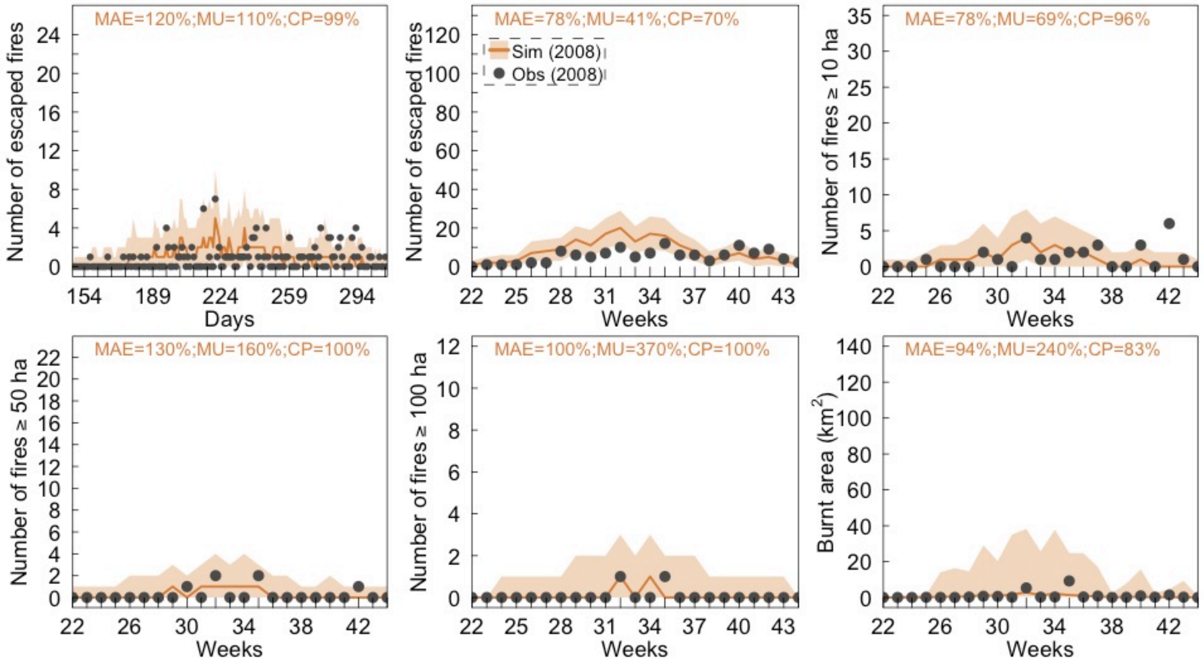
MAE=120%;MU=110%;CP=99%
Number of escaped fires
Days
MAE=78%;MU=41%;CP=70%
Sim (2008)
Obs (2008)
Number of escaped fires
Weeks
MAE=78%;MU=69%;CP=96%
Number of fires ≥ 10 ha
Weeks
MAE=130%;MU=160%;CP=100%
Number of fires ≥ 50 ha
Weeks
MAE=100%;MU=370%;CP=100%
Number of fires ≥ 100 ha
Weeks
MAE=94%;MU=240%;CP=83%
Burnt area (km²)
Weeks

O. 2009

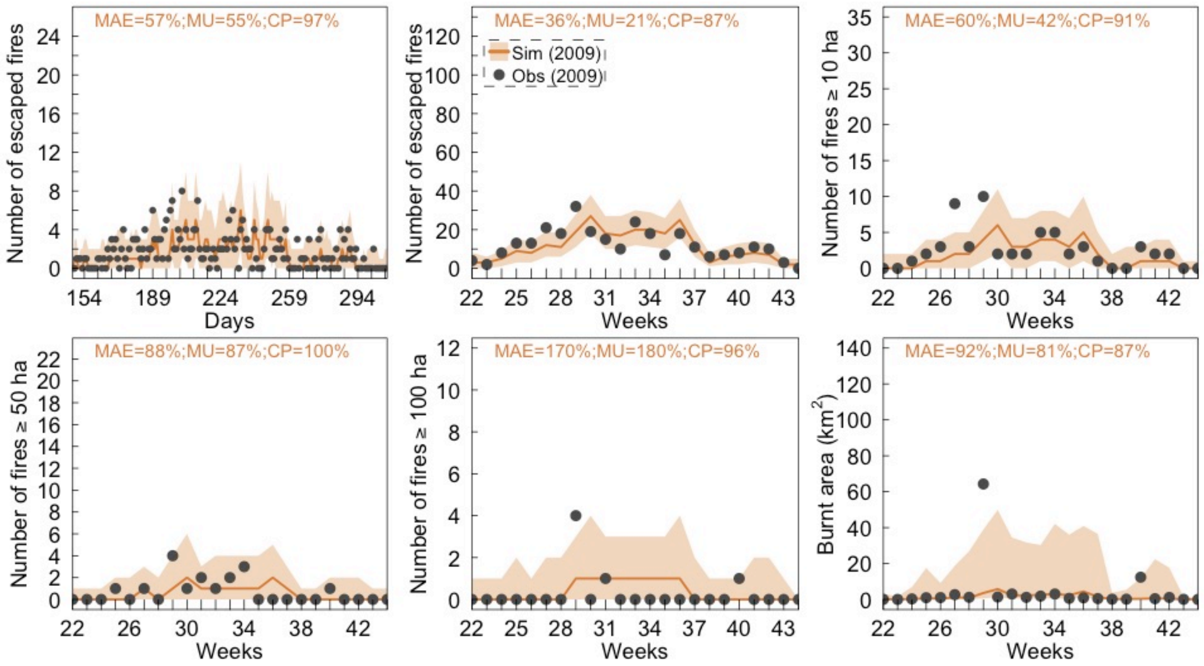
MAE=57%;MU=55%;CP=97%
Number of escaped fires
Days
MAE=36%;MU=21%;CP=87%
Sim (2009)
Obs (2009)
Number of escaped fires
Weeks
MAE=60%;MU=42%;CP=91%
Number of fires ≥ 10 ha
Weeks
MAE=88%;MU=87%;CP=100%
Number of fires ≥ 50 ha
Weeks
MAE=170%;MU=180%;CP=96%
Number of fires ≥ 100 ha
Weeks
MAE=92%;MU=81%;CP=87%
Burnt area (km2)
Weeks

**P. 2010**

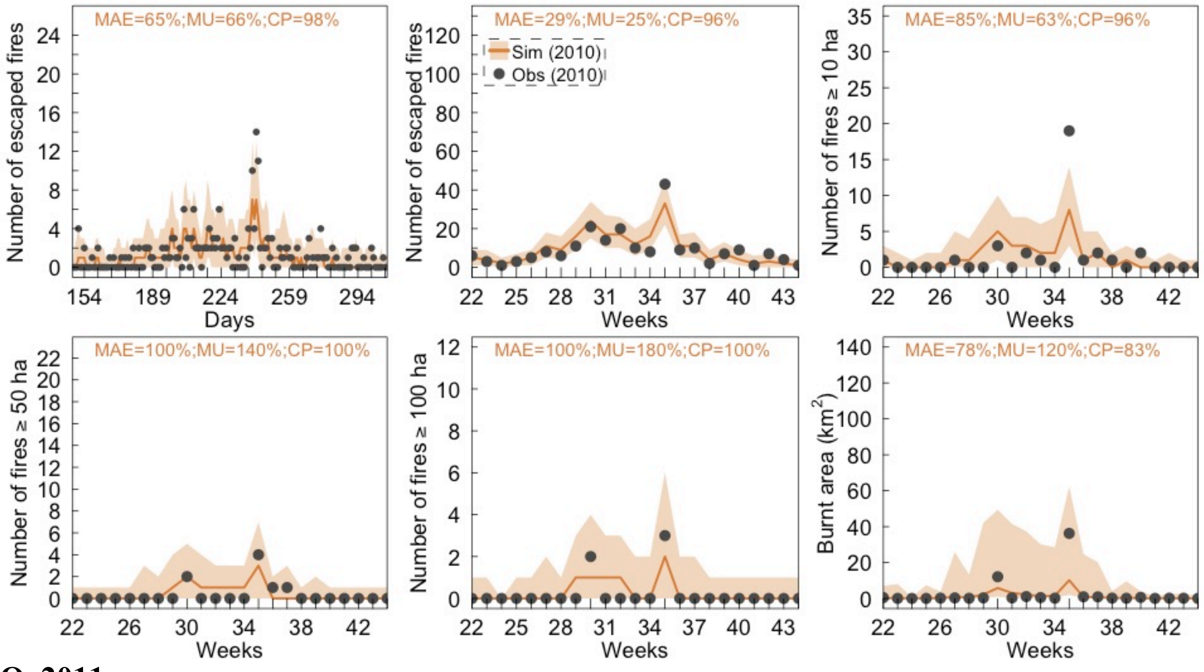
MAE=65%;MU=66%;CP=98%
Number of escaped fires
Days
MAE=29%;MU=25%;CP=96%
Sim (2010)
Obs (2010)
Number of escaped fires
Weeks
MAE=85%;MU=63%;CP=96%
Number of fires ≥ 10 ha
Weeks
MAE=100%;MU=140%;CP=100%
Number of fires ≥ 50 ha
Weeks
MAE=100%;MU=180%;CP=100%
Number of fires ≥ 100 ha
Weeks
MAE=78%;MU=120%;CP=83%
Burnt area (km2)
Weeks

**Q. 2011**

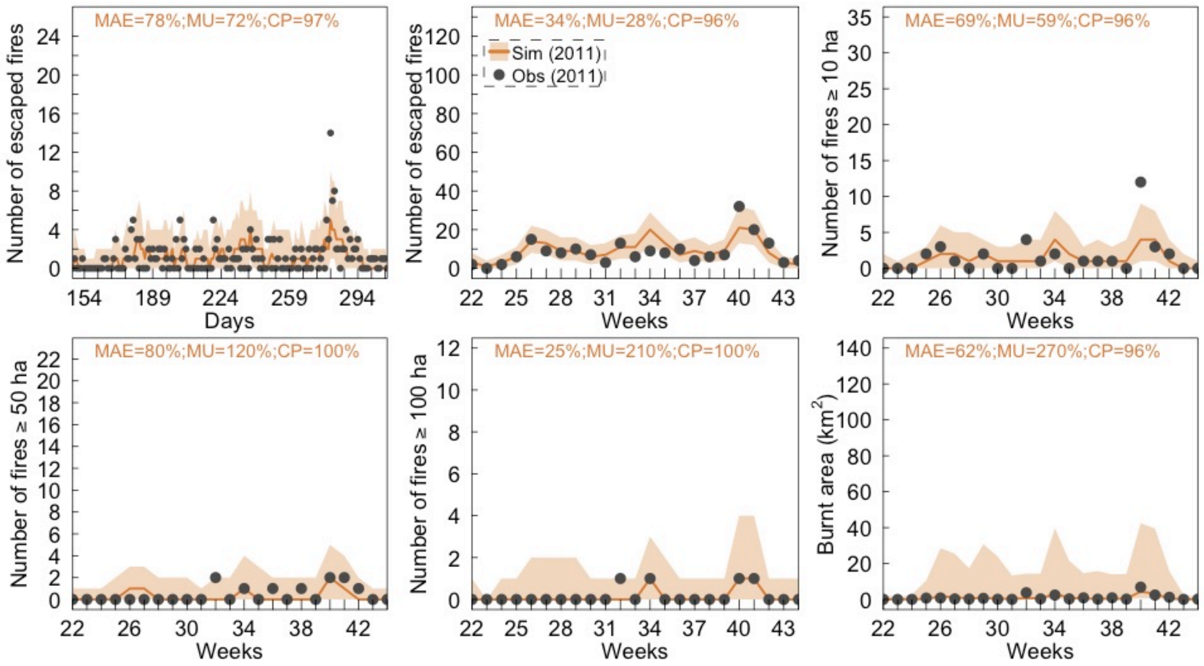
MAE=78%;MU=72%;CP=97%
Number of escaped fires
Days
MAE=34%;MU=28%;CP=96%
Sim (2011)
Obs (2011)
Weeks
MAE=69%;MU=59%;CP=96%
Number of fires ≥ 10 ha
MAE=80%;MU=120%;CP=100%
Number of fires ≥ 50 ha
MAE=25%;MU=210%;CP=100%
Number of fires ≥ 100 ha
MAE=62%;MU=270%;CP=96%
Burnt area (km2)

R. 2012

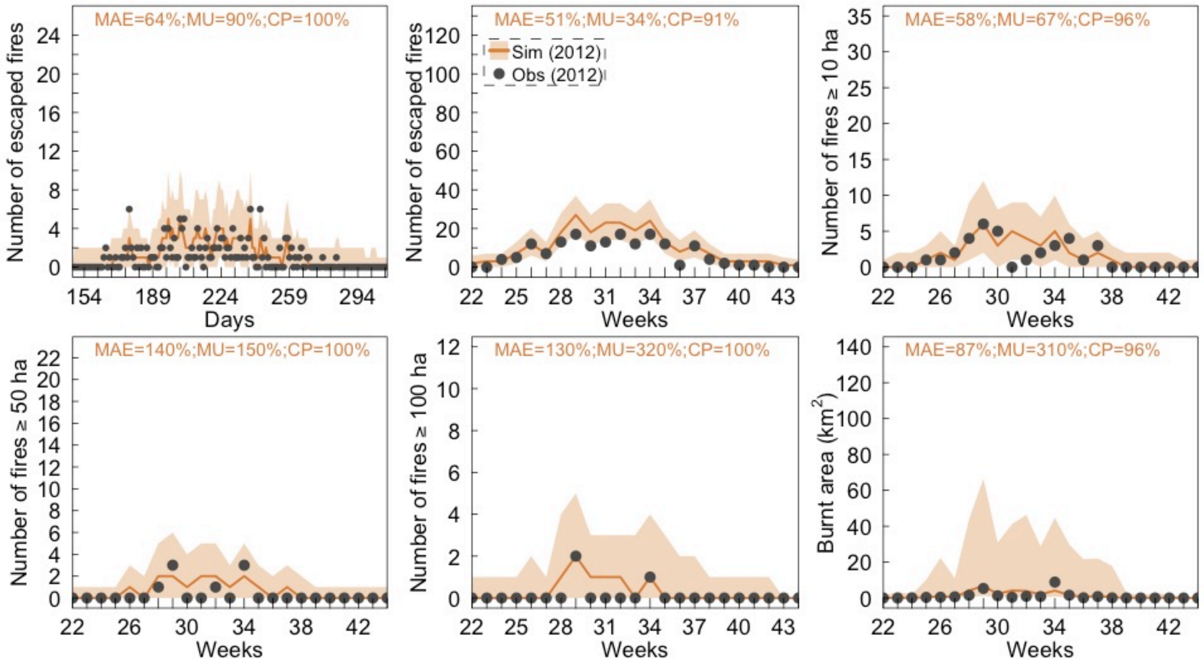
MAE=64%;MU=90%;CP=100%
Number of escaped fires
Days
MAE=51%;MU=34%;CP=91%
Sim (2012)
Obs (2012)
Weeks
MAE=58%;MU=67%;CP=96%
Number of fires ≥ 10 ha
MAE=140%;MU=150%;CP=100%
Number of fires ≥ 50 ha
MAE=130%;MU=320%;CP=100%
Number of fires ≥ 100 ha
MAE=87%;MU=310%;CP=96%
Burnt area (km2)

S. 2013

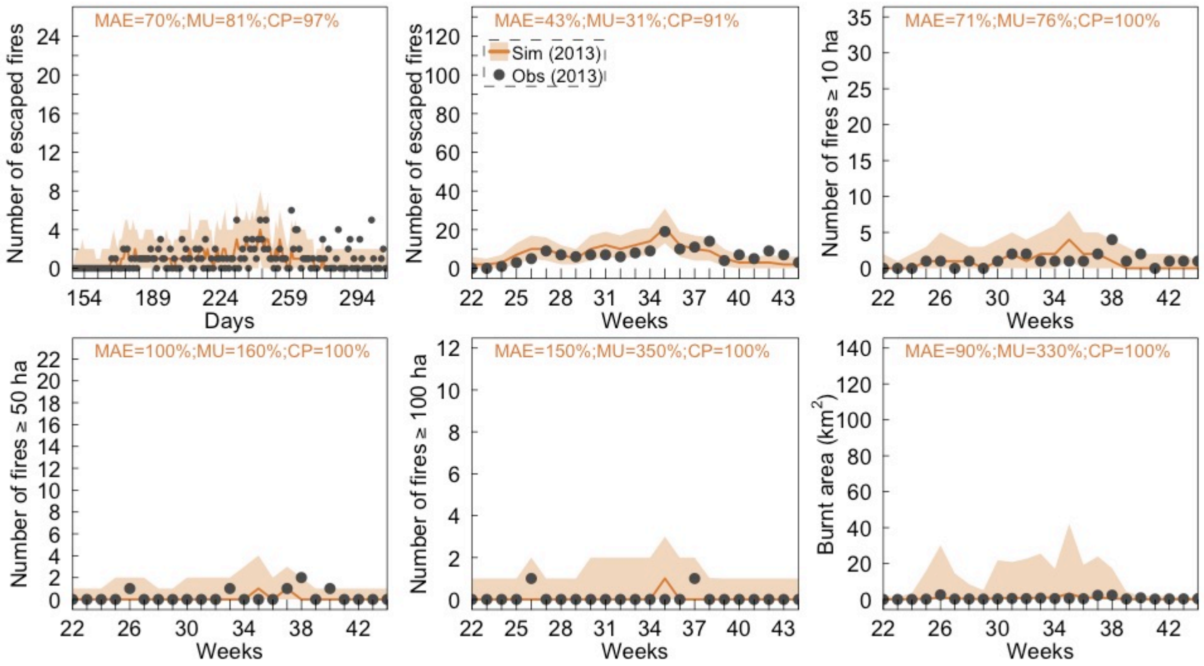
MAE=70%;MU=81%;CP=97%
Number of escaped fires
Days
MAE=43%;MU=31%;CP=91%
Sim (2013)
Obs (2013)
Number of escaped fires
Weeks
MAE=71%;MU=76%;CP=100%
Number of fires ≥ 10 ha
Weeks
MAE=100%;MU=160%;CP=100%
Number of fires ≥ 50 ha
Weeks
MAE=150%;MU=350%;CP=100%
Number of fires ≥ 100 ha
Weeks
MAE=90%;MU=330%;CP=100%
Burnt area (km²)
Weeks

**T. 2014**

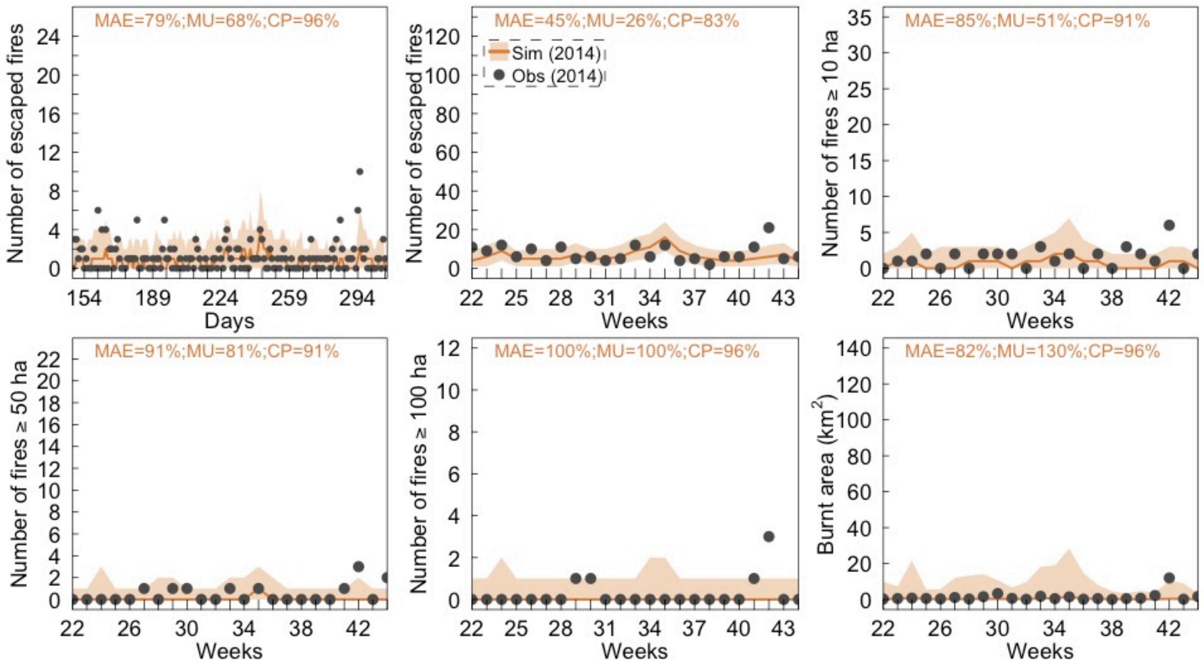
MAE=79%;MU=68%;CP=96%
Number of escaped fires
Days
MAE=45%;MU=26%;CP=83%
Sim (2014)
Obs (2014)
Number of escaped fires
Weeks
MAE=85%;MU=51%;CP=91%
Number of fires ≥ 10 ha
Weeks
MAE=91%;MU=81%;CP=91%
Number of fires ≥ 50 ha
Weeks
MAE=100%;MU=100%;CP=96%
Number of fires ≥ 100 ha
Weeks
MAE=82%;MU=130%;CP=96%
Burnt area (km²)
Weeks

**U. 2015**

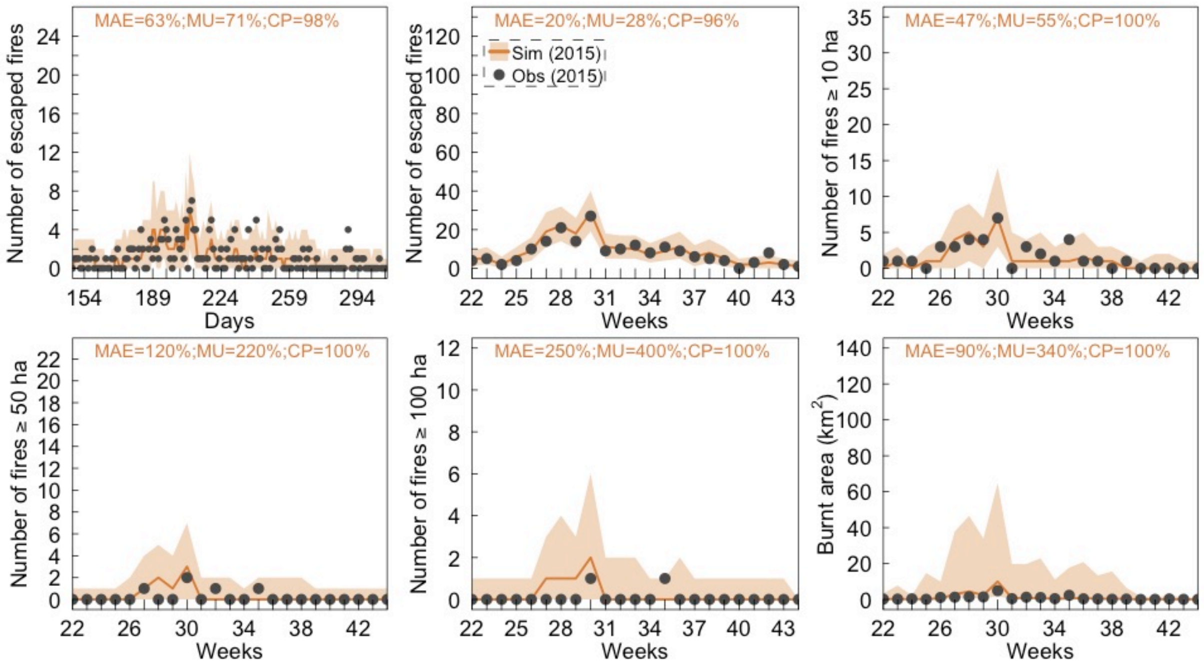
MAE=63%;MU=71%;CP=98%
Number of escaped fires
Days
MAE=20%;MU=28%;CP=96%
Sim (2015)
Obs (2015)
Number of escaped fires
Weeks
MAE=47%;MU=55%;CP=100%
Number of fires ≥ 10 ha
Weeks
MAE=120%;MU=220%;CP=100%
Number of fires ≥ 50 ha
Weeks
MAE=250%;MU=400%;CP=100%
Number of fires ≥ 100 ha
Weeks
MAE=90%;MU=340%;CP=100%
Burnt area (km$^2$)
Weeks

**V. 2016**

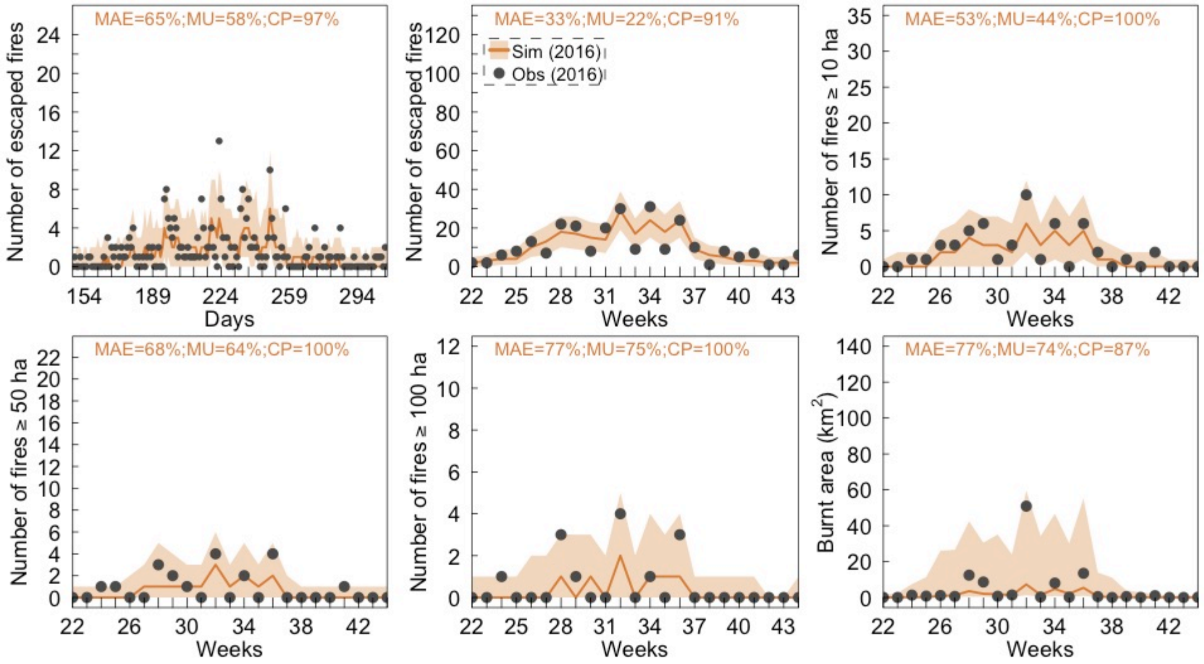
MAE=65%;MU=58%;CP=97%
Number of escaped fires
Days
MAE=33%;MU=22%;CP=91%
Sim (2016)
Obs (2016)
Number of escaped fires
Weeks
MAE=53%;MU=44%;CP=100%
Number of fires ≥ 10 ha
Weeks
MAE=68%;MU=64%;CP=100%
Number of fires ≥ 50 ha
Weeks
MAE=77%;MU=75%;CP=100%
Number of fires ≥ 100 ha
Weeks
MAE=77%;MU=74%;CP=87%
Burnt area (km$^2$)
Weeks

**W. 2017**

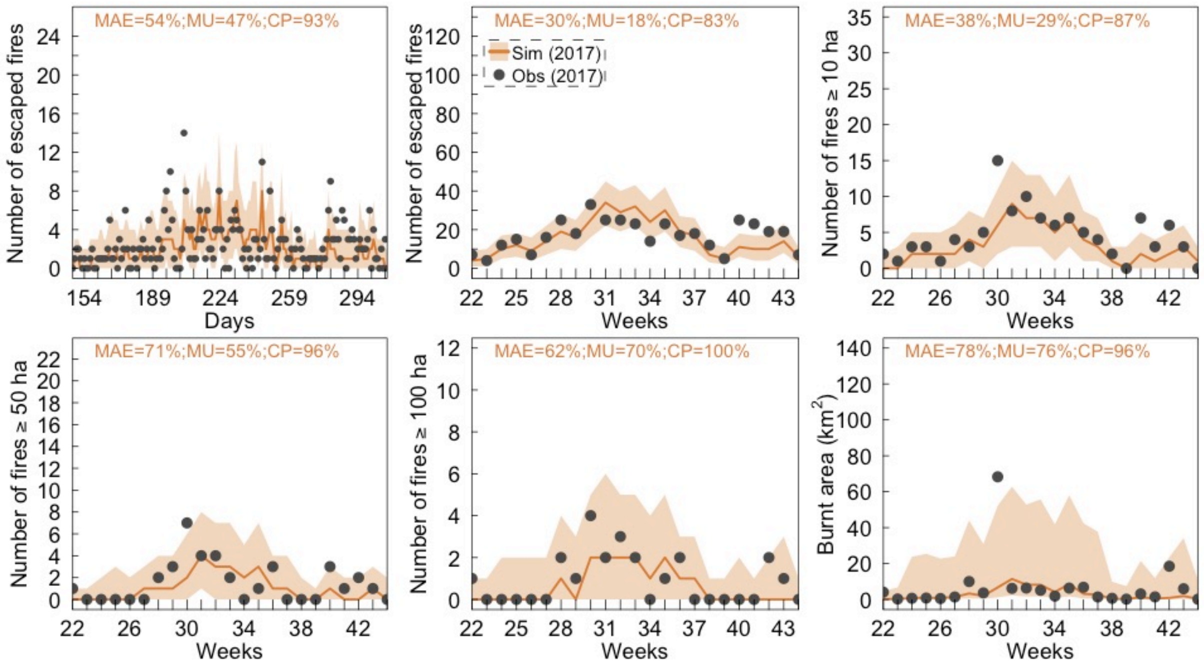
MAE=54%;MU=47%;CP=93%
Number of escaped fires
Days
MAE=30%;MU=18%;CP=83%
Sim (2017)
Obs (2017)
Number of escaped fires
Weeks
MAE=38%;MU=29%;CP=87%
Number of fires ≥ 10 ha
Weeks
MAE=71%;MU=55%;CP=96%
Number of fires ≥ 50 ha
Weeks
MAE=62%;MU=70%;CP=100%
Number of fires ≥ 100 ha
Weeks
MAE=78%;MU=76%;CP=96%
Burnt area (km$^2$)
Weeks

**X. 2018**

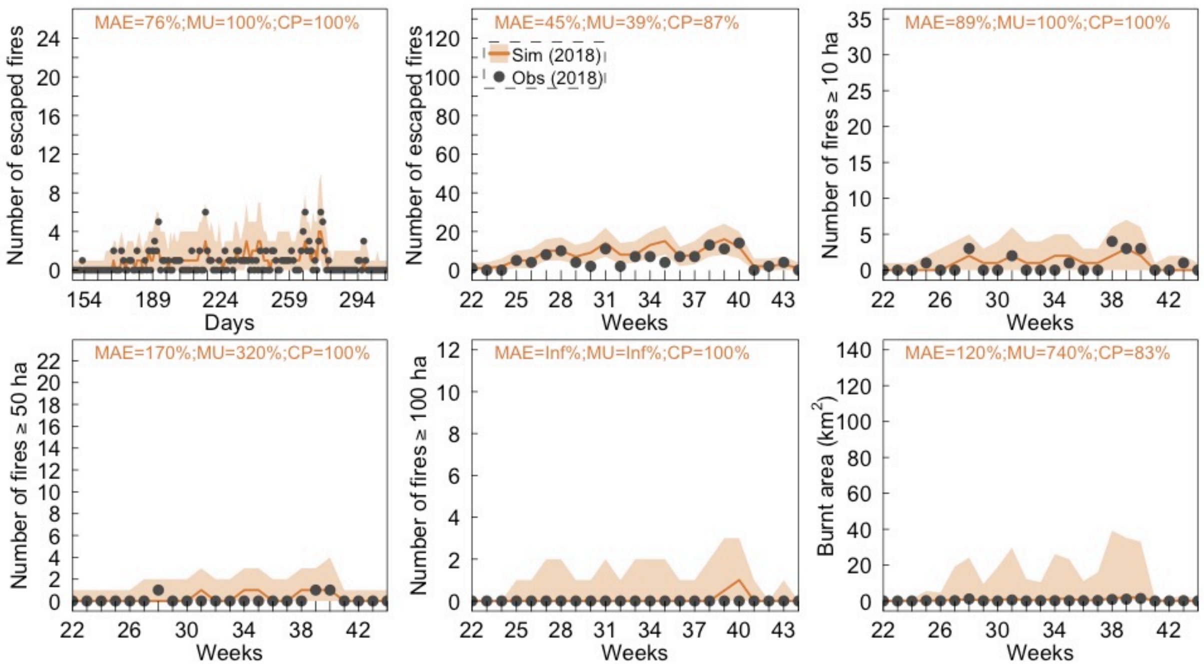
MAE=76%;MU=100%;CP=100%
Number of escaped fires
Days
MAE=45%;MU=39%;CP=87%
Sim (2018)
Obs (2018)
Number of escaped fires
Weeks
MAE=89%;MU=100%;CP=100%
Number of fires ≥ 10 ha
Weeks
MAE=170%;MU=320%;CP=100%
Number of fires ≥ 50 ha
Weeks
MAE=Inf%;MU=Inf%;CP=100%
Number of fires ≥ 100 ha
Weeks
MAE=120%;MU=740%;CP=83%
Burnt area (km$^2$)
Weeks

**Supplementary 5.** Same as FIG. 5 and 11 for other years. Comparison of simulated fire size distribution with observation: Expected trend (red line) was surrounded by the 95th confidence interval in orange and the 99.9th confidence interval (light orange), computed from 1000 simulations of fire sizes for year of interest.

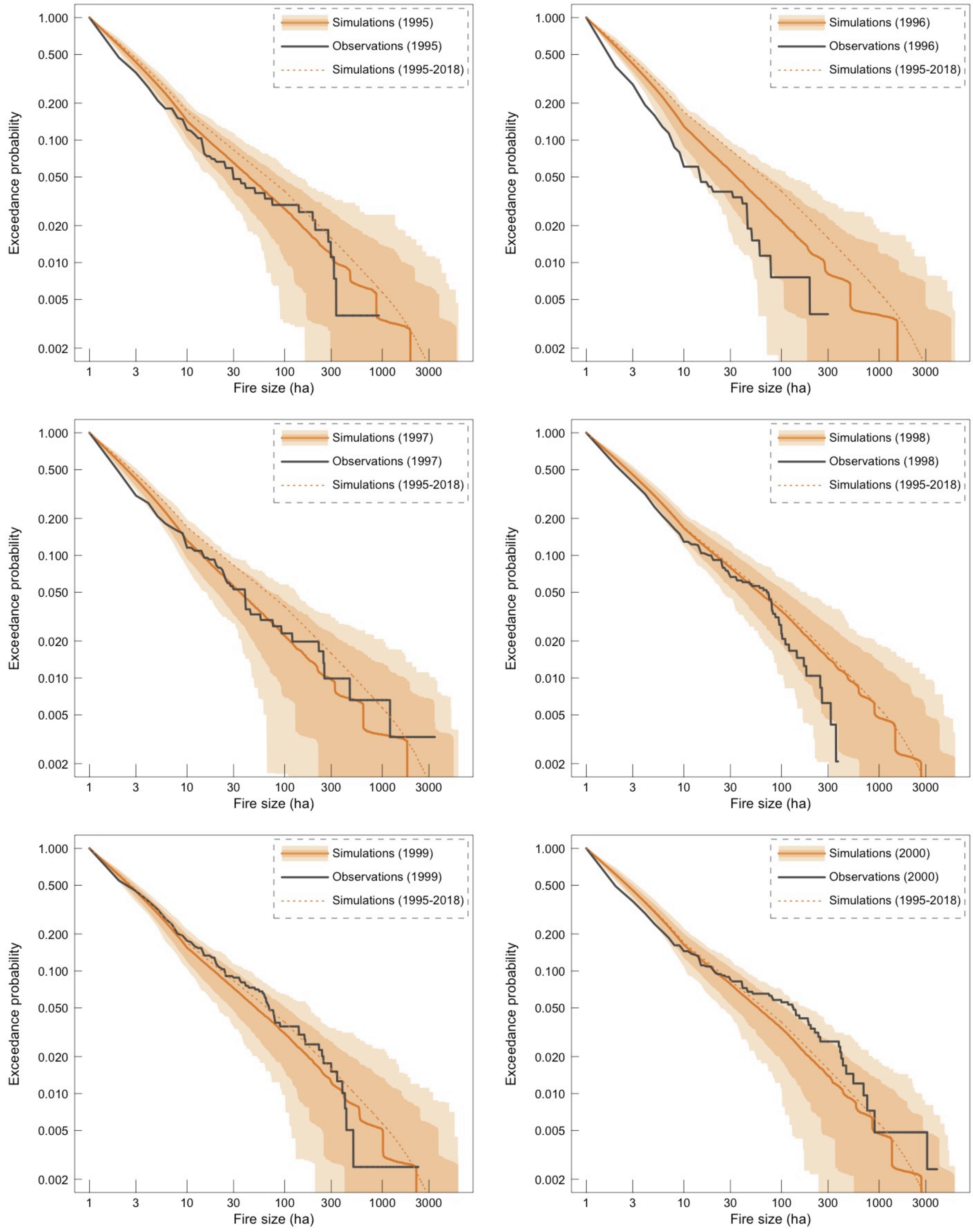

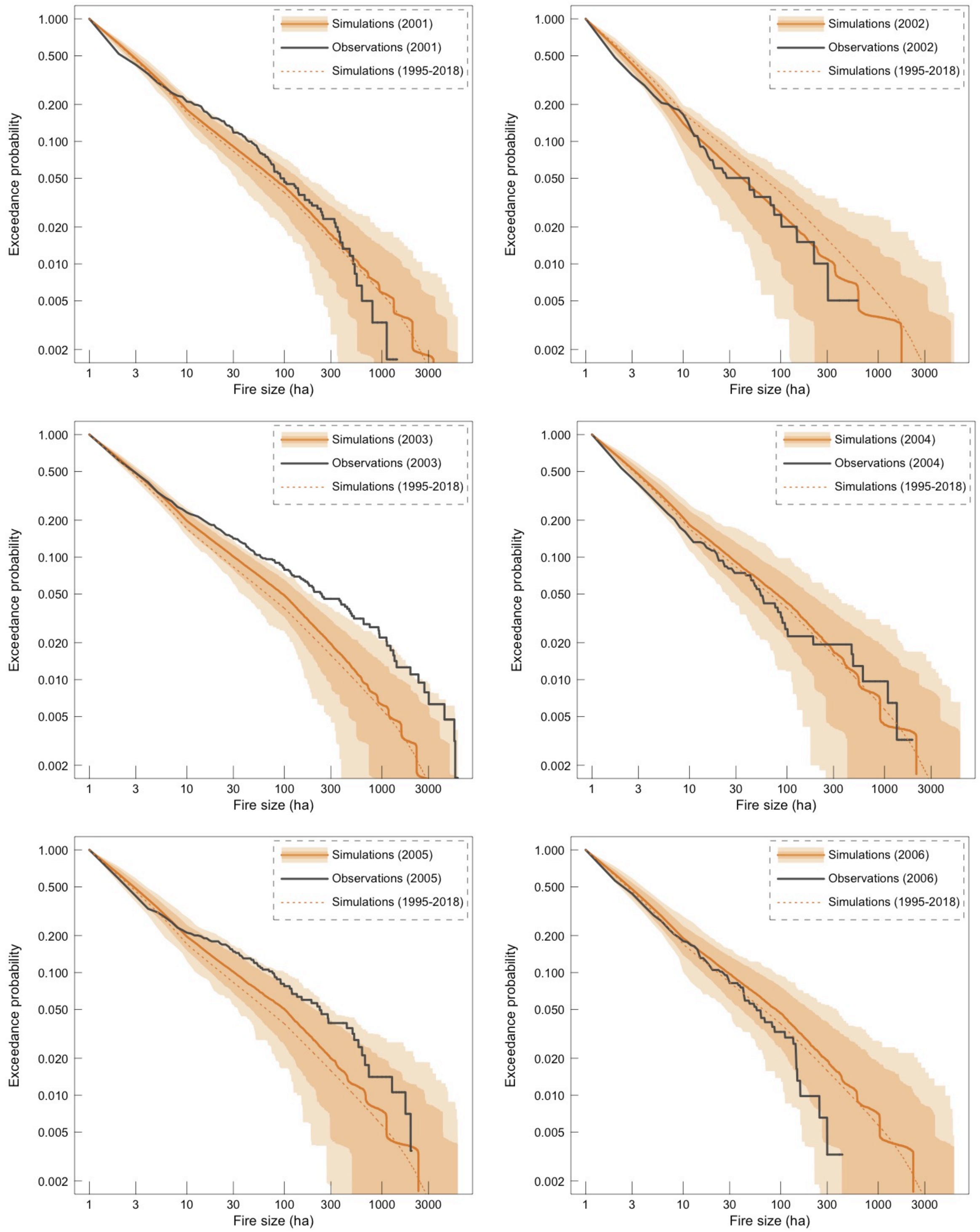

Simulations (2001)
Observations (2001)
Simulations (1995-2018)
Simulations (2002)
Observations (2002)
Simulations (1995-2018)
Simulations (2003)
Observations (2003)
Simulations (1995-2018)
Simulations (2004)
Observations (2004)
Simulations (1995-2018)
Simulations (2005)
Observations (2005)
Simulations (1995-2018)
Simulations (2006)
Observations (2006)
Simulations (1995-2018)
Exceedance probability
1.000
0.500
0.200
0.100
0.050
0.020
0.010
0.005
0.002
Fire size (ha)
1
3
10
30
100
300
1000
3000

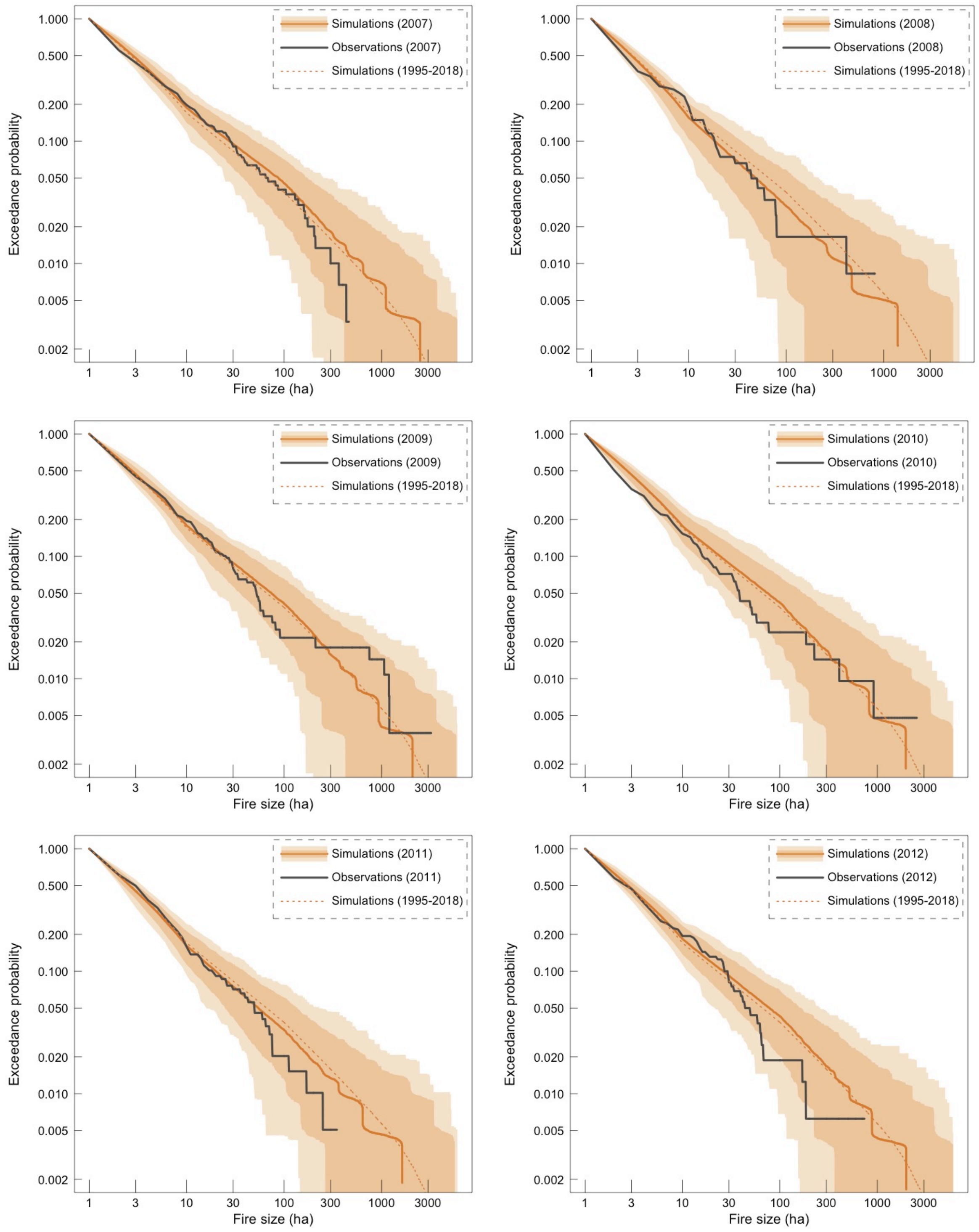

Simulations (2007)
Observations (2007)
Simulations (1995-2018)
Simulations (2008)
Observations (2008)
Simulations (1995-2018)
Simulations (2009)
Observations (2009)
Simulations (1995-2018)
Simulations (2010)
Observations (2010)
Simulations (1995-2018)
Simulations (2011)
Observations (2011)
Simulations (1995-2018)
Simulations (2012)
Observations (2012)
Simulations (1995-2018)
Exceedance probability
1.000
0.500
0.200
0.100
0.050
0.020
0.010
0.005
0.002
Fire size (ha)
1
3
10
30
100
300
1000
3000

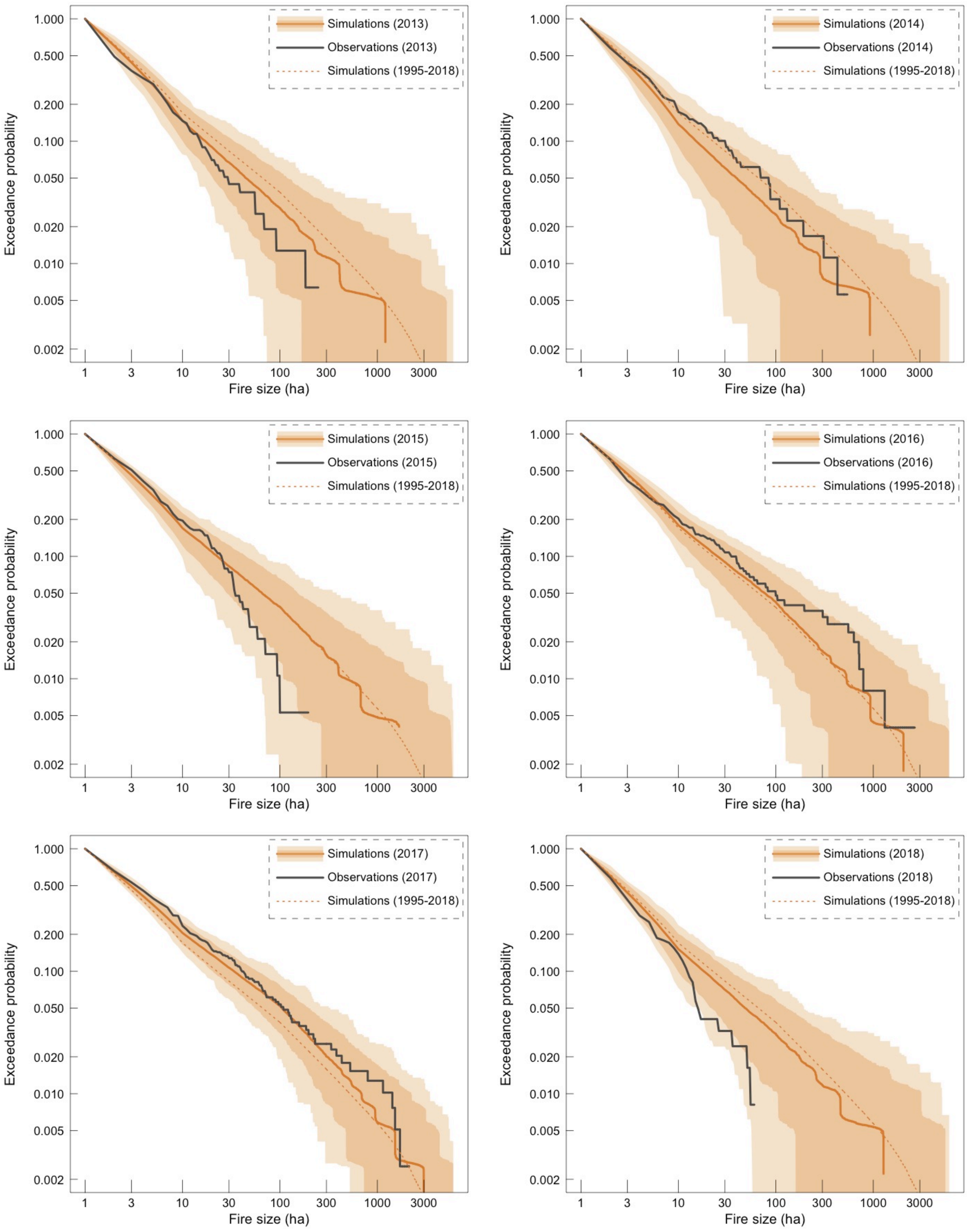
Simulations (2013)
Observations (2013)
Simulations (1995-2018)
Simulations (2014)
Observations (2014)
Simulations (1995-2018)
Simulations (2015)
Observations (2015)
Simulations (1995-2018)
Simulations (2016)
Observations (2016)
Simulations (1995-2018)
Simulations (2017)
Observations (2017)
Simulations (1995-2018)
Simulations (2018)
Observations (2018)
Simulations (1995-2018)
Exceedance probability
1.000
0.500
0.200
0.100
0.050
0.020
0.010
0.005
0.002
Fire size (ha)
1
3
10
30
100
300
1000
3000
